# Hyperexcitability of female serotonin neurons underlies sex-specific anxiety responses

**DOI:** 10.1101/2025.08.17.670715

**Authors:** Lucia Pizzoccaro, Suzanne van der Veldt, Fiona Henderson, Anne-Sophie Simard, Félix Perreault, Justine Fortin-Houde, Elsa Lalot, Guillaume Ducharme, Bénédicte Amilhon

**Affiliations:** CHU Sainte-Justine Azrieli Research Center, Montréal, Québec, Canada; Département de Neurosciences, Université de Montréal, Montréal, Québec, Canada; Groningen Institute for Evolutionary Life Sciences, University of Groningen, Groningen, the Netherlands

## Abstract

Mood and anxiety disorders display robust sex differences in prevalence, symptom profile, and treatment outcomes, yet the circuit mechanisms underlying this sex bias remain unclear. Here, we identify a serotonergic (5-HT) pathway from the median raphe region (MRR) to the ventral hippocampus (vHP) that drives sex-specific anxiety regulation in mice. Using a multimodal approach combining electrophysiology, fiber photometry, and optogenetics, we show that vHP-projecting 5-HT neurons (5-HT^vHP^) in the MRR are intrinsically hyperexcitable in females and exhibit delayed adaptation during exposure to aversive environments. At baseline, female mice displayed greater avoidance and reduced risk-assessment behavior. Optogenetic activation of this pathway selectively enhanced anxiety-like behavior and stress-related grooming in females, while leaving locomotion unaffected. Fiber photometry revealed that grooming episodes coincide with transient suppression of 5-HT^vHP^ activity, suggesting an adaptive feedback mechanism to downregulate serotonergic tone under elevated anxiety. Moreover, activation of this pathway disrupted hippocampal theta dynamics during habituation to a novel arena exclusively in females, revealing serotonergic modulation of anxiety and novelty processing. These results were consistent with the identification of sex-specific M-currents, which constrained excitability in male MRR 5-HT^vHP^ neurons, while being largely absent in females. Collectively, our findings uncover a hyperexcitable MRR-vHP serotonergic circuit that drives female-specific anxiety states, providing a mechanistic framework for understanding sex-specific vulnerability to mood and anxiety disorders.

## Introduction

Mood disorders disproportionately affect women, who are not only more likely to be diagnosed, but also exhibit different symptom profiles and treatment responses compared to men^1,2^. Although this sex bias is well-documented, its neurobiological underpinnings remain poorly understood and largely understudied. Elucidating how biological sex influences mood and vulnerability to mood disorders at the neural circuit level is essential for developing more effective treatment strategies.

Anxiety disorders which are among the most prevalent mood-related conditions, show nearly a twofold higher incidence in women relative to men^2^. Drugs increasing serotonin (5-hydroxytryptamine, 5-HT) transmission remain first-line treatment for anxiety and related mood disorders, demonstrating the central role of 5-HT in the neurobiology of anxiety^3^. Yet, paradoxically, acute manipulations of 5-HT signaling, whether via pharmacology or optogenetic stimulation, have shown that increased serotonergic activity can exacerbate anxiety-like behavior in rodents^4–8^. These conflicting effects underscore the developmental, electrophysiological and functional heterogeneity of the 5-HT system^9,10^, which comprises a diverse population of neurons distributed across multiple raphe nuclei, with widespread, frequently collateralized projections that collectively innervate the entire forebrain^4,11–15^. Moreover, multiple 5-HT receptors, some with opposing action, can be co-expressed in the same brain regions and neurons^16–20^. This complexity has long hampered our understanding of how serotonergic signalling regulates mood states.

Among the many targets of serotonergic cells, the ventral hippocampus (vHP), a core component of the limbic system, stands out as a critical node in the regulation of anxiety and behavioral inhibition^21–24^. The vHP shares connections with the lateral septum, prefrontal cortex (PFC), lateral hypothalamus and raphe, circuits previously described for their role in anxiety processing^25–29^. Lesions of the vHP consistently produce anxiolytic effects^21,23^. vHP pyramidal cells and interneurons represent aversive and potentially threatening environments^25,27,30–33^ and manipulation of vHP activity or outputs modulates anxiety-like behaviors in rodents^25,26,29,33,34^. In humans, reductions in anterior hippocampal volume are observed in individuals with anxiety disorders^35^, and the anterior hippocampus is engaged during approach-avoidance behaviors^36,37^, reinforcing the conserved role of this structure in anxiety regulation across species.

The median raphe region (MRR), comprising the median raphe nucleus (MnR) and the paramedian raphe (PMnR), is the main source of serotonergic inputs to the hippocampus in mice^13,38,39^. Although comparatively less explored than the dorsal raphe nucleus^9,40–43^, growing evidence highlights the MRR as a key group of 5-HT neurons with roles in aversion, avoidance and more^44–50^. While the impacts of 5-HT release on dorsal hippocampus (dHP) activity and functions have been documented^7,20,51–57^, effects of 5-HT on the vHP have received less attention. Altered 5-HT transmission in the vHP enhances anxiety-related behaviors, suggesting a critical pathway for 5-HT-mediated regulation of anxiety states^8,58–60^. The involvement of this pathway, and the extent to which sex-specific differences in vHP-projecting 5-HT neurons (5-HT^vHP^ neurons) from the MRR contribute to anxiety regulation, await clarification.

This study aims to advance understanding of how serotonergic circuits contribute to the sex disparity in mood disorders. We postulate that 5-HT^vHP^ neurons are a central component of the neural circuits governing anxiety. To test this hypothesis, we use a combination of circuit-specific fiber photometry, electrophysiology, optogenetics and behavioral analysis. We show that these neurons exhibit heightened intrinsic excitability, spontaneous activity and increased *in vivo* recruitment in females. We demonstrate that female hyperexcitability results from a striking sex difference in Kv7/M-channels activity, with their mediated M-current selectively constraining the excitability of male MRR 5-HT^vHP^ neurons. These sex differences translate into circuit- and behavior-specific consequences: 5-HT^vHP^ neurons in females exhibit delayed disengagement from aversive contexts, and female mice display greater baseline anxiety-like behavior, including increased avoidance, reduced risk assessment, and increased grooming. In females, 5-HT^vHP^ neurons selectively alter hippocampal theta dynamics during novelty exposure, and promote anxiety-like behavior, increased risk-assessment, and grooming when optogenetically activated. Together, our findings delineate a circuit-specific mechanism by which enhanced serotonergic activity in vHP-projecting neurons contributes to sexually divergent anxiety responses, offering insights in how sex differences at the cellular and circuit level may underlie vulnerability to mood disorders.

## Results

### MRR 5-HT**^vHP^** neurons are more excitable in female mice

Serotonergic neurons display strikingly diverse electrophysiological properties, including a wide range of firing frequencies, action potential waveforms and resting membrane potential. Although investigations of sex-differences in these electrophysiological properties are scarce, a handful of studies have reported differences in excitability of 5-HT neurons^61–64^. To probe the contribution of raphe-vHP circuits in sex-specific anxiety behaviors we examined whether 5-HT^vHP^ neurons differ in their intrinsic electrophysiological properties.

We first mapped the distribution of 5-HT^vHP^ neurons within raphe sub-regions. 5-HT^vHP^ neurons were identified using retrograde expression of eYFP, obtained with injections of a Cre-dependent retrogradely expressed adeno-associated virus in the vHP of SERT-Cre mice (Fig. 1a). Transfected neurons were similarly distributed across multiple raphe subregions for males and females, with labeled neurons primarily located in the B9 cell group, the MRR and the interfascicular part of the DR (DRI; Fig. 1b,c; Extended Data Fig. 1). Importantly, the proportion of 5-HT cells retrogradely labeled from the vHP was similar between males and females, representing in % of 5-HT neurons: 17.91 ± 0.19 and 18.90 ± 1.53 in B9, 14.36 ± 1.32 and 19.24 ± 2.82 in the MRR, and 15.18 ± 0.74 and 20.33 ± 2.94 in the DRI, in females and males respectively (Fig. 2c; Extended Data Fig. 1b-c). We performed whole-cell patch-clamp recordings in acute brain slices from retrogradely labeled eYFP-positive cells in the B9, MRR and DRI from male and female mice (Fig. 1, Extended Data Fig. 1d-e). In the B9 region, no sex differences were observed in firing frequency (Fig. 1d-f), resting membrane potential (RMP, Fig. 1g), input resistance (Fig. 1h), or spike frequency adaptation during sustained activation, quantified by the adaptation index (Fig. 1i, see methods). In contrast, 5-HT^vHP^ neurons in the MRR from female mice exhibited consistently higher firing rates across all depolarizing current steps (Fig. 1j-l) and a more depolarized RMP compared to males (Fig. 1m). Notably, these neurons showed a significantly reduced adaptation index compared to males (Fig. 1o). The absence of differences in input resistance (Fig. 1n) suggests that sex differences in excitability are not due to changes in passive membrane properties. In turn, 5-HT^vHP^ neurons in the DRI showed comparable firing frequency between sexes (Fig. 1p-r) but, similarly to observations in the MRR, their RMP was more depolarized in female mice (Fig. 1s). No differences between sexes were found for DRI 5-HT^vHP^ neurons input resistance (Fig. 1t) or adaptation index (Fig. 1u). Notably, the firing frequencies of female MRR 5-HT^vHP^ neurons were the highest among all regions and sexes (Extended Data Fig. 1f). Moreover, under baseline conditions, 29% of female MRR 5-HT^vHP^ neurons (5/17 cells) exhibited spontaneous firing, which was not observed in any neuron recorded from the B9 and DRI regions in either sex, or in male MRR 5-HT^vHP^ neurons.

**Figure 1:**
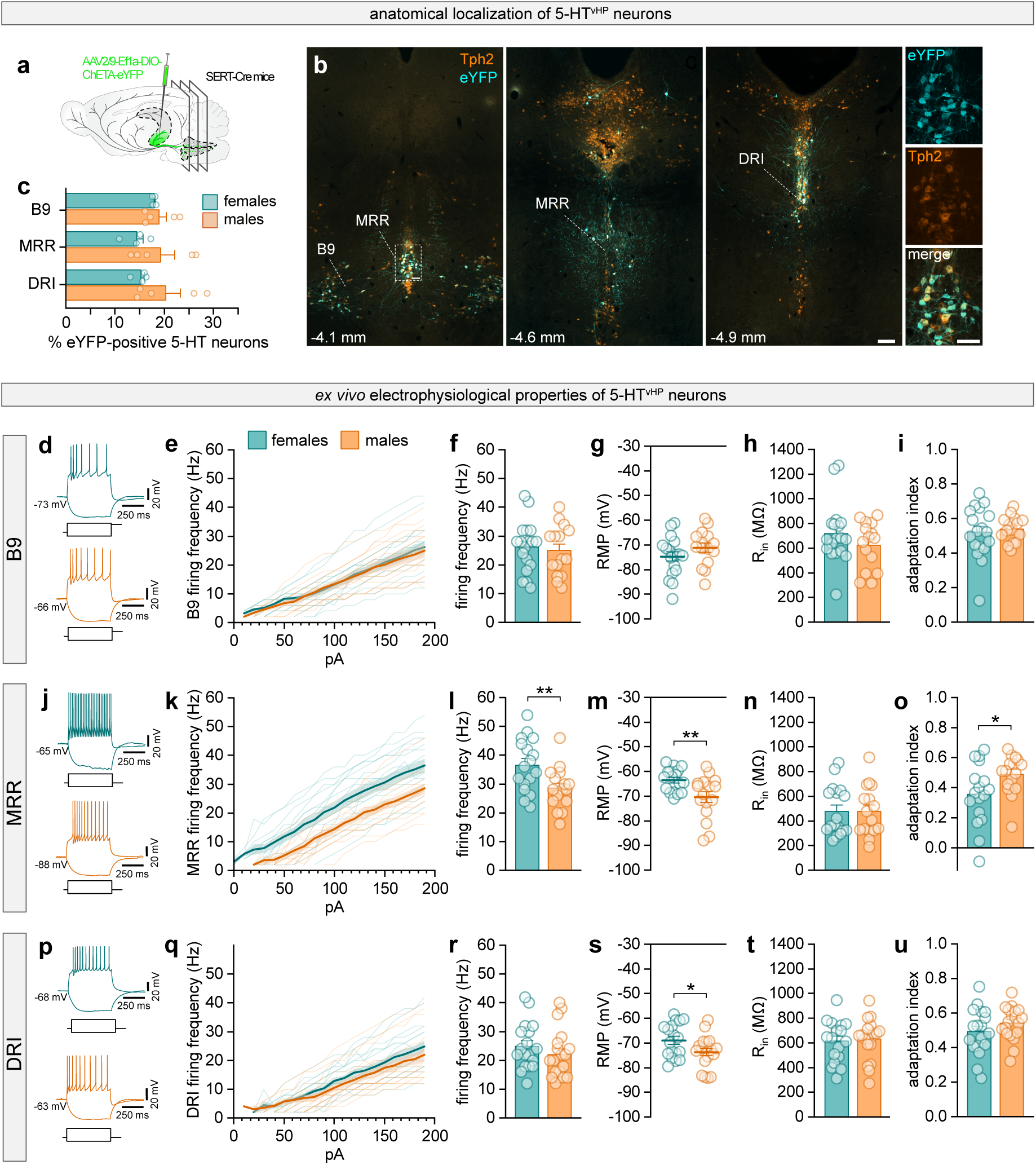
Female MRR 5-HT^vHP^ neurons exhibit increased intrinsic excitability. **a,** Schematic of experimental configuration: a retrograde AAV containing Cre-dependent construct eYFP was bilaterally infused into the vHP of SERT-Cre mice. **b,** Representative coronal sections showing eYFP-labeled neurons in the B9, MRR and DRI regions at different anteroposterior levels (distance from bregma in mm indicated in bottom left corner). 5-HT+ cells are detected using Tph2 immunostaining. Scale bars = 200 μm (overviews), 100 μm (magnified). **c,** Percentage of eYFP+ 5-HT neurons in raphe of females (teal, n = 4 mice) and males (orange, n = 4 mice). **d,** Examples of membrane potential responses to current steps (-50 pA for lower trace and +190 pA for upper trace) as part of the electrophysiological characterization of eYFP+ neurons recorded from female (teal) and male (orange) animals for 5-HT^vHP^, from the B9 nucleus. Number on the left indicates resting membrane potential. **e – i,** For B9 5-HT^vHP^ neurons, firing frequency responses as a function of current intensity stimulations (**e**), firing frequency at +190 pA (**f**), resting membrane potential (**g**), input resistance (**h**) and adaptation index (**i**) for females (teal, n = 18 cells from 9 mice) and males (orange, n = 15 cells from 10 mice). **j,** Same as **d,** for 5-HT^vHP^ neurons recorded from the MRR. **k – o,** Same as **e-i**, for eYFP+ neurons recorded from the MRR. **l,** two sample t-test, *P* = 0.0089, **m,** two sample t-test, *P* = 0.0079, and **o,** two sample t-test, *P* = 0.0357, for females (teal, n = 17 cells from 6 mice) and males (orange, n = 16 cells from 6 mice). **p**, Same as **d** and **j**, for eYFP+ neurons recorded from DRI. **q – u,** Same as for **e**-**i** and **k**-**o**, for eYFP+ neurons recorded from DRI. **s,** two sample t-test, *P* = 0.0207, for females (teal, n = 18 cells from 9 mice) and males (orange, n = 19 cells from 10 mice). All data are shown as mean ± s.e.m. *: *P* < 0.05; **: *P* < 0.01. For detailed statistical information, see Extended Data Table 1. B9: B9 nucleus; MRR: median raphe region; DRI: dorsal raphe, interfascicular subregion; eYFP: enhanced yellow fluorescent protein; Tph2: tryptophan hydroxylase 2.

**Figure 2:**
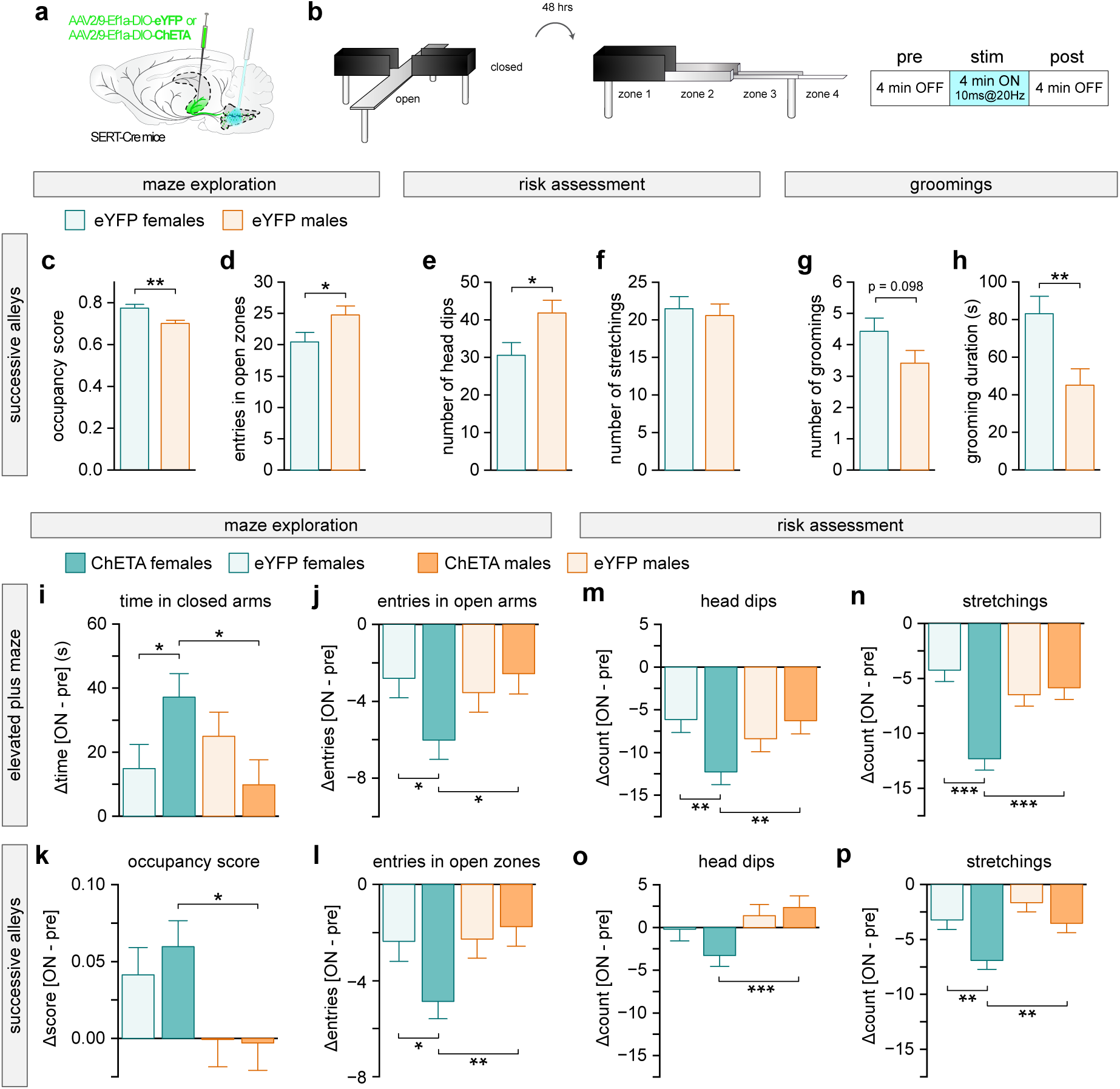
5-HT^vHP^ activation increases anxiety-like behaviors and decreases risk assessment selectively in females. **a,** Schematic of experimental configuration: a retrograde AAV containing Cre-dependent excitatory opsin ChETA or eYFP control is bilaterally infused in the vHP of SERT-Cre mice of both sexes and an optic fiber is implanted above the MRR region. **b,** Experimental set up: schematic of EPM (left) and SA (middle) apparatus with optogenetic stimulation paradigm (right). **c-h**, Comparison of male versus female behaviors for eYFP control mice, during 12 minutes of SA exploration, analyzed by two sample t-test. Significant comparisons are reported as *P* values. Occupancy score, *P* = 0.0034 (**c**), entries in open zones, *P* = 0.0499 (**d**), number of head dips, *P* = 0.0249 (e), number of stretchings (f), number of groomings (g), and grooming duration, *P* = 0.0199 (h) for female (light teal, n = 20), and male (light orange, n = 20). **i**-**p**, Change (stim – pre) in behavioral measures during EPM and SA testing. Data are from two independent experiments and analyzed by three-way ANOVA with Holm’s multiple comparison testing. **i**, Time in closed arms of the EPM: females (ChETA vs control), *P* = 0.0378; ChETA (female vs male), *P* = 0.0126. **j,** Open arm entries of the EPM: females (ChETA vs control), *P* = 0.0278; ChETA (female vs male). *P* = 0.0204. **k,** Occupancy score (see methods) in the SA: ChETA (female vs male), *P* = 0.0134. **l,** Number of entries into open zone (zone 2+3+4) : females (ChETA vs control), *P* = 0.0269; ChETA (female vs male), *P* = 0.0059. **m,** Number of head dips in the EPM: females (ChETA vs control), *P* = 0.0052; ChETA (female vs male), *P* = 0.0068. **n,** Number of stretchings in the EPM: females (ChETA vs control), *P* < 0.0001; ChETA (female vs male), *P* < 0.0001. **o,** Number of head dips in the SA: ChETA (female vs male), *P* = 0.0009. **p**, Number of stretchings in the SA: females (ChETA vs control), *P* = 0.0026; ChETA (female vs male), *P* = 0.0039. Data are shown as mean ± s.e.m. *: *P* < 0.05, **: *P* < 0.01; ***: *P* < 0.001. For detailed statistical information, see Extended Data Table 1. For an overview of all behavioral variables analyzed, see Extended Data Table 2.

To determine whether these differences in excitability reflect a general property of female MRR serotonergic neurons or are specific to the vHP-projecting population, we performed additional recordings from non-vHP projecting MRR 5-HT neurons (MRR 5-HT^non-vHP^, Extended Data Fig. 1g-p). Using a dual-labeling strategy, MRR 5-HT neurons were locally labeled with mCherry, while MRR 5-HT^vHP^ neurons were retrogradely labeled from the vHP with eYFP (Extended Data Fig. 1g-j). Expression of mCherry was highly specific to serotonergic neurons, with 96.47 ± 0.40 % of mCherry+ neurons also expressing Tph2. In contrast to the vHP-projecting population, MRR 5-HT^non-vHP^ neurons did not exhibit sex differences in firing frequency (Extended Data Fig. 1k-m), resting membrane potential (Extended Data Fig. 1n), input resistance (Extended Data Fig. 1o) or adaptation index (Extended Data Fig. 1p).

Together, these findings demonstrate that while 5-HT^vHP^ neurons are similarly distributed in male and female mice, they display subregion- and sex-specific differences in intrinsic properties. Whereas B9 and DRI 5-HT^vHP^ neurons were largely similar between the sexes, female MRR 5-HT^vHP^ neurons are uniquely hyperexcitable, exhibiting more depolarized resting potentials, higher firing rates, reduced spike frequency adaptation and spontaneous activity.

### Optogenetic activation of 5-HT^vHP^ neurons promotes anxiety-like behavior and decreases risk-assessment in females

The MnR is the primary source of 5-HT inputs to the vHP in mice^13^ and has previously been implicated in the regulation of anxiety behavior^44,65,66^. As females show higher anxiety than males across species^1,2,67^, we hypothesized that increased excitability of MRR 5-HT^vHP^ female neurons might contribute to sex differences in the behavioral response to aversive contexts and potential threat. We used the elevated plus maze (EPM), a classical anxiety assay, and the successive alleys (SA) task, that both reliably elicits approach-avoidance conflict behaviors in male and female mice^68,69^, and tested whether stimulating 5-HT^vHP^ neurons would alter anxiety-like and risk assessment behaviors.

To assess the functional consequences of selective activation of 5-HT^vHP^ neurons, we expressed an excitatory opsin, ChETA, or a control eYFP construct (Fig. 2a). ChETA expression was highly specific to 5-HT neurons, with only 0.21 ± 0.11% of ChETA-expressing neurons lacking Tph2 co-expression. Light was delivered to the raphe during subsequent exposures to the EPM and SA tasks (Fig. 2b, Extended Data Fig. 2). Two cohorts of male and female mice were tested by experimenters of opposite sex; cohort, sex and stimulation were included as factors in the analysis (Extended Data Table 1, see methods).

We first analyzed eYFP-expressing control mice to determine whether males and females differed in baseline measures of anxiety-like behavior (Fig. 2c-h). While behavior was largely similar in the EPM (Extended Data Fig. 3a-h), females exhibited decreased exploration of the SA, as reflected by higher occupancy scores (Fig 2c, see methods) and less entries in open zones (Fig. 2d). If reduced exploration of SA open areas reflects heightened anxiety, rather than a general decrease in exploratory drive, it should be accompanied by a reduction in risk assessment behaviors. Head dipping, a behavior in which the animal lowers its head over the edge of an open arm, was decreased in females while stretch-attend postures, where mice elongate their bodies in a low stance to cautiously assess their environment, was similar between sexes (Fig. 2e-f). Increased anxiety-like behavior in females in the SA task was also accompanied by an increase in the total time spent grooming over the 12 minutes of test (Fig. 2g-h). No differences were found in total distance travelled or mean velocity between the sexes (Extended Data Fig. 3g-j). Taken together, these metrics are indicative of higher baseline anxiety levels in female mice compared to males.

We next assessed the effects of 5-HT^vHP^ neuron optogenetic activation. In the EPM, activation of 5-HT^vHP^ neurons increased time spent in the closed arms in female ChETA mice as compared to both female eYFP controls and male ChETA mice (Fig. 2i, Extended Data Fig 3k-m). Likewise, during activation female ChETA mice made fewer open arm entries than both same-sex eYFP controls and male ChETA mice (Fig. 2j), consistent with increased anxiety-like behavior. These sex-specific effects were also evident in the SA task. Activation of 5-HT^vHP^ neurons reduced exploration of aversive zones in female ChETA mice, as reflected by higher occupancy scores (Fig. 2k) and fewer entries in the open alleys (Fig. 2l), compared to both female eYFP controls and male ChETA mice. Importantly, these behavioral changes occurred largely independent of alterations in gross locomotor activity. In the EPM, optogenetic stimulation did not significantly affect distance traveled (Extended Data Fig. 3q). In the SA task, however, stimulation of 5-HT^vHP^ neurons reduced distance traveled in female ChETA animals compared to male ChETA mice (Extended Data Fig. 3r), consistent with increased avoidance.

We also analyzed how optogenetic activation of 5-HT^vHP^ neurons impacted risk-assessment behaviors. Head dipping was decreased upon stimulation of 5-HT^vHP^ neurons in female ChETA mice in the EPM (Fig. 2m). Stretch-attend postures were also strongly reduced in female ChETA animals in the EPM (Fig. 2n). This pattern was replicated in the SA, where optogenetic activation similarly reduced both head dipping (Fig. 2o) and stretch-attend postures (Fig. 2p) in female ChETA mice compared to eYFP controls. Together, these findings indicate that activation of 5-HT^vHP^ neurons enhances anxiety-like states in females, suppressing risk-assessment behaviors characteristic of approach-avoidance conflict, rather than merely reducing overall exploratory drive.

Because serotonergic axons form extensive collaterals^14^, we asked whether off-target effects via collateral projections could contribute to the observed behavioral changes. Using eYFP labeling of 5-HT^vHP^ neurons, we mapped their axonal distribution and confirmed collateral innervation across several regions, including the dHP, lateral and medial septum and prefrontal cortex (Extended Data Fig. 4). We observed no sex differences in the pattern or density of these collateral projections, suggesting that sex-specific behavioral effects are unlikely to arise from differential collateralization between the sexes. In addition, to determine whether 5-HT innervation of the vHP itself differs between sexes, we quantified endogenous 5-HT axons density in the vHP of females and males across vHP subregions and layers. We found no sex difference in the surface occupied by 5-HT axons (Extended Data Fig. 5), indicating that the observed behavioral effects are unlikely to reflect gross sex differences in serotonergic terminal density within the vHP.

Collectively, these results suggest that heightened intrinsic excitability of female MRR 5-HT^vHP^ neurons translates into increased baseline anxiety-like behaviors in females and an increased behavioral response to optogenetic activation.

### Female MRR 5-HT^vHP^ neuron exhibit greater recruitment during optogenetic activation

We next asked whether intrinsic electrophysiological differences translate into heightened neuronal activation of female MRR 5-HT^vHP^ neurons during behavior. To address this, we quantified c-fos expression as a proxy for *in vivo* activity following exposure to the SA paradigm combined with 20 Hz optogenetic stimulation. Mice were perfused 70 minutes post-test to capture peak c-fos induction. Immunohistochemical analysis revealed c-fos expression in eYFP-labeled neurons across the B9, MRR and DRI (Fig. 3a). Quantification of the percentage of eYFP+ neurons co-expressing c-fos revealed a significant main effect of optogenetic stimulation across all three subregions in both sexes (Fig. 3b). In the B9 and DRI, optogenetic stimulation robustly increased the number of c-fos+ neurons in both males and females to a similar extent (Fig. 3b). In contrast, in the MRR, a two-way ANOVA revealed a significant main effect of treatment and a significant sex x treatment interaction, consistent with enhanced excitability of MRR 5-HT neurons in females (Fig. 3b). To quantify this differential increase in c-fos, we computed the stimulation-induced change from eYFP-controls in each sex, expressed as [(female ChETA - female eYFP) - (male ChETA - male eYFP)] (Fig. 3c). The magnitude of increase relative to eYFP controls was similar between sexes in the B9 and DRI, while in the MRR, ChETA-expressing females showed a larger recruitment of 5-HT^vHP^ neurons compared to males. These findings indicate that while optogenetic activation of 5-HT^vHP^ neurons reliably drives neuronal activity across raphe subregions, MRR activation was greater in females, mirroring the observed sex-specific behavioral effects and increased intrinsic excitability of female MRR 5-HT^vHP^ neurons.

**Figure 3:**
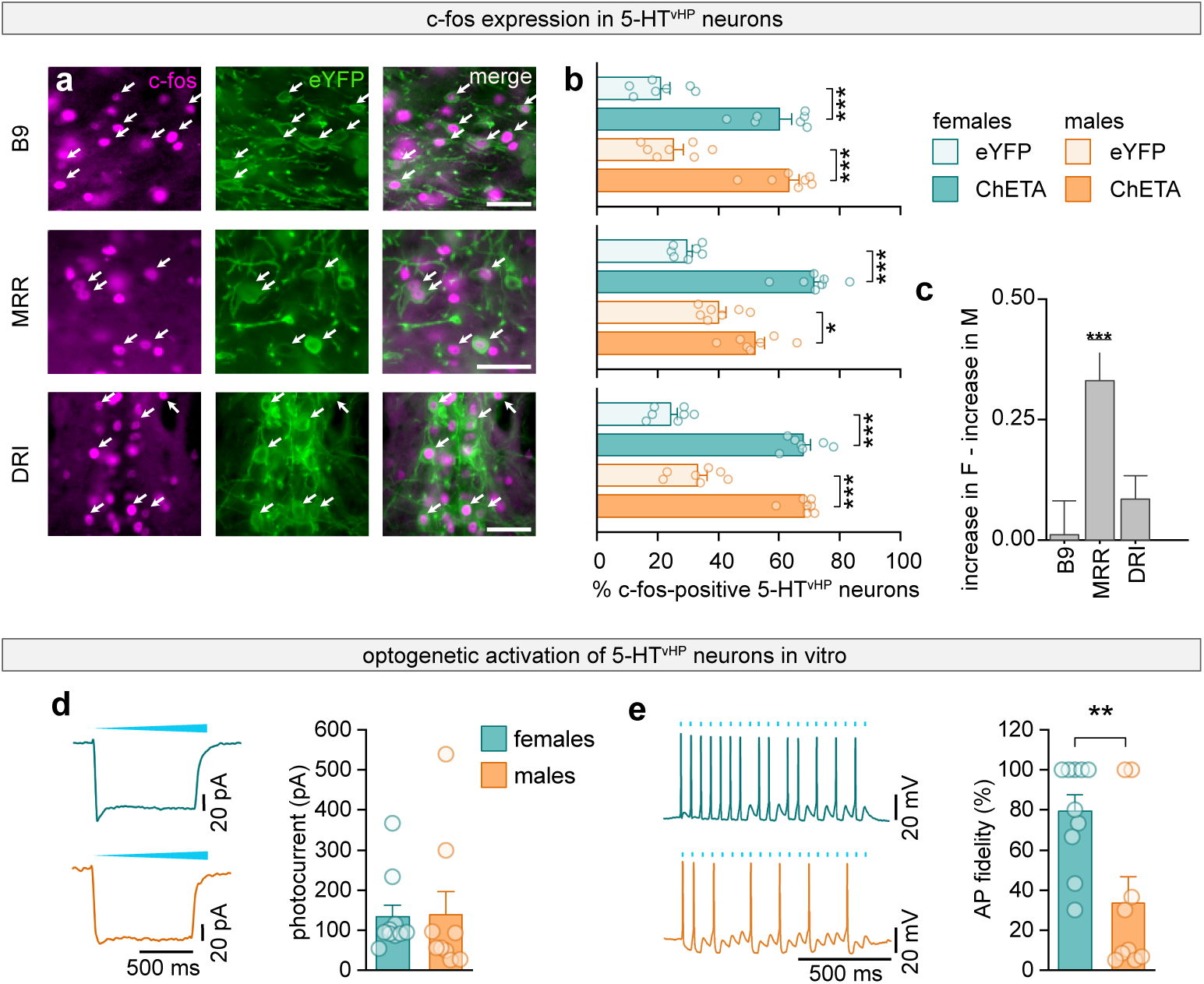
Optogenetic stimulation preferentially recruits female MRR 5-HT^vHP^ neurons. **a,** Representative labeling of cFos (magenta) and retrograde ChETA-eYFP (green) expression in coronal sections of the B9, MRR and DRI. Arrows indicate examples of c-fos+/eYFP+ neurons. Scale bars = 50 μm. **b,** Percentage of eYFP+ neurons co-expressing c-fos in females (teal, eYFP: n = 7; ChETA: n = 7) and males (orange, eYFP: n = 7; ChETA: n = 7). Top: B9; two-way ANOVA with two-sample t-tests: females (ChETA vs control), *P* < 0.0001; males (ChETA vs control): *P* < 0.0001. Middle: MRR; two-way ANOVA with two-sample t-tests: females (ChETA vs control): *P* < 0.0001; males (ChETA vs control): *P* = 0.0114. Bottom: DRI; two-way ANOVA with two-sample t-tests, females (ChETA vs control), *P* < 0.0001; males (ChETA vs control: *P* < 0.0001. **c**, Stimulation induced change (ChETA – eYFP) for females relative to males in the number of c-fos+/eYFP+ neurons for each subregion. One sample t-test, MRR: P < 0.001. **d**, Left: Representative light-evoked photocurrents from ChETA expressing MRR 5-HT^vHP^ neurons in female (teal, top) and male (orange, bottom) mice. Right: Quantification of steady-state photocurrent amplitude for females and males. Two-sample t-test, females vs. males P = 0.9515. **e**, Left: Representative current-clamp recordings from female (teal) and male (orange) ChETA expressing neurons during 20 Hz optical stimulation. Blue ticks indicate individual 10-ms light pulses. Right: Action potential fidelity during 20 Hz optical stimulation, calculated as percentage of light pulses that elicited an action potential, for neurons recorded in females (teal) and males (orange). Two-sample t-test females vs. males *P* = 0.0076. **d-e**, Females: n = 10 cells from 6 mice; males: n = 9 cells from 5 mice. Data are shown as mean ± s.e.m. *: *P* < 0.05, **: *P* < 0.01, ***: *P* < 0.001; For detailed statistical information, see Extended Data Table 1.

To further investigate sex differences in the response of 5-HT^vHP^ neurons to optogenetic activation, we performed additional *ex vivo* recordings from ChETA expressing 5-HT^vHP^ neurons (Fig. 3d-e). Photostimulation evoked comparable photocurrent amplitudes in male and female neurons, demonstrating similar opsin expression and light-evoked current induction across sexes (Fig. 3d). However, during 20 Hz optical stimulation, female neurons exhibited significantly higher action potential fidelity than male neurons (Fig. 3e), indicating a more efficient recruitment of female MRR 5-HT^vHP^ neurons by equivalent photoactivation.

Taken together, these results support the idea that hyperexcitability of the female MRR 5-HT^vHP^ system drives their heightened recruitment during behavior, as revealed by c-fos mapping. *Ex vivo* recordings further suggest that MRR 5-HT^vHP^ female neuron hyperexcitability is driven by differences in intrinsic neuronal properties. Sex-specific recruitment of MRR 5-HT^vHP^ neurons in potentially threatening contexts provides a circuit mechanism underlying differences between males and females in anxiety-like behaviors.

### Activation of 5-HT**^vHP^** neurons promotes anxiety-related grooming in females

Self-grooming is increasingly recognized as a displacement behavior indicative of elevated stress and conflict, as well as a translationally relevant readout of affective states^70–72^. Under conditions of heightened arousal, grooming can serve as an adaptive coping strategy^73^. We tested whether optogenetic activation of 5-HT^vHP^ neurons was sufficient to alter grooming behavior across different environments. During EPM and SA exploration, both anxiogenic contexts, optogenetic activation produced a robust increase in time spent grooming selectively in female ChETA mice relative to female eYFP controls and male ChETA mice (Fig. 4a-b; Extended Data Fig. 3e-g), as well as number of grooming episodes and total grooming duration (Extended Data Table 2). We quantified grooming during exploration of an open field with low-threat characteristics: a small open field in low lighting conditions (39 cm length, 70 ± 5 lux), designed to minimize aversive features such as bright illumination and open space. In contrast to EPM and SA, stimulation of 5-HT^vHP^ in the open field did not significantly affect grooming duration in either sex (Fig. 4c, Extended Data Table 2). These findings indicate that vHP-projecting serotonergic neurons contribute to the expression of stress-related grooming in a sex-and context-dependent manner. The absence of effects in the open field suggests that elevated environmental threat is necessary for this circuit to drive displacement-like behaviors.

**Figure 4:**
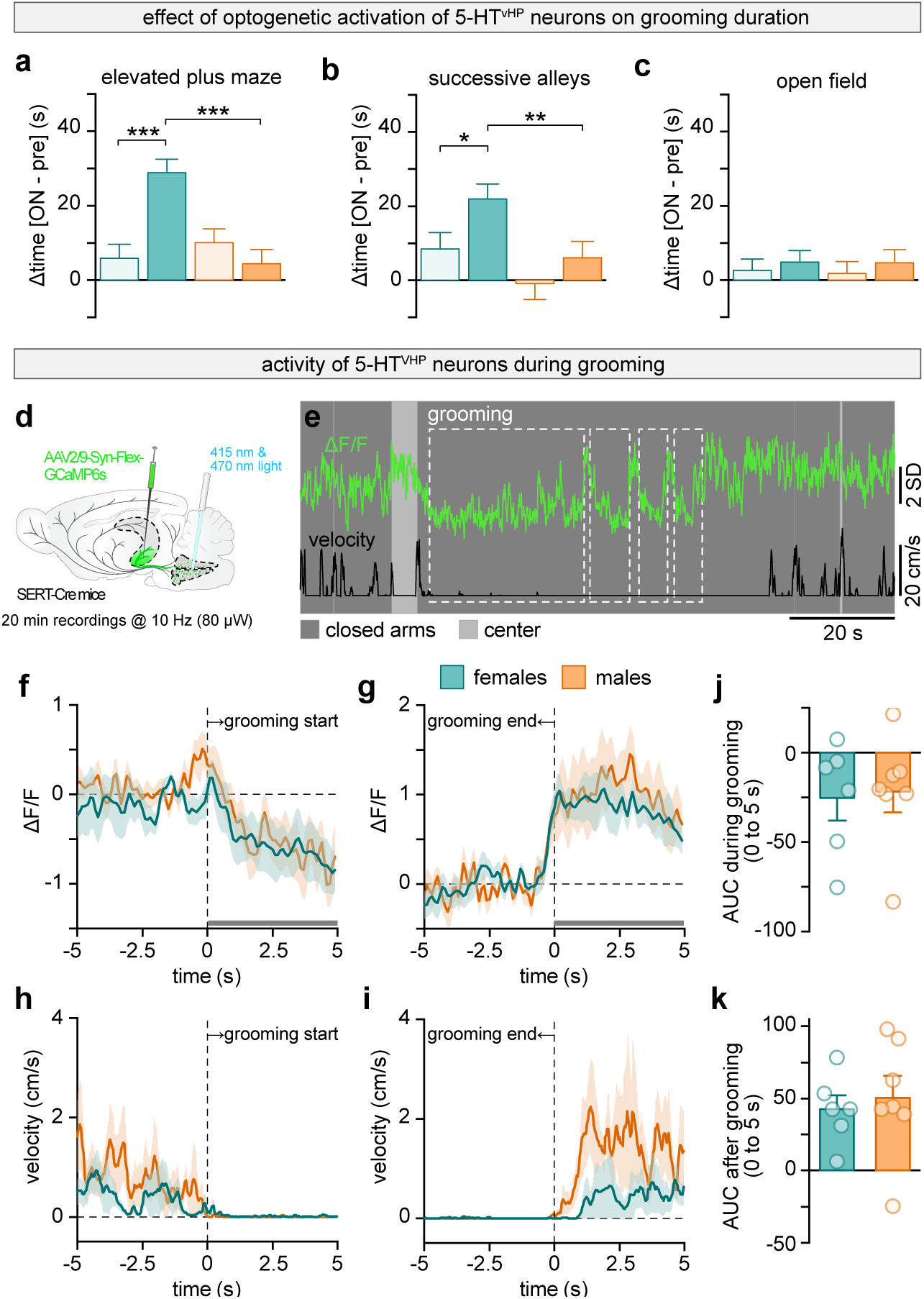
Activation of 5-HT^vHP^ neurons increases grooming in a context-dependent manner. **a,** Change in total grooming time (stim – pre) in the EPM. Three-way ANOVA with Holms multiple comparison testing: females (ChETA vs control), *P* = < 0.0001; ChETA (female vs male), *P* < 0.0001. **b,** Same as **a**, but in the SA. Three-way ANOVA with Holms multiple comparison testing: females (ChETA vs control), *P* = 0.0316; ChETA (female vs male), *P* = 0.0098. **c,** Same as **a**, but in the open field. For **a**-**c**, data are from two independent experiments and were analyzed by three-way ANOVA. **d,** Schematic of experimental configuration: a retrograde AAV containing the Cre-dependent calcium indicator GCaMP6s was infused bilaterally in the vHP of SERT-Cre mice. 5-HT^vHP^ population activity was recorded using an optic fiber implanted above the MRR. **e,** Example 5-HT^vHP^ Ca^2+^ ΔF/F (green) over time, with locomotion velocity (black) during EPM exploration. **f – g,** ΔF/F aligned to grooming onset (**f**) and end (**g**) as indicated by vertical dotted line, for females (teal, n = 6) and males (orange, n = 7). **h – i,** Locomotor velocity for the same animals aligned to start (**h**) and (**i**) end of grooming bouts. **j,** Quantification of the AUC for 0 - 5 s of panel **f** after grooming onset. **k,** Quantification of the AUC for 0 - 5 s of panel **g** after grooming offset. Data are shown as mean ± s.e.m. *: *P* < 0.05, **: *P* < 0.01; For detailed statistical information, see Extended Data Table 1. For an overview of all behavioral variables analyzed, see Extended Data Table 2.

Activation of 5-HT^vHP^ neurons increases anxiety levels and promotes stress-induced grooming, presumably through increased recruitment of hyperexcitable MRR 5-HT^vHP^ neurons in female mice. To monitor activity-related calcium dynamics from this projection-defined population, we used *in vivo* fiber photometry in freely moving SERT-Cre mice. A retrogradely transported AAV encoding the calcium (Ca^2+^) indicator GCaMP6s was bilaterally infused in the vHP, enabling selective monitoring of Ca^2+^ dynamics in MRR 5-HT^vHP^ neurons (Fig. 4d). Fiber placement and viral expression were histologically confirmed to target MRR (Extended Data Fig. 6a-b). GCaMP6s expression was highly specific to 5-HT neurons, with 99.86% of GCaMP6s-expressing cells in the raphe co-expressing the selective 5-HT neuron marker tryptophan hydroxylase 2 (Tph2, n = 2 mice, 709 neurons, data not shown). To assess 5-HT^vHP^ neuronal dynamics related to grooming under anxiogenic conditions, we tested mice in the EPM and aligned population activity to onset and end of grooming bouts. Activity of 5-HT^vHP^ closely tracked grooming behavior (Fig. 4e): grooming episodes coincided with an immediate reduction in calcium signal in both sexes (Fig. 4f). Population activity returned upon grooming termination (Fig. 4g), consistent with the idea that grooming acts as a strategy to transiently downregulate heightened serotonergic activity. Quantification of event-aligned fluorescence revealed no significant sex differences in activity dynamics, either during the first 5 seconds of grooming initiation (Fig. 4j) or in the 5-second periods following termination (Fig. 4k). Grooming bouts were predominantly surrounded by low locomotor activity (<5 cm/s) in both males and females (Fig. 4h-i), indicating that increased 5-HT^vHP^ activity following grooming likely reflects the resumption of exploration. This suggests that grooming may serve as an alternative strategy to downregulate heightened serotonergic activity, consistent with its role as a displacement behavior under conditions of conflict or arousal.

### 5-HTvHP neurons in the MRR display sex-specific activity dynamics in the elevated plus maze

The MRR has been implicated in the response to aversive stimuli^46,47,74^. Notably, the MRR receives direct input from the lateral habenula (LHb), a core structure in aversion-related circuits that signals negative motivational value and modulates serotonergic systems^48,75–77^. Prior work has shown that EPM exploration globally increases 5-HT levels in the vHP, as measured over slow time scales with microdialysis^58,78^. We therefore hypothesized that exploration of a novel and potentially threatening environment such as the EPM would engage 5-HT^vHP^ neurons.

Using the same fiber photometry approach described above, we recorded activity-related calcium dynamics from the somas of retrogradely labelled 5-HT^vHP^ neurons in the MRR during EPM exploration (Fig. 5). We tracked mouse movements and identified locomotor epochs corresponding to transitions from closed arms (“safer” zones) to open arms (aversive zones) of the EPM (Fig. 5a), or transitions restricted to the closed arms (Fig. 5c). Closed-to-open epochs were defined as the mouse spending minimally 10 s in a closed arm before exploring center/open arms for at least 2 s. Closed-to- closed epochs were defined as transitions between closed arms with a maximum of 1 s spent in the center. Changes in fluorescence were averaged and aligned to the timepoint where center-of-body marker crossed the closed arm-center limit. In both sexes and trajectories, 5-HT^vHP^ population activity peaked around transitions (Fig. 5a,c). We observed similar transient increases in 5-HT^vHP^ population activity when mice initiated high velocity locomotion bouts in the EPM, independent of location in the maze, or in the open field (OF) (Extended Data Fig. 6c-j). Velocity and area under the curve (AUC) of fluorescence signal during 1 s surrounding the locomotion onset did not differ between sexes (Extended Data Fig. 6d,f,h,j). Consistent with previous evidence^47,79,80^, these results indicate that 5-HT^vHP^ neurons are engaged during locomotion. Transitions from closed arms into the aversive open compartments were associated with a ramping increase of 5-HT^vHP^ population activity in both sexes (Fig. 5a), suggesting progressive recruitment of this population prior to open-arm entry. By contrast, 5-HT^vHP^ activity remained stable before transitions between closed arms (Fig. 5c).

**Figure 5:**
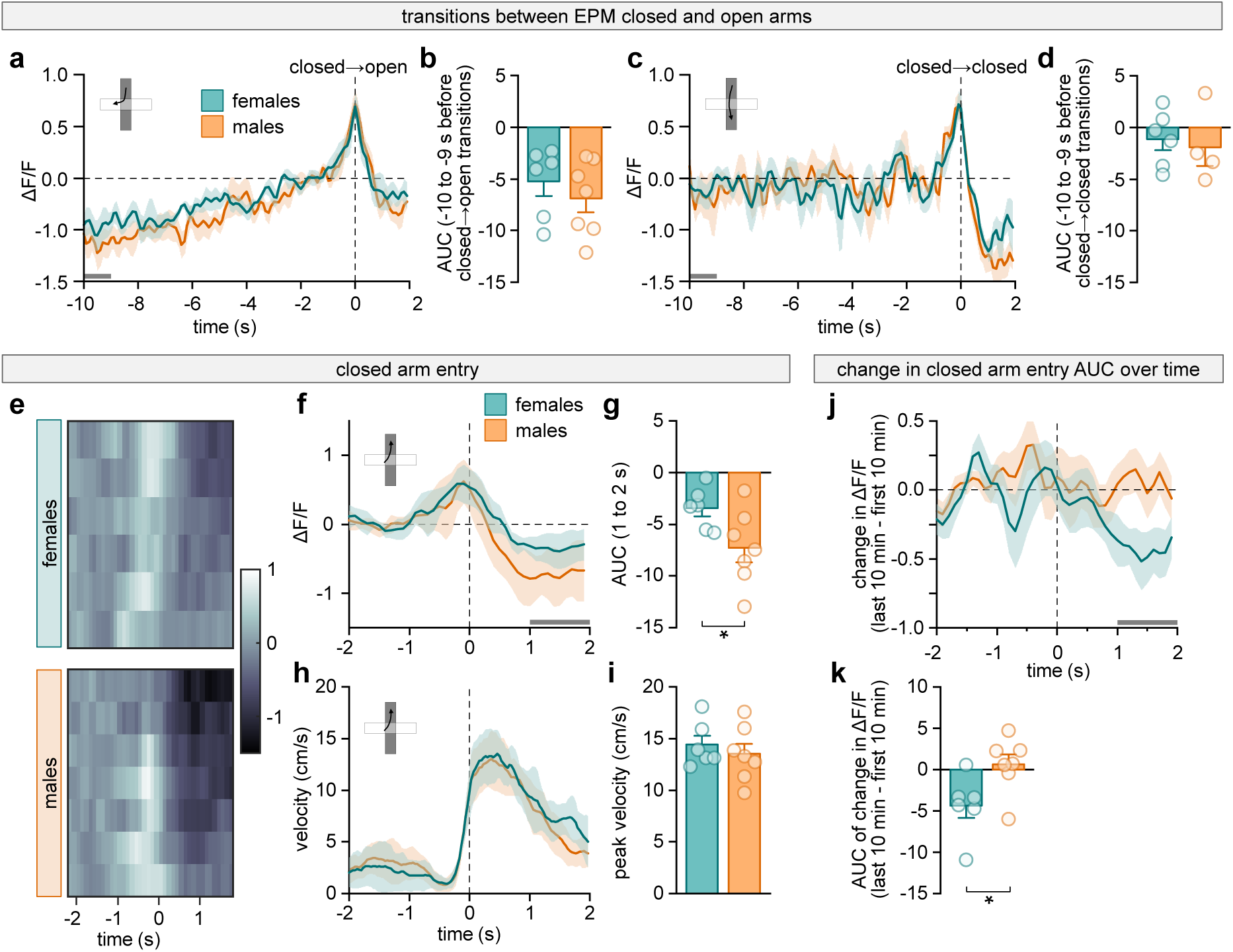
Female 5-HT^vHP^ neurons exhibit delayed disengagement from an aversive context. **a,** Mean normalized ΔF/F aligned to transitions from a closed arm to an open arm for females (teal, n = 6) and males (orange, n= 7). Dotted vertical line indicates timepoint where center-of-body marker crossed the center-closed arm limit. Gray bar indicates the time window used for panel **b**. **b,** Area under the curve (AUC) of ΔF/F obtained over 10 - 9 s preceding closed arm exit (gray bar in **a**). **c,** Same as **a**, but for closed→closed arm transitions. Dotted vertical line indicates timepoint where center-of-body marker crossed the closed-arm center limit exiting the first closed arm. Gray bar indicates the time window used for panel **d**. **d,** AUC of ΔF/F obtained over 10 - 9 s preceding transition timepoint (gray bar in **c**). **e,** Mean ΔF/F for closed arm entries from center/open zones for each female (top, n = 6) and each male (bottom, n = 7). **f,** Mean normalized ΔF/F for closed arm entries from center/open zones for females (teal, n = 6) and males (orange, n = 7). Dotted vertical line indicates timepoint where center-of-body marker crossed the center-closed arm limit. Gray bar indicates the period used for panel **g**. **g,** AUC of ΔF/F obtained over 1 - 2 s after closed arm entry (gray bar in **f**). Two-sample t-test, *P* = 0.0430. **h,** Velocity around closed arm entries for females (teal, n = 6) and males (orange, n = 7). **i,** Peak velocity quantification (-0.5 to 0.5 s) for females (teal, n = 6) and males (orange, n = 7). **j,** Change in mean ΔF/F (last 10 min – first 10 min of the test) after closed arm entry (1 - 2 s) for females (teal, n = 6) and males (orange, n = 7). Dotted vertical line indicates timepoint where center-of-body marker crossed into the closed arm. Gray bar indicates the period used for panel **k**. **k,** Corresponding AUC quantification of change in ΔF/F for period indicated by gray bar in **j**. Two-sample t-test, *P* = 0.0284. All data are mean ± s.e.m; *: *P* < 0.05. For detailed statistical information, see Extended Data Table 1.

We also focused more closely on 5-HT^vHP^ activity around transitions in and out of the “safer” closed arms of the EPM. When mice retreated from center/open arms back into the closed areas (≥ 2 s in the center/open followed by ≥ 2 s in closed) activity of 5-HT^vHP^ neurons decreased in both sexes; however, this reduction was significantly attenuated in females (Fig. 5e-g), despite comparable velocities (Fig. 5h-i). This sex difference remained even when transitions accompanied by other scored behaviors during the peri-entry analysis window were excluded, with no significant effect of behavioral exclusion on the calculated AUC (Extended Data Fig. 6p). These results suggest that 5-HT^vHP^ population activity remains elevated in females upon return to a less aversive environment, indicating a sex difference in how this circuit disengages from anxiogenic contexts. Conversely, exiting the closed arms did not lead to a subsequent change in 5-HT^vHP^ population activity, and no differences between males and females were observed (Extended Data Fig. 6k-n).

During EPM exploration, locomotor activity typically declines over time, likely reflecting habituation as animals become familiar with the environment^81,82^. We therefore examined how the decrease in activity of 5-HT^vHP^ neurons during closed arms retreats evolved over the duration of the test. Specifically, we correlated AUC of the calcium signal during the 1 to 2 second period following closed arm entry (as in Fig. 5f) with elapsed time in the EPM for both female and male mice. In males, the decrease in 5-HT^vHP^ neurons population activity remained stable across the session (see example correlation Extended Data Fig. 6o). In contrast, in females, this suppression correlated with time (see example correlation Extended Data Fig. 6o): during the early phases of the test, 5-HT^vHP^ neuron activity remained high upon retreat into a closed arm, but a reduction gradually emerged as the session progressed and animals habituated to the test apparatus. Changes in AUC over time were quantified by subtracting the retreat-related response in the last 10 min of the EPM to response in the first 10 min (Fig. 5j). While activity of MRR 5-HT^vHP^ neurons remained stable in male mice, post-retreat responses in females evolved with time (Fig. 5j-k) showing that during early exploration of the EPM, female 5-HT^vHP^ neurons maintain elevated activity upon retreat into less aversive compartments of the maze. Together, these findings indicate that 5-HT^vHP^ neurons exhibit sex-specific activity dynamics during exploration and habituation in the EPM, with hyperexcitable female 5-HT^vHP^ neurons disengaging more slowly from aversive contexts.

### Hippocampal response to novelty is modulated by 5-HT selectively in female mice

Approach-avoidance conflict behaviors arise from competing drives to explore novelty and avoid aversive features^83^, a pattern readily observed in the EPM. We hypothesized that the sex-specific dynamics of MRR 5-HT^vHP^ neuron activity might partly reflect a delayed familiarization to a novel environment in females, and next interrogated the roles of 5-HT^vHP^ neurons in modulating the response to novelty. We therefore asked whether activation of these neurons alters the hippocampal network dynamics that normally accompany familiarization with a novel environment.

To measure the hippocampal response to novelty and its modulation by 5-HT we combined local field potential (LFP) recordings from both the dHP and vHP with optogenetic activation of 5-HT^vHP^ neurons. Theta oscillations (6-10 Hz) dominate the hippocampus LFP when mice are exploring an environment^84–86^. Theta oscillation properties exhibit a well-established linear relationship with locomotion, with increased speed being accompanied by higher oscillation frequency^87–90^. Two metrics can be extracted from the speed-frequency relationship: the slope of the linear relationship and its y-axis intercept. Prior work in the dHP has shown that environmental novelty reduces the slope, whereas familiarization is accompanied by progressive slope steepening^89,91,92^. In contrast, anxiolytic drugs consistently reduce the intercept, independent of their mechanism of action^93^, suggesting that this component reflects an anxiety-related contribution to hippocampal theta dynamics. We took advantage of the novelty- versus anxiety-related properties of the speed-theta frequency relationship to dissociate the impact of 5-HT^vHP^ neuron activation on either state.

Because these familiarity-related changes have been characterized primarily in the dHP, we first asked whether they also could be detected in the vHP under our experimental conditions. In an independent experiment without photostimulation, LFP were simultaneously recorded in the dHP and vHP while mice explored a novel square open field (Extended Data Fig. 8a-b). Theta frequency was linearly modulated by the animal’s speed in both the dHP and vHP (Extended Data Fig 8c). When comparing the first (novel, grey lines) to second 4-min exploration epochs (familiar, red lines), we found an increase in slope of the speed-frequency relationship (Extended Data Fig. 8d) while the intercept decreased (Extended Data Fig. 8e). These changes are consistent with progressive familiarization^89,91,93^ and establish that novelty-related theta dynamics can be detected in both dHP and vHP in our experimental paradigm.

We next tested whether optogenetic activation of 5-HT^vHP^ neurons modulates these novelty-and anxiety-related theta dynamics in males and females. During free exploration of a novel environment, LFP were recorded simultaneously from vHP and dHP during a 4-minute baseline epoch followed by a 4-minute 20 Hz optogenetic stimulation epoch, where 5-HT^vHP^ neurons were activated (Fig. 6a-c, Extended Data Fig. 9). We first controlled for the absence of altered locomotor behavior in response to light delivery (Extended Data Fig. 10). In line with prior reports that DR, but not MnR, 5-HT neurons modulate rodent locomotion, we observed a minor and short-lived (less than 10 seconds) decrease in velocity in female mice at stimulation onset (Extended Data Fig. 10a-b)^7,65,94,95^. When the full duration of optogenetic activation was considered, we observed no major changes in locomotor behavior (Extended Data Fig. 10e-g). Overall, optogenetic activation did not produce sustained locomotor suppression allowing the relationship between theta frequency and running speed to be assessed independently of gross changes in locomotor activity.

**Figure 6:**
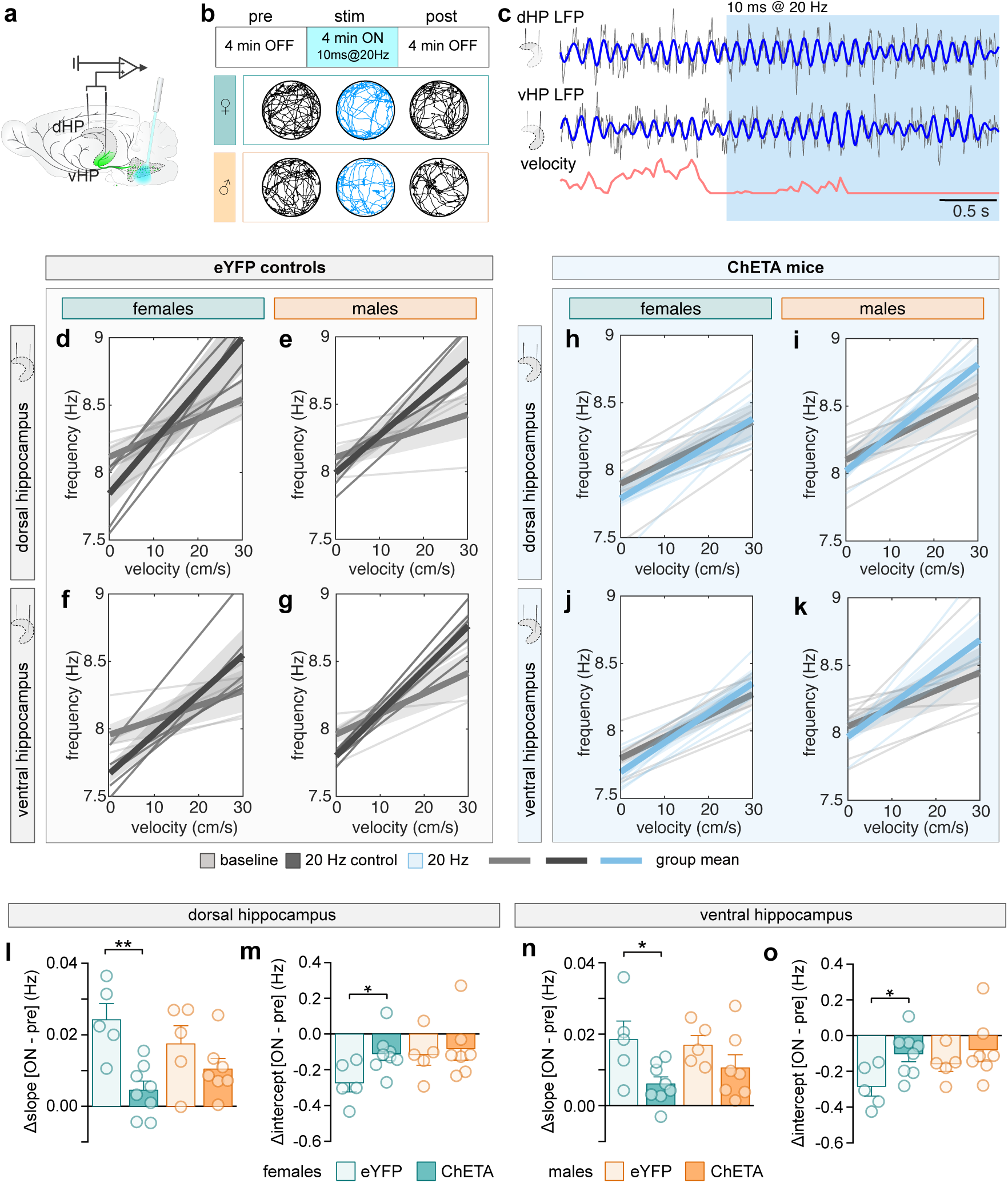
Activation of 5-HT^vHP^ neurons disrupts theta-velocity relationship selectively in females. **a,** Schematic of simultaneous multi-site local field potential recording with pathway specific optogenetic stimulation. **b,** Stimulation paradigm and example trajectories for a female (top) and male (bottom) animal for baseline (pre) and stimulation (stim) period. **c,** Example raw LFP traces (gray) recorded from dHP (top) and vHP (middle) and corresponding velocity (bottom), for animals exploring the open field, 2.5 s before and after stimulation onset. Blue: theta filtered traces. Red: instantaneous animal velocity. **d – g,** Instantaneous theta frequency as a function of running speed for eYFP control mice in dHP (**d**, females n = 5 and **e**, males n = 5) and vHP (**f**, females n = 5 and **g**, males n = 5). Speed frequency relationship was computed during pre (0-4 min, gray) and stim (4-8 min, black) epochs. Thin lines: individual animals; thick lines: group mean. Shaded areas represent s.e.m. **h – k,** Same as **d-g** in ChETA mice in dHP (**h**, females n = 8 and **i**, males n = 7) and vHP (**j**, females n = 8 and **k**, males n = 7), for pre (0-4 min, gray) and stim (4-8 min, blue) epochs. **l,** For the dHP, difference between slopes during optogenetic stimulation and baseline epochs. Two sample t-test: females (ChETA vs control), *P* = 0.0016. **m,** For the dHP, difference between intercepts during optogenetic stimulation and baseline epochs. Two-sample t-test: females (ChETA vs control), *P* = 0.0278. **n,** Same as **l**, for vHP. Two sample t-test: females (ChETA vs control), *P* = 0.0251. **o,** Same as **m**, for vHP. Two sample t-test: females (ChETA vs control). *P* = 0.0228. All data are shown as mean ± s.e.m; *: *P* < 0.05, **: *P* < 0.01; dHP: dorsal hippocampus; vHP: ventral hippocampus. For detailed statistical information, see Extended Data Table 1.

As expected, dHP theta frequency linearly scaled with running speed. In eYFP controls the slope of this relationship increased from the first 4 minutes of exploration (baseline, grey lines) to the subsequent 4 minutes (stimulation epoch, black lines) in both females (Fig. 6d) and males (Fig. 6e, see Extended Data Fig. 9a-d for representative recordings and power spectrums). A similar pattern was observed in the vHP (Fig. 6f-g, Extended Data Fig. 9e-h). Photoactivation of 5-HT^vHP^ neurons had sex-specific effects on the modulation of hippocampal novelty response. Male ChETA mice showed the same progressive slope increase as eYFP controls in both hippocampal regions (Fig. 6i,k,l,n). In contrast, female ChETA mice failed to show the familiarity-related slope increase in either the dHP (Fig. 6h,l) or vHP (Fig. 6j,n). Thus, activation of 5-HT^vHP^ neurons selectively disrupted the normal evolution of the theta frequency-velocity slope during familiarization in females.

Moreover, theta frequency at intercept, a metric modulated by anxiolysis, was also affected by 5-HT^vHP^ neuron activation in a sex-specific manner. In males, the change in intercept across epochs was comparable between eYFP controls and ChETA groups in both dHP (Fig. 6m) and vHP (Fig. 6o). By contrast, photoactivation of 5-HT^vHP^ neurons prevented the decrease in intercept observed in female eYFP mice, in both the dHP (Fig. 6m) and the vHP (Fig. 6o), suggesting a sex specific effect of increasing 5-HT^vHP^ neuron activity on anxiety levels. Taken together, activation of this pathway in females attenuates the normal vHP and dHP activity changes associated with familiarization during exploration of a new environment. These findings show that 5-HT^vHP^ neurons selectively influence theta oscillations properties underlying novelty processing and anxiety-related behavior in a sex-dependent fashion.

Using double retrograde tracing we confirmed that, under our experimental conditions, a subset of 5-HT^vHP^ neurons also innervate the dHP, likely contributing to the similar modulation of theta oscillations observed in both dHP and vHP during 5-HT^vHP^ activation (Extended Data Fig. 11).

Altogether, these findings demonstrate that, in a novel and low-threat environment, 5-HT^vHP^ neurons shape circuit-level signatures of novelty in a sex-dependent fashion, without prominent influence on locomotion. In addition, sex-specific changes in the intercept of the speed-theta frequency relationship suggest that increased 5-HT^vHP^ neurons activity modulates anxiety levels predominantly in females.

### Sex-specific M-current underlies excitability differences in MRR 5-HT^vHP^ neurons

Finally, we sought to identify the cellular mechanism underlying sex differences in intrinsic excitability and recruitment during anxiety-related behavior of female MRR 5-HT^vHP^ neurons. Kv7/M channels, encoded by the KCNQ genes, are major determinants of subthreshold membrane excitability. These channels mediate the M-current, a non-inactivating voltage-gated K+ current that is active near resting membrane potential and dampens neuronal excitability by stabilizing membrane potential and suppressing spontaneous and burst firing^96–99^. Previous studies have shown that Kv7/M channels are heterogeneously expressed in DR and MRR 5-HT neurons, where they regulate firing frequency and spike frequency adaptation^100–103^. Notably, M-current-expressing neurons are mostly concentrated in the dorsal part of the MnR, where 5-HT^vHP^ neurons were preferentially localized (Fig. 1a-c, Extended Data Fig. 1a-c). Whether functional M-current differs between male and female 5-HT^vHP^ neurons, however, remains unknown.

We hypothesized that sex-specific expression of the M-current contributes to the difference in excitability between male and females MRR 5-HT^vHP^ neurons. To test this, we examined the effects of M-current blockade on the intrinsic electrophysiological properties of identified MRR 5-HT^vHP^ neurons (Fig. 7, Extended Data Fig. 12). We first identified the presence of M-current by using the selective blocker XE991. The XE991-sensitive current amplitude was quantified as the difference between tail currents before and after bath application of 20 μM XE991 at repolarizing steps between –20 mV to – 60 mV. Neurons exhibiting a relaxation current greater than 20 pA at –50 mV were classified as M-current-positive. We found that all recorded male MRR 5-HT^vHP^ neurons (10/10 cells) exhibited functional M-current. In contrast, in females XE991-sensitive currents remained below 20 pA across all repolarizing steps, with the exception of one neuron (1/10 cells).

**Figure 7:**
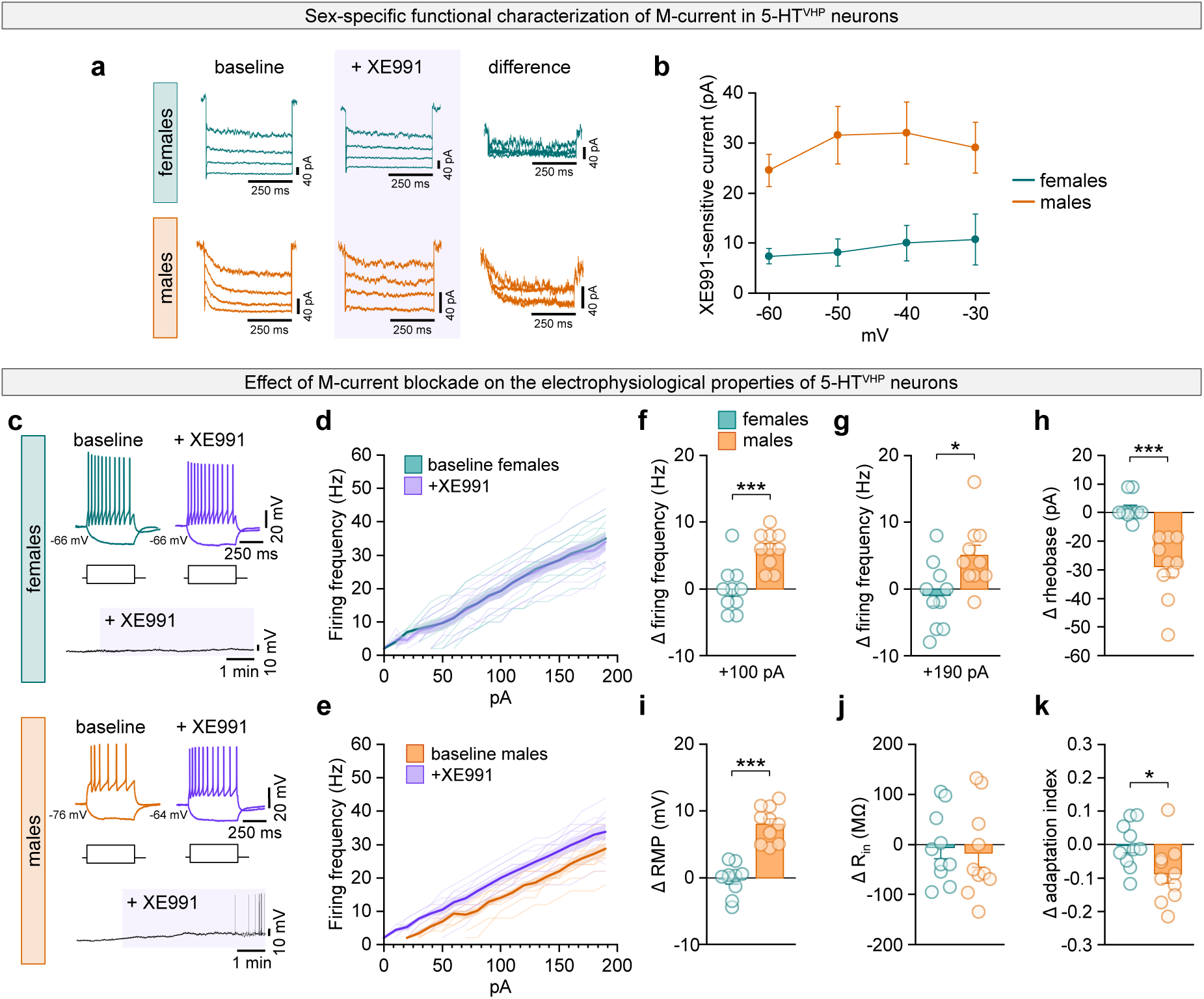
Sex-specific M-current underlies differences in MRR 5-HT^vHP^ neuronal excitability. **a**, Representative current traces at baseline (left), with M-current blocker XE991 (middle), and the corresponding XE-991-sensitive components (difference between baseline and XE991 traces, right). **b**, Average amplitude of XE-991-sensitive currents as a function of the repolarizing voltage steps for females (teal, n = 10 cells from 7 mice) and males (orange, n = 10 from 9 mice). **c**, Examples of membrane potential responses to current steps (-30 pA for lower trace and + 190 pA for upper trace) as part of the electrophysiological characterization of eYFP+ neurons recorded from female (top panel) and male (bottom panel) mice in the MRR at baseline (teal or orange) and with M-current blocker XE991 (purple); below, examples of time course change in resting membrane potential following bath application of XE991. **d-e**, firing frequency in response depolarizing current injection at baseline (teal) and with XE991 (purple) in females (**d**), and males (orange, baseline; purple, XE991, **e**). **f-k**, Change (+XE991 – baseline) in firing frequency at +100 pA, two-sample t-test *P* = 4.4019e-4 (**f**), firing frequency at +190 pA, two-sample t-test *P* = 0.0124 (**g**), rheobase, two sample t-test *P* < 0.0001 (**h**), resting membrane potential, two-sample t-test *P* < 0.0001 (**i**), input resistance (**j**) and adaptation index, two-sample t-test *P* = 0.0299 (**k**) for females (teal, n = 10 cells from 7 mice) and males (orange, n = 10 cells from 9 mice). Data are shown as mean ± s.e.m. *: *P* < 0.05, **: *P* < 0.01, ***: *P* < 0.001; For detailed statistical information, see Extended Data Table 1.

We next examined the effect of M-current blockade on the electrophysiological properties of male and female MRR 5-HT^vHP^ neurons (Fig. 7c-k, Extended Data Fig. 12). Consistent with the near absence of M-currents in females, application of XE991 did not alter their electrophysiological properties (Fig. 7d,f-k). By contrast, XE991 application significantly heightened the excitability of 5-HT^vHP^ neurons in male mice. M-current blockade increased firing frequency to levels comparable to female MRR 5-HT^vHP^ neurons (Fig. 7e-g and Extended Data Fig. 12m-n), reduced rheobase (Fig. 7h), depolarized the resting membrane potential (Fig. 7i), and decreased adaptation index (Fig. 7k) without affecting input resistance (Fig. 7j). Moreover, following XE991 application 40% of male neurons (4/10 cells) exhibited spontaneous firing.

Together, these findings demonstrate that functional M-current is present in male but largely absent in female MRR 5-HT^vHP^ neurons. By selectively constraining the excitability of male neurons, M-current contributes directly to the sex- and pathway-specific differences in intrinsic excitability of MRR 5-HT^vHP^ neurons.

## Discussion

Mood and anxiety disorders disproportionately affect women, yet the neural substrates driving this sex bias remain poorly defined. Here, we identify serotonergic projections from the MRR to the vHP as a key neural substrate underlying sexually divergent anxiety responses. By integrating viral circuit tracing, retrograde fiber photometry, *in vivo* local field potential recordings, pathway specific optogenetics and patch-clamp electrophysiology, we demonstrate that this circuit displays robust sex differences that converge across multiple levels of analysis. Female MRR 5-HT^vHP^ neurons display increased intrinsic excitability, greater functional recruitment and a causal role in promoting anxiety-like responses. These differences are underpinned by sex-specific Kv7/M-channel activity, with M-current selectively dampening the excitability of male MRR 5-HT^vHP^ neurons. These findings offer a mechanistic framework for understanding how serotonergic circuits can differentially shape anxiety-related behaviors in females and contribute to sex-specific risk for mood disorders.

Consistent with previous work^13,76,104,105^, we find that 5-HT^vHP^ neurons are distributed over multiple raphe subregions, notably the B9, MRR and DRI, with no differences in total number of 5-HT or 5-HT^vHP^ neurons between sexes. Our *ex vivo* electrophysiological recordings revealed that MRR 5-HT^vHP^ neurons are intrinsically more excitable in females, exhibiting increased spontaneous activity, more depolarized resting membrane potentials and higher firing rates compared to males. These sex differences in excitability seem restricted to the vHP-projecting subpopulation, as 5-HT^non-vHP^ neurons exhibited comparable excitability in both sexes. DRI 5-HT^vHP^ neurons in females also show depolarized membrane potentials, suggesting that intrinsic excitability may be an emergent property of subsets of serotonergic populations with shared developmental origins, connectivity and functions^11,106,107^. A few studies also report sex differences in the excitability of DR 5-HT neurons, but without specifying localization within DRI or projection targets^61,62^. In contrast, 5-HT^vHP^ neurons from the B9 group showed no sex differences in intrinsic properties. Together, these findings highlight striking electrophysiological heterogeneity in raphe-vHP circuits arising from B9, MRR and DRI. These raphe-vHP circuits might also show neurochemical diversity. The MRR and B9 contain 5-HT neuron populations of different embryonic origins (derived from rhombomeres 1, 2 and 3)^108,109^, that diverge into neurochemically distinct subtypes. Notably, the MnR contains serotonergic neurons co-expressing the vesicular glutamate transporter VGLUT3, which project to the vHP^109–112^, raising the possibility that a subset of the virus-transfected neurons in our study co-release glutamate and serotonin^53,113,114^. These diverse properties of raphe-vHP populations suggest that vHP functions are modulated by multiple specialized raphe circuits.

Our data link the intrinsic excitability of female MRR 5-HT^vHP^ neurons to heightened functional activation *in vivo*. Consistent with increased baseline firing rates in MRR 5-HT^vHP^ neurons observed in slice electrophysiology, c-fos mapping revealed greater recruitment of MRR 5-HT^vHP^ neurons in female mice following optogenetic stimulation and behavioral testing. Given that c-fos is an indirect marker of neuronal recruitment, we further examined this relationship *ex vivo*. Despite comparable photocurrent amplitudes, female MRR 5-HT^vHP^ neurons exhibited higher action potential fidelity during 20 Hz stimulation. In addition, *in vivo* fiber photometry recordings from MRR 5-HT^vHP^ neurons revealed that this projection-defined population is dynamically, and sex-specifically, engaged during transitions into the more aversive compartments of the EPM. While 5-HT^vHP^ activity declined upon retreat into closed arms in both sexes, this decline was attenuated in females and emerged only gradually over the session. This delayed disengagement suggests a sex-specific persistence of neuronal activation, potentially reflecting slower updating of internal safety predictions under environmental novelty or threat. Strikingly, 5-HT^vHP^ neurons also displayed ramping activity in the seconds preceding exploration of the more aversive parts of the EPM, reminiscent of DR neuron ramping activity patterns observed during anticipation of reward or punishment^41,115–117^. Such progressive increases in DR 5-HT neuron activity over long timescales have recently been proposed to reflect a prospective coding for value^42^, a framework that possibly extends to MRR 5-HT neurons. Notably, MRR and DR 5-HT neurons respond in opposite directions to rewarding stimuli^47^, and also differ during locomotion: whereas DR neurons decrease their activity during locomotion bouts in the open field or avoidance tests^95^, our findings confirm that in the same settings, MRR activity increases.

At the network level, targeted optogenetic activation of 5-HT^vHP^ neurons selectively disrupted dHP and vHP theta dynamics in female mice. Stimulation flattened the theta frequency-running speed relationship, mirroring the pattern typically seen during novelty exposure and consistent with a failure to habituate to the environment^89,91,93^. In contrast, control animals and male ChETA mice exhibited the expected progressive slope increase over time, reflecting familiarization. These findings extend previous results showing serotonergic modulation of hippocampal theta^118–121^ by demonstrating a sex-dependent effect. Given that, during anxiety states, theta synchrony facilitates long-range communication between the hippocampus and other regions, such as the PFC^28,29,122^, serotonergic modulation of vHP theta could maintain persistent vigilance. This interpretation is supported by our anatomical mapping which revealed collateral projections from 5-HT^vHP^ neurons to the PFC, raising the possibility that their activation coordinates distributed circuits implicated in anxiety regulation.

Behaviorally, optogenetic activation of 5-HT^vHP^ neurons increased anxiety-like behavior selectively in females. In ethologically relevant tests, including the EPM and SA, female ChETA mice exhibited increased avoidance of aversive zones, reduced risk-assessment behaviors such as head-dipping and stretch-attend postures, and increased stress-related grooming. Notably, grooming was selectively elevated in anxiogenic environments but unaffected in the open field, highlighting the context dependence of this circuit’s behavioral impact, in line with past work^46^. The observation that grooming episodes were associated with rapid decreases in calcium signal in both sexes supports the view of grooming as an adaptive displacement behavior^72,123,124^, which transiently suppresses serotonergic tone.

Importantly, the behavioral effects of 5-HT^vHP^ neuron activation occurred in the near absence of locomotor changes, supporting a specific role in anxiety modulation behavior rather than general arousal. While broad activation of the MnR has previously been shown to increase anxiety-like behavior^8,44,65^, our data refine this understanding by demonstrating that selective stimulation of the vHP projection pathway is sufficient to amplify anxiety-related behaviors in females. Importantly, our c-fos data show robust activation of 5-HT neurons in both sexes and across all raphe sub-regions, indicating that the absence of an anxiety-related behavioral outcome in males cannot be attributed to inefficient optogenetic activation. This dissociation between efficient optogenetic activation and behavioral outcome suggests that, in males, the vHP-projecting serotonergic pathway primarily contributes to behavioral domains not captured by the behavioral paradigms used here. These sex-specific behavioral outcomes further underscore the critical role of the MRR in driving anxiety-like behavior. While B9 and DRI are recruited to a similar extent in males and females, differential recruitment of a small sub-population of MRR serotonergic neurons was sufficient to generate robust sex differences in anxiety-like behavior.

The MnR provides dense serotonergic innervation to a wide array of forebrain regions^76^, and individual 5-HT neurons collateralize extensively^104,105^. Although 5-HT^vHP^ neurons also collateralize to other forebrain regions, including the dHP and lateral septum, which could contribute to their behavioral effects, we found no sex differences in collateral density or distribution, suggesting that differential downstream targeting is unlikely to account for the sex-specific responses. The postsynaptic mechanisms mediating serotonergic modulation of vHP outputs remain unknown. Given that virtually all 5-HT receptor subtypes are expressed in the vHP^125^, serotonergic input to this region could act on multiple parallel output pathways, including to anxiety-related regions such as prefrontal cortex, lateral hypothalamic area or lateral septum^25,126–129^. The details of 5-HT modulation of vHP long-range principal cells or interneurons, and their impact on sex specific anxiety responses await clarification.

Finally, we propose that sex-specific expression of M-currents is the mechanism underlying heightened excitability of female MRR 5-HT^vHP^ neurons. Kv7/M channels modulate excitability of serotonergic neurons in the DR and MRR, yet whether their expression differs between sexes has never been investigated^101,102^. We found that M-currents are present in all recorded male MRR 5-HT^vHP^ neurons but are largely absent in females. Pharmacological blockade of M-current selectively increases neuronal excitability of male neurons, highlighting Kv7/M channel activity as a key determinant of sex-specific excitability in the MRR-vHP serotonergic pathway. The absence of Kv7/M channel activity in females is consistent with the heightened recruitment of 5-HT^vHP^ neurons during baseline and optogenetically driven anxiety-like behavior. Whether the sex-specific expression of the M-current is driven by differential KCNQ genes expression or Kv7 subunit compositions remains to be determined. Overall, our findings advance an emerging body of work delineating the roles of topographically organized serotonergic circuits in the modulation of emotion-related behaviors^18,47,130,131^. Our study directly links hyperexcitability of the female MRR 5-HT^vHP^ pathway to enhanced anxiety-like responses. These findings align with the higher prevalence of anxiety and mood disorders in women^132–134^, and suggests that sex-specific serotonergic circuit function may constitute a core vulnerability factor across affective disorders. By delineating a projection-specific serotonergic pathway that differentially modulates anxiety-related behaviors in females, our work highlights the potential for sex-informed, circuit-targeted interventions in mood and anxiety disorders.

## Acknowledgements

We thank Dr. Elke Küster-Schöck from the Centre Hospitalier Universitaire Sainte-Justine Imaging center for microscopy guidance, and Denise Carrier, Véronique Pelletier and the animal facility team for help with colony maintenance. We thank all members of the Amilhon lab for their valuable comments on the manuscript, and undergraduate students Clémence Brin, Camille Pons, Charlène Manti, Morgane Roger for their help with various aspects of the study.

## Contributions

L.P. performed patch-clamp electrophysiology experiments. S.v.d.V. performed LFP experiments. F.H. performed fiber photometry and double-projection experiments. A.-S.S. and F.P. performed optogenetic experiments. J.F.H. and A.-S.S. collected collateral dataset. A.-S.S., F.P. and F.H. performed c-fos experiments. E.L. performed vHP 5-HT fiber density experiments. S.v.d.V. and G.D. developed analysis pipelines and code. G.D. provided technical support for all experiments and statistical analysis. All authors analyzed data, performed statistical analysis and provided material for the manuscript. S.v.d.V., L.P., and B.A. prepared figures and wrote the manuscript with contribution and input from all authors. B.A. conceived and supervised all aspects of the project, and secured funding. L.P. and S.v.d.V contributed equally to this work.

## Ethics declarations

Competing interests: The authors declare no competing interests.

## Materials and methods

### Animals

Male or female SERT-Cre (B6.FVB(Cg)-Tg(Slc6a4-cre)ET33Gsat/Mmucd line, MMR,#031028-UCD;) transgenic mice were bred with female or male C57BL/6J mice (Jackson Laboratory, #000664) to generate experimental animals. All mice used in this study were between 60-90 days old at the time of experiments. Animals were group-housed on a 12-hour light/dark cycle at 24°C and 40% humidity with ad libitum access to food and water. All procedures involving animals were approved by the Comité Institutionnel des Bonnes Pratiques Animales en Recherche (CIBPAR) of the Centre de Recherche Azrieli du CHU Sainte-Justine (protocol 2023-5039), in accordance with the standards of the Canadian Council on Animal Care.

### Stereotaxic surgeries

Animals were anesthetized with isoflurane (5% induction, 0.5-1.5% maintenance) and placed in a stereotaxic frame (David Kopf instruments). Carprofen (2 mg/kg) was administered subcutaneously before surgery. Body temperature was maintained at 37°C with a heating pad (Stoelting) and eyes were hydrated with ocular gel (Optixcare, Aventix). Craniotomies were performed using a mountable drill (Stoelting). Viral injections were performed using a glass pipette backfilled with mineral oil connected to a Nanoject III (Drummond Scientific) at a flow rate of 1 nL/second. To target 5-HT^vHP^ neurons, a retrogradely transported adeno-associated virus (AAV) of serotype 2/9 was bilaterally infused in the vHP. Each hippocampus received 400 nL of viral solution injected at two sites (stereotaxic coordinates: [AP] -3.52, [ML] ±3.25, [DV] -4.17 and -3.47). All stereotaxic coordinates are taken from bregma (in mm).

MRR 5-HT^non-vHP^ neurons were distinguished from MRR 5-HT^vHP^ neurons using a dual-labeling strategy. SERT-Cre mice received 50 nL of AAV2/9-hSyn-DIO-mCherry into the MRR (stereotaxic coordinates: [AP] -4.48, [ML] -0.8, [DV] -4.53; Angle: 10°) to label all 5-HT neurons, followed by bilateral injections of AAV2/9-EF1a-DIO-eYFP into the vHP, as described above, to retrogradely label 5-HT^vHP^ neurons. 5-HT^non-vHP^ neurons were therefore identified as mCherry+/eYFP- neurons.

For electrode and optic fiber implantation surgeries, the skull was exposed and carefully cleaned with ethanol before craniotomies were drilled. Electrodes and/or optic fiber were then slowly lowered into position and maintained in place using a combination of C&B-Metabond and dental acrylic (Parkell) and 2-3 stainless-steel anchor screws (Antrin Miniature Specialties Inc). Electrodes and optic fiber stereotaxic coordinates are described in the corresponding sections below. Following surgery, mice were monitored for at least 3 days and allowed to recover for at least one week before subsequent procedures or behavioral testing.

### Fiber photometry recordings

To monitor activity-related calcium dynamics in 5-HT^vHP^ neuronal somata at the level of the MRR, SERT-Cre mice received bilateral injections of AAV2/9-Syn-Flex-GCaMP6s-WPRE-SV40 (CS1028, University of Pennsylvania Vector Core) targeting the vHP using the methodology described above, resulting in retrograde expression of the calcium indicator GCaMP6s in 5-HT^vHP^ neurons. Four weeks later, animals were implanted with a 200 µm low-autofluorescence optic fiber cannula (Neurophotometrics) targeting GCaMP6s transfected neurons in the MRR ([AP] – 4.20 to – 5.30 with a 10° rostro-caudal angle, [ML] 0.00, [DV] – 3.5 to – 4.45).

Following implantation and recovery, mice were habituated for at least one week. They were handled daily, connected to sham patchcords in individual home cages containing familiar bedding and nesting material, and then returned to their littermates before the dark cycle. Behavioral testing was conducted on separate days in either a in a 40 x 40 cm light grey square open field (40 x 40 cm light grey square) or an EPM consisting of two closed arms (30 x 5 x 19.8 cm) and two open arms (30 x 5 x 0.9 cm), and a 5 x 5 cm center zone. On each testing day, mice were first habituated for 1 hour in individual cages in the experimental room. After habituation, mice were connected to the patchcord and returned to the cage for 20 minutes during which photometry recording was initiated. Animals were then placed in the behavioral apparatus for 20 minutes of free exploration.

Excitation light (470 nm and 415 nm) was alternately pulsed at 10 Hz at ∼80 µW (measured at the fiber tip) using an FP3002 photometry console (Neurophotometrics). Arenas were cleaned with water and 70% ethanol between animals. Room illumination was maintained at 70 ± 10 lux. Fluorescence was recorded sequentially for both the 470 nm calcium-dependent signal and the 415 nm isosbestic control. Injection sites, viral expression, and fiber placements were verified post hoc by histology, as described below.

Bleaching and movement artifacts were corrected using the method previously described^135^. Briefly, we used the coefficient obtained from a linear fit (obtained from iteratively reweighted least squares regression) of the 415 nm signal to the 470 nm signal, to scale the 415 nm signal. The scaled 415 nm signal was subtracted from the 470 nm signal to obtain ΔF, and ΔF was divided by the scaled 415 nm signal to yield ΔF/F. Signals were z-scored across the recording session and low-pass filtered at 3 Hz prior to further analysis. Peri-event plots were generated for each mouse by averaging z-scored ΔF/F around behavioral events and subtracting the baseline (average signal 1-2 s prior to the event). Within-mouse averages were then averaged across animals to obtain group means. These normalization steps express fluorescence changes relative to each animal’s own baseline signal.

Mouse body parts were tracked using SLEAP^136^, and the center of the body was used for all subsequent analyses. Animal positions were filtered to reduce jitter: the animal position at a specific frame was updated only if the raw animal position for that frame was located further than ½ of the mouse length away from the previous position. Movement initiation was defined as the first frame where velocity exceeded 1 cm/s, provided the mouse moved < 2 cm in the preceding 1 s period and > 10 cm in the subsequent second.

For the transitions between closed and open arms in the EPM (Fig. 1d-g), closed→open transitions were defined as the mouse spending at least 10 s in a closed arm before entering center/open arms for ≥ 2 s. Closed→closed transitions were defined as transitions between closed arms with ≤ 1 s spent in the center. Changes in fluorescence were aligned to the timepoint at which center-of-body crossed the closed arm-center boundary. For time-locked analysis of closed-arm entries and exits (Fig. 1h-s), arm entries were included in the analysis if the mouse spent ≥ 2 s in the center/open arm before entering the closed arm, and ≥ 2 s in the closed arm after entry. Exit transitions were defined similarly, requiring at least 2 s in the closed arm before transition and at least 2 s in the center or open arm after exit. Grooming onset and offset were manually scored frame-by-frame using custom Matlab code.

### Local field potential recordings

SERT-Cre mice received bilateral vHP injection of either AAV2/9-EF1a-DIO-hChR2(E123T/T159C)-eYFP (#35509-AAV9, Addgene), to cause CRE-dependent expression of the excitatory opsin ChETA in vHP projecting 5-HT neurons, or AA2/9-EF1a-DIO-eYFP (#27056-AAV9, Addgene) for control mice, using the methodology described above. Three weeks after virus injection, animals were implanted with a 200 μm optic fiber (#FG200UEA, Thorlabs) above the MRR ([AP] -4.35, [ML] -0.55 at 7° angle, [DV] -3.23), and custom recording electrodes in the hippocampus. For local field potential (LFP) recordings, custom bipolar electrodes were constructed using 50 µm PFA coated tungsten wire (#585088, A-M Systems) and targeted to the ventral ([AP] -3.65, [ML] 3.50, [DV] -3.00) and dorsal hippocampus ([AP]: -1.58, [ML] 1.40, [DV] -1.50). A ground screw was placed over the cerebellum and an EMG electrode was implanted in the trapezius muscle. All electrode and fiber placements were confirmed by histology, as described below. LFP were recorded and digitized at 20 KHz using an Open Ephys system^137^, while mice explored a novel, circular open field (39 cm diameter) or a novel, square open field (40 cm).

Videos were processed with DeepLabCut^138,139^ (version 2.3rc3) to extract body positions, using the body center as the animal’s position. LFP recordings were down sampled to 1 kHz and aligned with the behavioral tracking data. LFPs were band-pass filtered (5-10 Hz) and subjected to a Hilbert transform to obtain the analytic signal. The power of the signal was taken as the magnitude of the analytic signal squared. The instantaneous phase of the analytic signal was extracted and unwrapped to provide an estimation of instantaneous theta frequency. Periods with power >6 SD were considered movement artefacts and were removed from analysis along with the flacking 250ms of data on each side.

Animal positions were filtered to reduce jitter as described in the fiberphotometry section. Running speed was computed from the filtered positions by dividing Δd/Δt where d is the distance and t is time. Finally, to characterize the relationship between theta frequency and running speed, the median of the instantaneous frequency was taken over a time window of approximately 33.3ms to yield a frequency estimate corresponding to each frame of the video (sampled at 30 Hz). A first order polynomial was fitted to theta frequency and animal speed between 1 and 30 cm/s to extract the slope and intercept of the speed – theta frequency relationship.

For visualization of stimulation-evoked LFP activity, multitaper spectrograms were calculated for the 10 s preceding and 10 s following photostimulation onset using Chronux *mtspecgramc* function, with a moving window of [1 0.5] s, tapers [1 1], and a frequency range of 2-30 Hz. Representative raw and theta-filtered (5-10 Hz) LFP traces were extracted from the 2.5 s immediately preceding and following stimulation onset. To assess stimulation related changes in spectral profile, power spectra were calculated separately for the pre-stimulation and stimulation epochs using the Chronux *mtspectrumc* function using non-overlapping 0.5 s epochs during running (>10 cm/s). Spectra were averaged across pre and stimulation periods for each animal and normalized.

### Optogenetic manipulations and behavior

5-HT^vHP^ neurons of SERT-Cre mice were transfected with either the excitatory opsin ChETA or eYFP, and an optic fiber was implanted as described above. Mice were allowed a 2-week recovery period post-implantation, followed by 5 days of handling (∼10min/day) to minimize stress related to the experimenter, the behavior room or the fiber optic connection on the day of the experiment. On testing days, mice were transferred to the experimental room ≥ 45 min before behavioral assays and placed in individual cages containing familiar bedding and nesting material. All experiments were conducted between 9 AM and 2 PM to reduce circadian variability.

Behavioral testing occurred in the following order with 48 h intervals between tests: open field, EPM, and SA. Arenas were cleaned with 70% ethanol between mice to reduce olfactory cues, and light sensitivity was maintained at 70 ± 5 lux at the center of each apparatus. Mice were allowed to freely explore the maze for 12 min, with optogenetic stimulation delivered during minutes 4-8. Stimulation consisted of blue light (450 nm) pulses at 20 Hz (10 ms pulse width) at ∼12-15 mW measured at the fiber implant tip. Viral expression and fiber placement were confirmed post hoc (see Histology).

Mouse body parts were extracted from each video using SLEAP^136^, using the center of the body as the animal’s position. From this raw animal position, the time spent and the number of entries in each zone of the different mazes could be computed. Entries into a zone were recorded if the raw animal position was within the zone for at least 1s. To compute the distance traveled and animal velocity, we filtered the raw animal position as described in the previous section. From this filtered position, the distance traveled was computed as the sum of the frame-to-frame displacement, and velocity was obtained by multiplying frame-by-frame displacements by the video sampling frequency. The time immobile was taken as the total time where the animal was not locomoting and the mean velocity while moving was calculated from all periods of locomotion.

For the successive alleys an occupancy index was calculated as:

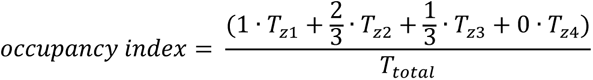

Where *T*_z1_ -*T_z4_* are time spent in zones 1-4 and *T_total_* is total test duration. Values near 1 indicate that mice spent most of their time in less aversive zones (zone 1). Values near 0.5 reflect uniform exploration across zones. These scores can also be interpreted as the normalized mean position of the mouse along the alleys from zone 1 to zone 4. Grooming, rearing, stretching and head-dipping behaviors were manually scored by the same experimenter blind to treatment (eYFP or ChETA).

### Histology

For all immunohistochemistry experiments, mice were deeply anesthetized with an intraperitoneal injection of ketamine/xylazine/acepromazine (80/12.5/2.5 mg/kg, respectively). Animals were transcardially perfused with 45 mL of 4% paraformaldehyde (PFA) in phosphate-buffered saline (PBS), and brains were extracted and postfixed in PFA at 4°C for 24-48 h. For cryosectioning, brains were cryoprotected using 15% sucrose in PBS for 24 h, embedded in Tissue-Plus™ O.C.T. Compound (#23-730-571, Fisher Scientific), and frozen in isopentane. Standard immunohistochemistry was performed using antibody solutions prepared in PBS with 0.45% gelatin and 0.25% triton. Image processing and analysis were performed using FIJI software (ImageJ, version 2.14.0). When necessary, Tph2 immunolabelling was used to delineate the raphe nuclei.

#### Histological validation of viral expression, electrode and fiber placement

For anatomical validation of viral expression and optical fiber implantation sites, either frozen coronal sections (25 μm) were prepared and cut on a cryostat (Leica Biosystems CM3050 S) or fixed, non-frozen coronal sections (40 μm) were prepared and cut on a vibratome (Leica VT1200), and stained for eYFP or GCaMP6s or mCherry and Tph2 as described above, using the following primary and secondary antibodies: anti-Tph2 in guinea pig (1:1000; #TPH2-GP-Af900, Frontier Institute), anti-GFP in chicken (1:2000; #A10262, Invitrogen), anti-RFP in rabbit (1:10000; #200-101-379, VWR (Rockland)), anti-guinea pig A647 in goat (1:2000; #A21450, ThermoFisher) anti-chicken A488 in donkey (1:2000; #703-545-155, Jackson ImmunoResearch), and anti-rabbit A555 in donkey (1:2000; #A31572, Life Technologies). Sections were mounted on slides using Fluoromount-G containing DAPI (ThermoFisher). Images were acquired using a Leica DMi8 wide-field inverted microscope with 10X and 20X objectives, equipped with a Leica DFC9000 camera, and LAS X software (Leica). One-quarter of the sections were stained for eYFP or mCherry and Tph2, as described above, to assess viral expression and optical fiber placement, while the remaining three quarters were directly mounted to assess HP electrode placement. These sections were imaged using the Zeiss AxioScan.Z1 with 10X objective and ZEN Microscopy Software (Zeiss).

#### Collaterals

To assess the proportion of 5-HT neurons projecting to both the dHP and vHP (Extended Data Fig. 4), adult SERT-Cre males (n = 3) and females (n = 4) were injected as described above with AAV2/9-EF1a-DIO-hChR2(E123T/T159C)-eYFP (#35509-AAV9, Addgene) in the left vHP (2 injections of 400 nL of virus). Five weeks later, neurons projecting to the left dHP were retrogradely labeled with CTB-A647, using 100 µg of CTB-A647 previously suspended in 20 µl of PBS1x at 4 °C and short-term stored at 4 °C. dHP injections were performed at: [AP] -1.90 and -2.40, [ML] -1.35 and -2.00, [DV] -1.55 and -1.65. Mice were sacrificed 10 days post CTB injection. Coronal sections were cut (35 µm) on a vibratome (Leica VT1200) and labeled for eYFP and Tph2 using the following primary and secondary antibodies: anti-GFP in goat (1:5000; #NB100-1770, Novus Biologicals), anti-Tph2 in rabbit (1:2000; #NB100-74555, Novus Biological), anti-goat-A488 in donkey (1:2000; #A11055, Life Technologies) and anti-rabbit-A555 in donkey (1:2000; #A31572, Life Technologies). The distribution of HP-projecting 5-HT neurons was quantified on sections collected at 105 µm intervals along the rostro-caudal axis of the raphe nuclei (B5-B9).

For qualitative assessment of 5-HT^vHP^ collaterals (Extended Data Fig. 7), a separate group of SERT-Cre males (n = 4) and females (n = 4) was used. 5-HT^vHP^ neurons were transfected using bilateral injections as described above. Free-floating coronal and horizontal slices (25 μm) were cut on a cryostat (Leica Biosystems CM3050 S) and labeled for eYFP and 5-HT using the following primary and secondary antibodies: anti-GFP in chicken (1:2000; #A10262, Invitrogen), anti-5-HT in goat (1:2000; #ab66047, Abcam), anti-chicken-A488 in donkey (1:2000; #703-545-155, Jackson ImmunoResearch) and anti-goat-A555 in donkey (1:2000; #A21432, Thermo Fisher Scientific). All regions with collaterals were identified using anatomical landmarks and a mouse brain atlas^140^ as reference. All images were acquired using a Leica DMi8 wide-field inverted microscope as described above.

#### C-fos assessment of neural activity

C-fos expression was assessed in mice expressing ChETA or eYFP in 5-HT^vHP^ neurons. Mice were sacrificed 70 minutes after completing 12 minutes in the SA paradigm with optogenetic stimulation performed as described above. Fixed, non-frozen brains were cut into 40 µm sections using a vibratome (Leica VT1200) and triple-labeled for c-fos, eYFP and Tph2 using the same protocol, with the addition of anti-c-fos in rabbit (1:2000; #2250, Cell Signalling) and anti-rabbit –A555 in donkey (1:2000; #A31572, Life Technologies). The distribution of 5-HT+/c-fos+/eYFP+ neurons was quantified on sections collected at 120 µm intervals along the rostro-caudal axis of the raphe nuclei.

#### Quantification of 5-HT fiber density in the vHP

Adult C57BL/6 male (n = 6) and female (n = 7) fixed, non-frozen brains were cut using a vibratome (Leica VT1200) and 40 µm coronal sections of the vHP were collected and labeled with anti-5-HT in goat (1:2000; #ab66047, Abcam) and anti-goat A488 in donkey (1:2000; #A11055, Life Technologies). Images were acquired using a Leica TSC SP8 confocal microscope with 40 X objective. ROIs were acquired on a single focal plane across CA1, CA3, DG and VS areas, and analyzed using FIJI. Briefly, background was subtracted, images were binarized and % of surface occupied by 5-HT staining was extracted.

### Patch-clamp

Mice were deeply anesthetized with an intraperitoneal injection of ketamine/xylazine/acepromazine (80/12.5/2.5 mg/kg, respectively) and brains were prepared for sectioning on a vibratome using a modified protocol from Ting and colleagues^141^. Each mouse was transcardially perfused with an NMDG-based solution (in mM: 93 NMDG, 93 HCl, 2.5 KCl, 1.2 NaH2PO4, 30 NaHCO3, 20 HEPES, 25 glucose, 5 sodium ascorbate, 2 thiourea, 3 sodium pyruvate, 10 MgSO4, and 0.5 CaCl2 at pH 7.4, oxygenated with carbogen). Coronal sections (300 μm) were cut using a vibratome (Leica VT1200) and incubated at 32°C for 10 to 12 minutes in the same NMDG solution. Slices were then transferred to artificial cerebrospinal fluid (aCSF) at room temperature (in mM: 124 NaCl, 24 NaHCO3, 2.5 KCl, 1.2 NaH2PO4, 5 HEPES, 12.5 glucose, 2 MgSO4, and 2 CaCl2, pH 7.4, oxygenated with carbogen). During recordings, slices were continuously perfused with aCSF at 28°C at 3 mL/min.

5-HT^vHP^ neurons were visually identified via eYFP fluorescence, while 5-HT^non-vHP^ neurons were visually identified via mCherry fluorescence. Whole-seal recordings were obtained using an intrapipette solution (in mM: 126 K-gluconate, 4 KCl, 10 HEPES, 4 Mg2ATP, 0.3 Na2GTP, and 10 PO-Creatine, adjusted to pH 7.25 with KOH; 272 mOsm). Pipette had a resistance between 4 to 8 MΩ. Recordings were performed using an Axon Multiclamp 700B amplifier (Molecular Devices). After achieving whole-cell configuration, resting membrane potential was recorded for 4 min prior to additional protocols. In current-clamp, 500 ms hyperpolarizing steps were applied to assess hyperpolarization-activated current, passive membrane conductance and input resistance. 500 ms depolarizing current steps were applied to assess firing frequency, rheobase and action potential features. The adaptation index was calculated using the last depolarizing current step as:

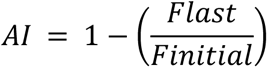

where Flast represents the average frequency of the last two interspike intervals (ISIs), and Finitial is the average frequency of the first three ISIs^142^. All membrane potentials were corrected offline for a liquid junction potential (LJP) of -15.8mV.

#### Optogenetic activation of 5-HT^vHP^ neurons ex vivo

ChETA-expressing 5-HT^vHP^ neurons were held at –55 mV in voltage-clamp and a ramp of increasing blue-light intensity (from 0 to 10.19 mW/mm^2^ intensity at fiber tip) was applied to measure steady-state photocurrent amplitude. In current clamp, trains of 10 ms blue-light pulse width were delivered at 20 Hz for 1 s to assess action potential fidelity.

#### Pharmacological blockade of M-current on 5-HT^vHP^ neurons electrophysiological properties

Neurons were held at –20 mV in voltage-clamp and 500 ms-long repolarizing steps were applied from –30 to -60 mV to measure M-current^98,143^. Resting membrane potential was recorded for minimum 2 min prior to bath application of 20 μM XE991 (Tocris Bioscience), a specific blocker of the M-current, to pharmacologically isolate the current^144^. The effect of drug application on resting membrane potential was recorded for the consecutive 5 min. Then, current-clamp and voltage-clamp protocols were repeated to measure the XE991-sensitive current and the effect of M-current blockade on input resistance and neuronal excitability. To identify the M-current, the amplitude of peak deactivation currents (tail currents) elicited by a 500 ms square voltage step to –30, -40, -50, and –60 mV from a holding potential of –20 mV were quantified. Tail current was calculated as the difference between the peak of a 30-ms segment, taken 10–40 ms into the hyperpolarizing step, and the average during the last 50 ms of that step^101,145^. The XE991-sensitive current was obtained as the digital subtraction of the control and XE991-resistant current traces. Neurons that presented a relaxation current greater than 20 pA at –50 mV were considered M-current-positive.

### Statistics

All data are presented as mean ± standard error of the mean (s.e.m.), and the corresponding statistical tests are described in the figure legends and in Extended Data Table 1, and Extended Data Table 2. Origin 2024 (OriginLab) was used for all basic statistical analysis, except for behavioral experiments described in Figure 4 and Figure 5, which were analyzed with R (version 4.4.1 in RStudio 2025.05.0).

Behavioral experiments were conducted on two independent cohorts, and therefore analyzed with three-way ANOVA with sex, treatment and cohort as factors. The ANOVA model included all main effects (sex, treatment and cohort) and interactions (sex * treatment, sex * cohort, treatment * cohort and sex * treatment * cohort). Post hoc tests were performed for planned comparisons (female-eYFP vs. male-eYFP, female-eYFP vs. female-ChR2, male-eYFP vs. male-ChR2 and female-ChR2 vs. male-ChR2) using Holm’s correction for multiple testing. Statistical tests were performed in R using the *aov* function (stats package) for the three-way ANOVA and the *emmeans* function (*emmeans* package) for post hoc comparisons. Homogeneity of variance was tested using Levene’s test *(leveneTest* function, *car* package) using median^146^. For clarity, only the plots of the marginal means are shown in the main text, while the full group breakdown is shown in Extended Data Table 1. A threshold of P < 0.05 was considered statistically significant. Figures use the following significance notation: *, *P* < 0.05; **, *P* < 0.01; ***, *P* < 0.001

---

**Extended Data Fig. 1:**
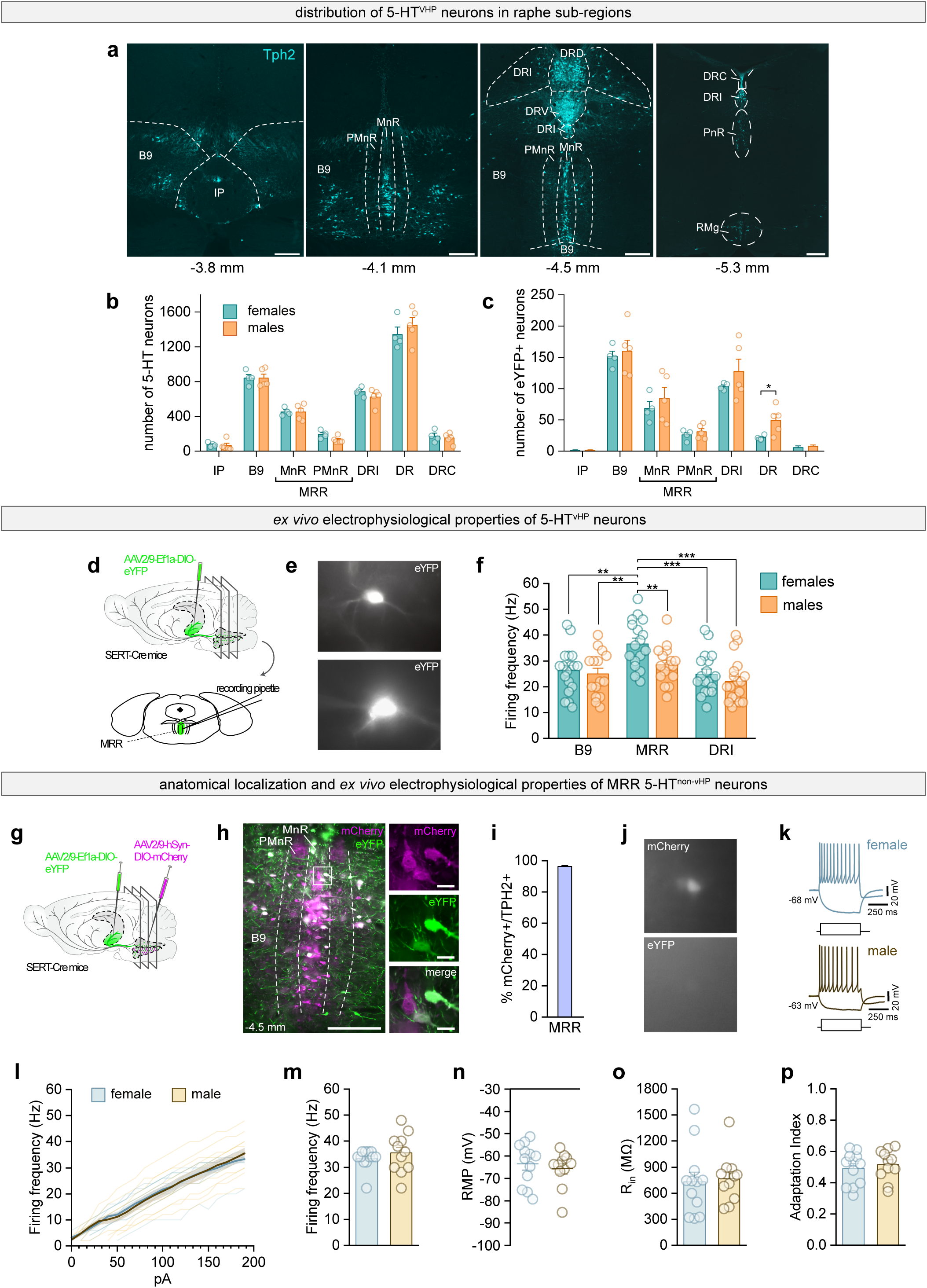
Sex-independent distribution of 5-HT^vHP^ neurons across raphe subregions. **a,** Schematic delineations of raphe regions with 5-HT+ neurons (Tph2+, blue). Distance from bregma shown in mm. Scale bars = 250 μm. **b,** Total number of 5-HT+ neurons for each raphe subregion, for both sexes (females: n = 4; males: n = 5). **c,** Total number of eYFP+ neurons per subregion, for both sexes. For the DR, two-sample t-test, *P* = 0.0394. **d,** Schematic of experimental strategy to assess *ex vivo* electrophysiological characteristics of 5-HT^vHP^ cells in raphe slices. **e,** Representative examples of eYFP-positive (eYFP+, white) cell. **f**, Same data as shown in Fig. 1f, l and r, grouped on the same graph for direct comparison and statistical testing, showing that female MRR 5-HT^vHP^ neurons have the highest firing frequency among all regions and sexes. Two sample t-tests, MRR: female vs male, *P* = 0.0089; Female B9 vs MRR, *P* = 0.0017; Female MRR vs DRI, *P* = 0.0003; Female MRR vs male DRI, *P* = 0.0010; Female MRR vs Male B9, *P* < 0.0001. **g**, Schematic of experimental configuration: a retrograde AAV containing Cre-dependent construct eYFP was bilaterally infused into the vHP of SERT-Cre mice. A Cre-dependent mCherry construct was infused locally into the MRR of the same mice. **h**, Representative coronal sections showing eYFP- and mCherry-labeled neurons in the MRR. 5-HT+ cells were detected using Tph2 immunostaining (not shown). Scale bars = 100 μm (overview), 20 μm (magnified). **i**, Percentage of mCherry-expressing Tph2+ neurons in the MRR (n = 2 mice). **j**, Left: Representative examples of mCherry+ (top) and eYFP- (bottom) cell visualized before patch-clamp recording. **k,** Examples of membrane potential responses to current steps (-30 pA for lower trace and + 190 pA for upper trace) of MRR 5-HT^non-vHP^ neurons (mCherry+, eYFP-) in female (grey) and male (brown) animals. **l-p**, Electrophysiological characterization of 5-HT^non-vHP^ neurons, with firing frequency in response to depolarizing current steps (**l**), firing frequency at +190 pA (**m**), resting membrane potential (**n**), input resistance (**o**), and adaptation index (**p**) for females (light teal, n = 12 cells from 5 mice) and males (light orange, n = 11 cells from 4 mice). All data are shown as mean ± s.e.m; **: *P* < 0.01, ***: *P* = < 0.001; For detailed statistical information, see Extended Data Table 1. Abbreviations: IP: interpeduncular nucleus, B9: B9 nucleus; MnR: median part of the median raphe region; PMnR: paramedian part of the median raphe region; MRR: median raphe region; DRI: dorsal raphe, interfascicular subregion; DR: dorsal raphe; DRC: caudal part of the dorsal raphe; Tph2: tryptophan hydroxylase type 2; eYFP: enhanced yellow fluorescent protein; For detailed statistical information, see Extended Data Table 1.

**Extended Data Fig. 2:**
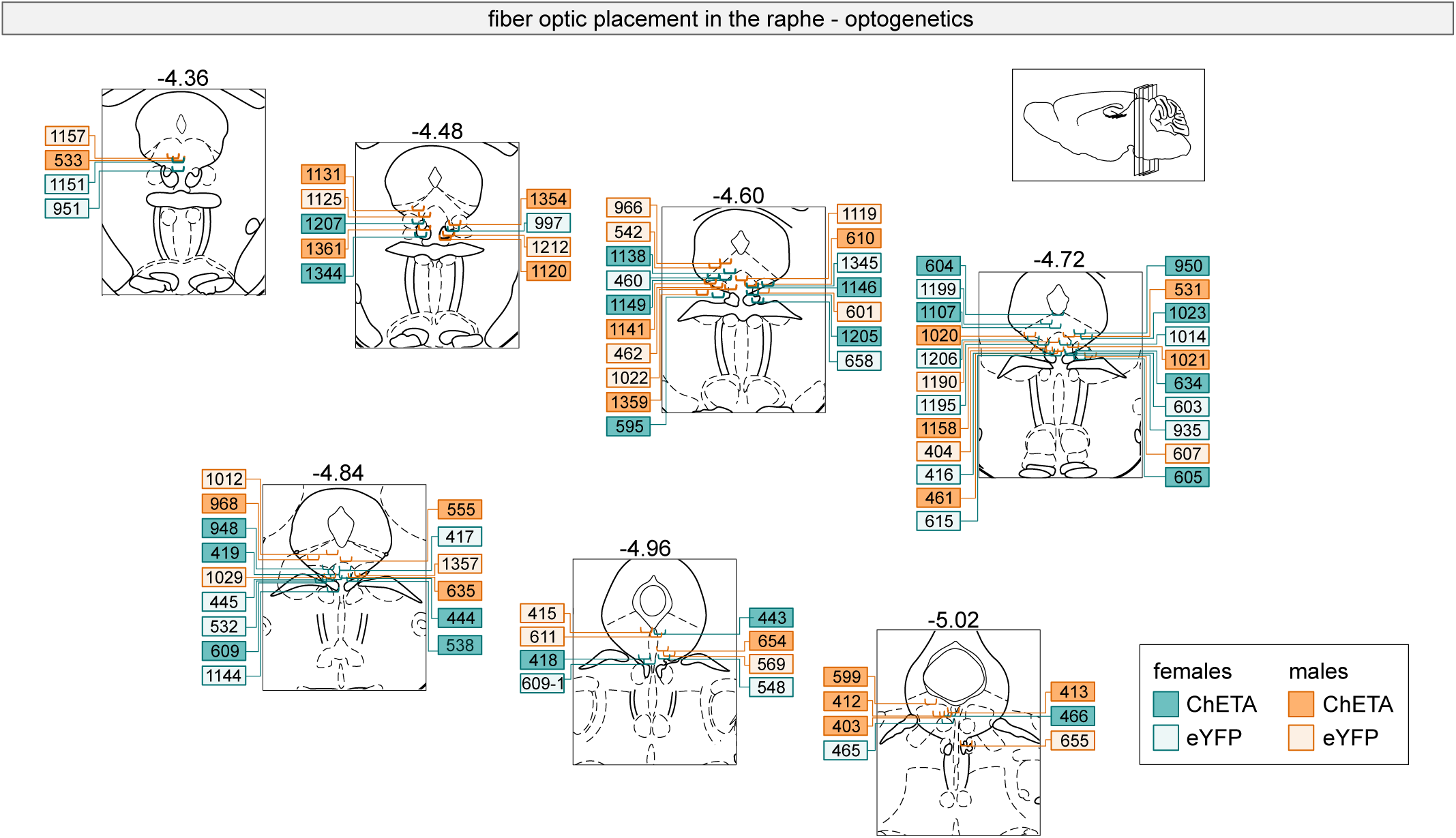
Optic fiber placements for optogenetic experiments. **a,** Fiber optic placement in raphe of experimental and control mice used for optogenetic stimulation during behavioral tests. Numbers above diagrams indicate distance from bregma in mm. Numbers in boxes indicate identification number of experimental subject.

**Extended Data Fig. 3.**
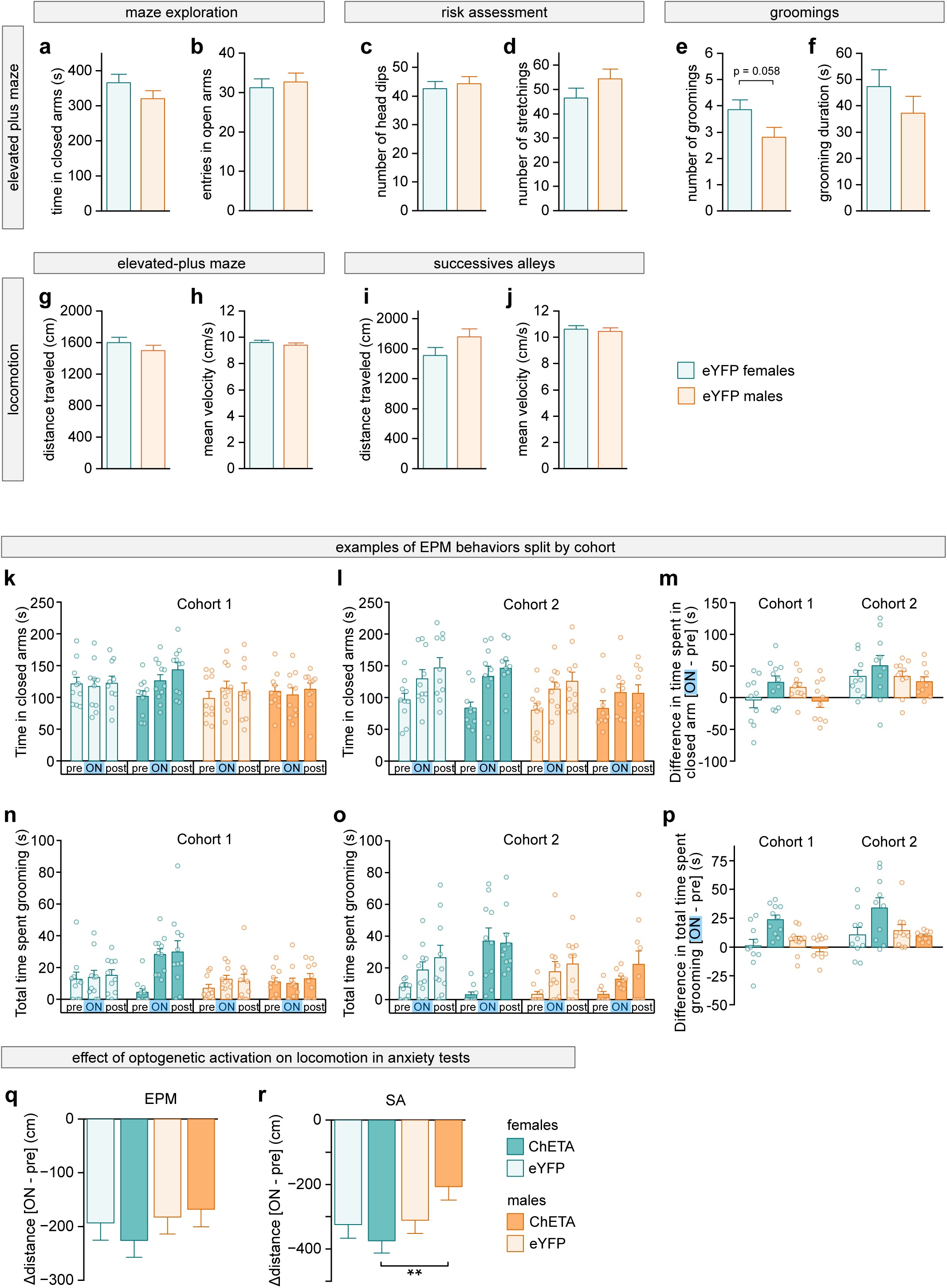
Baseline sex comparison of behavior and effect of 5-HT^vHP^ activation. **a**-**h**, Comparison of female (n = 20) and male (n = 20 eYFP) control mice behavior quantified across the complete 12-min EPM session for time spent in the closed arms (**a**), entries in the open arms (**b**), number of head dips (**c**), number of stretchings (**d**), number of groomings (**e**), grooming duration (**f**), total distance travelled (**g**) and mean velocity (h). **i**-**j**, Same as g-h, but across the complete successive-alleys session. **k-l,** Time spent in the closed arms during the pre-stimulation (pre), stimulation (stim) and post-stimulation (post) epochs for cohort 1 (**k**) and cohort 2 (**l**). **m,** Change in closed-arm time for both cohorts. **n-o,** Total time spent grooming during pre, stim and post epochs for cohort 1 (**n**) and cohort 2 (**o**). **p,** Change in grooming time for both cohorts. **q,** Difference between the distance travelled during the 4 min of optogenetic stimulation (ON) and the first 4 min of baseline (pre) periods in the elevated-plus maze for each group. No difference was observed across sexes or groups. **r,** Same as **q**, but in the successive alleys (SA). Three-way ANOVA with Holms multiple comparison testing, ChETA (female vs male), *P* = 0.0041. Data are shown as mean ± s.e.m. **: *P* < 0.01. Data are from two independent experiments and were analyzed by three-way ANOVA. For detailed statistical information and for the numbers of female and male mice used, see Extended Data Table 1. For an overview of all behavioral variables analyzed, see Extended Data Table 2.

**Extended Data Fig. 4:**
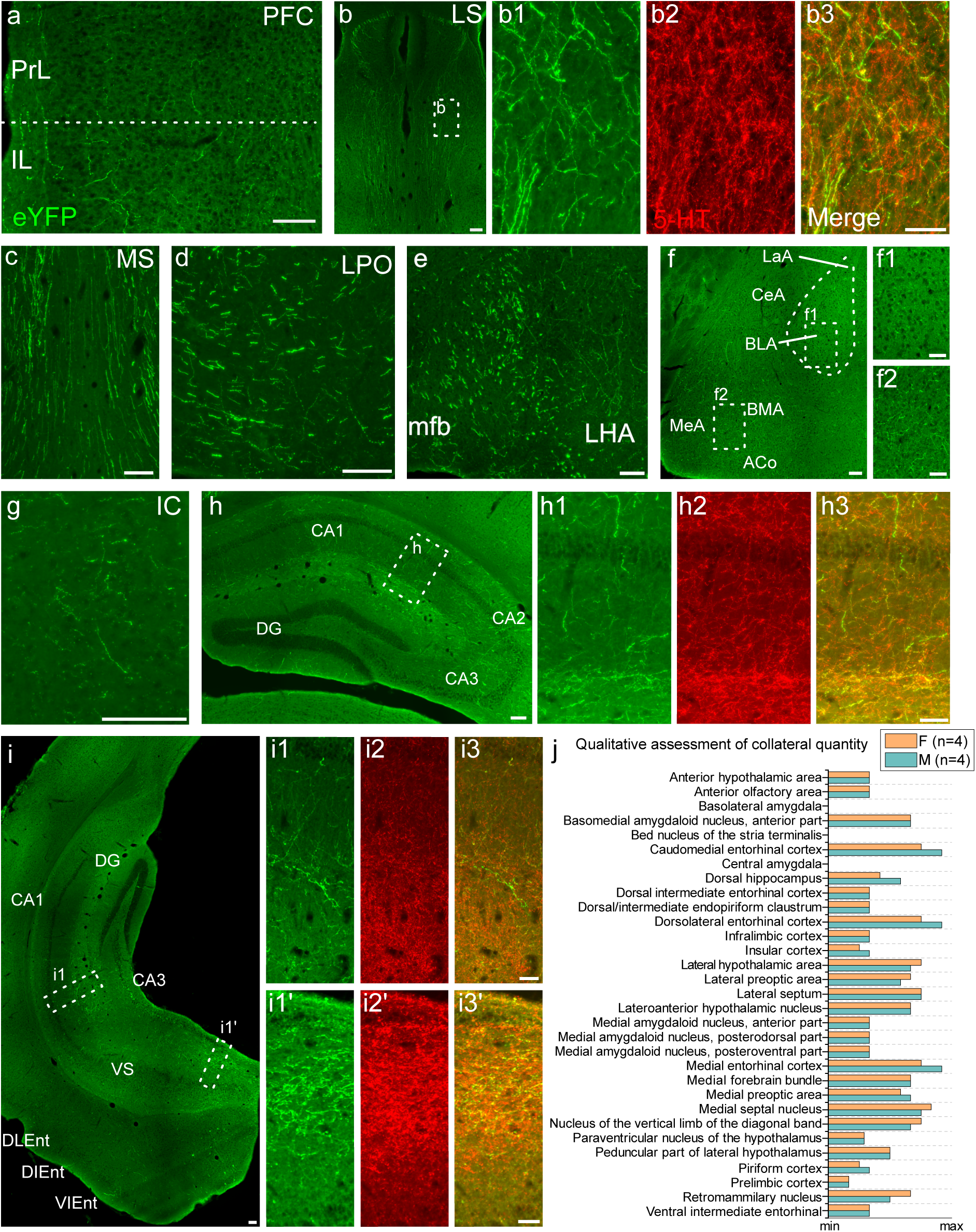
5-HT^vHP^ neuron axon collateralization to the forebrain. eYFP-positive axon collaterals are detected with anti-GFP immunostaining. **a,** Sparse axons in the prefrontal cortex (PFC). **b,** A high density of axons was observed in the lateral septum (LS). Overlay between eYFP fibers and 5-HT immunostaining (**b1**-**b3**). **c,** A high density of axon collaterals is found lateral to the medial septum (MS). **d,** Dense axon collaterals are found in the preoptic area with the highest density in the lateral part (LPO). **e,** Axon collaterals can be seen passing through the medial forebrain bundle (mfb) with some also seen in the lateral hypothalamic area (LHA). **f,** Medial amygdaloid nucleus (MeA), basomedial amygdaloid nucleus (BMA) and anterior cortical amygdaloid area (ACo) all contain a high density of eYFP-positive axons while very few to none are observed in the central amygdala (CeA), lateral amygdala (LaA) and the basolateral amygdala (BLA). **g,** Sparse axons are consistently present in the insular cortex (IC). **h,** 5-HT^vHP^ neurons send axon collaterals to the dHP. **i,** Example of the injection site in the vHP with eYFP-positive axons covering the dentate gyrus (DG), CA1, CA3 and VS (ventral subiculum). High density of eYFP labeled axons are present in CA1 (**i1-i3**) and CA3 (**i1’-i3’**). Axon collaterals are also observed in the dorsolateral, dorsal intermediate and ventral intermediate part of the entorhinal cortex (DLEnt, DIEnt and VIEnt, respectively) but leakage from the injection site should be considered. **j,** Qualitative assessment of 5-HT collaterals density in female (n = 4) and male (n = 4) mice. ACo: anterior cortical amygdaloid area; BLA: basolateral amygdala; BMA: basomedial amygdaloid nucleus; CeA: central amygdala; DG: dentate gyrus; DIEnt: dorsal intermediate enthorinal cortex; DLEnt: dorsolateral entorhinal cortex; eYFP: enhanced yellow fluorescent protein; IC: insular cortex; LaA: lateral amygdala; LHA: lateral hypothalamic area; LPO: lateral part of preoptic area; LS: lateral septum; MeA: medial amygdaloid nucleus; mfb: medial forebrain bundle; MS: medial septum; PFC: prefrontal cortex; VIEnt: ventral intermediate part of the entorhinal cortex; VS: ventral subiculum; Scale bar: 100 μm in A-I, 50 μm in b1-b3; f1-f2; h1-h3; i1-i3; i1’-i3’.

**Extended Data Fig. 5:**
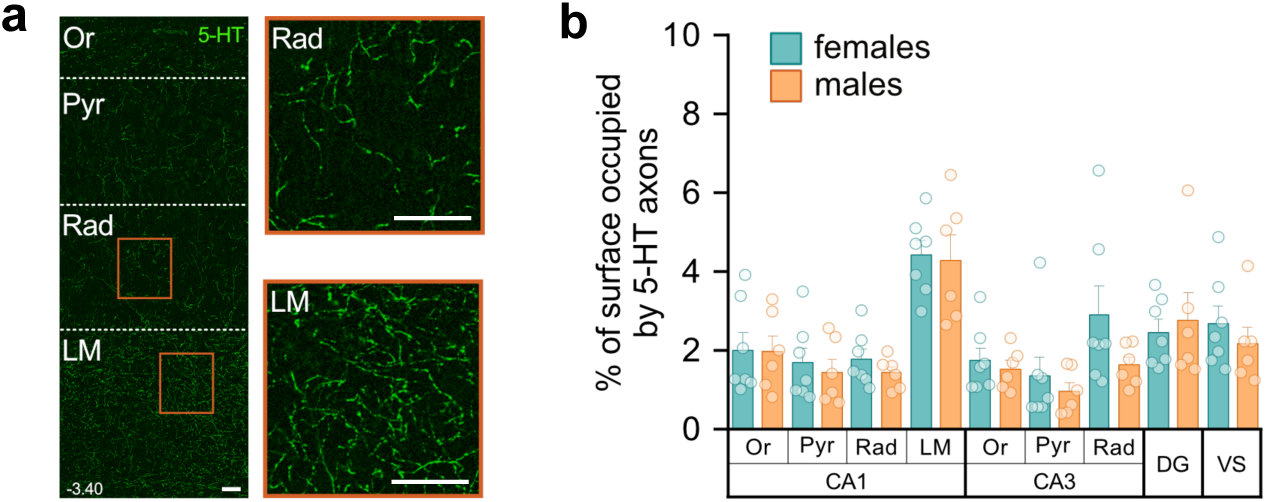
Male and female mice display comparable 5-HT^vHP^ axon density in the vHP. **a**, Serotonergic axons are detected using 5-HT immunostaining and are visible in the stratum oriens (Or), pyramidale (Pyr), radiatum (Rad) and stratum lacunosum-moleculare (LM). Scale bars = 30 μm. **b**, Axon density quantified as the % of surface occupied by 5-HT-positive axons in the regions of interest across vHP layers in females (n = 7) and male (n = 6) mice. All data are shown as mean ± s.e.m. For detailed statistical information, see Extended Data Table 1. DG: Dentate gyrus; LM: stratum lacunosum-moleculare; Or: stratum oriens; Pyr: stratum pyramidale; Rad: stratum radiatum; VS: ventral subiculum.

**Extended Data Fig. 6:**
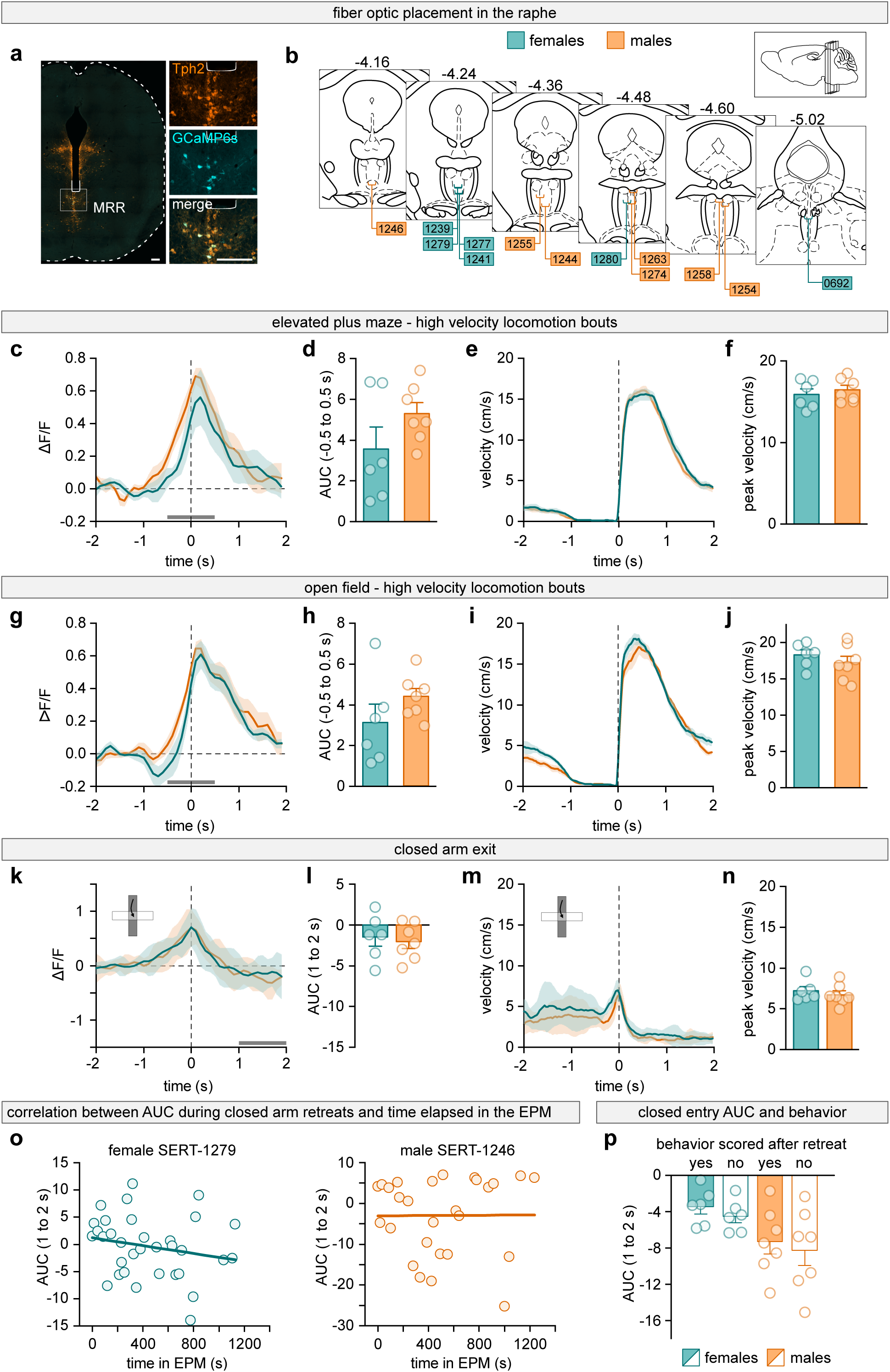
Locomotion-related activity of 5-HT^vHP^ neurons does not differ between sexes. **a,** Example fiber placement above the MRR, with magnified view of cell bodies (right). 5-HT neurons (Tph2+) are shown in orange; GCaMP6s-labelled 5-HT^vHP^ neurons shown in cyan. Scale bars = 200 μm. **b,** Optic fiber placements for all animals used for fiber photometry experiments (n = 6 females, n = 7 males). Numbers in boxes indicate individual animal IDs; brain schematics show distance from bregma in mm. **c,** Mean normalized GCaMP6s fluorescence signal aligned to the initiation of high velocity locomotion bouts for females (teal, n = 6) and males (orange, n = 7) in the EPM. Dotted vertical line: onset of locomotion. Gray bar indicates the period used for panel **d**. **d,** AUC of ΔF/F from –0.5 to +0.5 s around high velocity locomotion onset in the EPM. **e,** Corresponding instantaneous velocity across all detected high velocity locomotion bouts. Dotted vertical line indicates onset of locomotor bout. **f,** Quantification of peak velocity for each animal in **e. g,** Same as **c,** but for the open field. Gray bar indicates the period used for panel **h**. **h,** Same as **d,** but for the open field. **i,** Same as **e,** but for the open field. **j,** Same as **f,** but for the open field. **k,** Mean normalized ΔF/F for closed arm exits towards center/open zones for females (teal, n = 6) and males (orange, n = 7). Dotted vertical line indicates timepoint where center-of-body marker crossed the center-closed arm limit. Gray bar indicates the period used for panel **l**. **l,** AUC of ΔF/F obtained over 1 - 2 s after closed arm entry (gray bar in **k**). Two-sample t-test, *P* = 0.0430. **m,** Velocity around closed arm exits for females (teal, n = 6) and males (orange, n = 7). **n,** Peak velocity quantification (-0.5 to 0.5 s) for females (teal, n = 6) and males (orange, n = 7). **o,** Example correlations between AUC of ΔF/F obtained over 1 - 2 s after each closed arm entry and time spent in the EPM for a representative female (left) and for a representative male (right). **p**, AUC of ΔF/F from 1-2 s after closed-arm entry, calculated before and after excluding entries containing any other manually scored behavior during the peri-entry analysis window. Behavioral exclusion did not significantly effect AUC values (two-way ANOVA, main effect of exclusion, P = 0.43; females, n = 6; males, n = 7). **c** – **n**, all data are shown as mean ± s.e.m. For detailed statistical information, see Extended Data Table 1.

**Extended Data Fig. 7:**
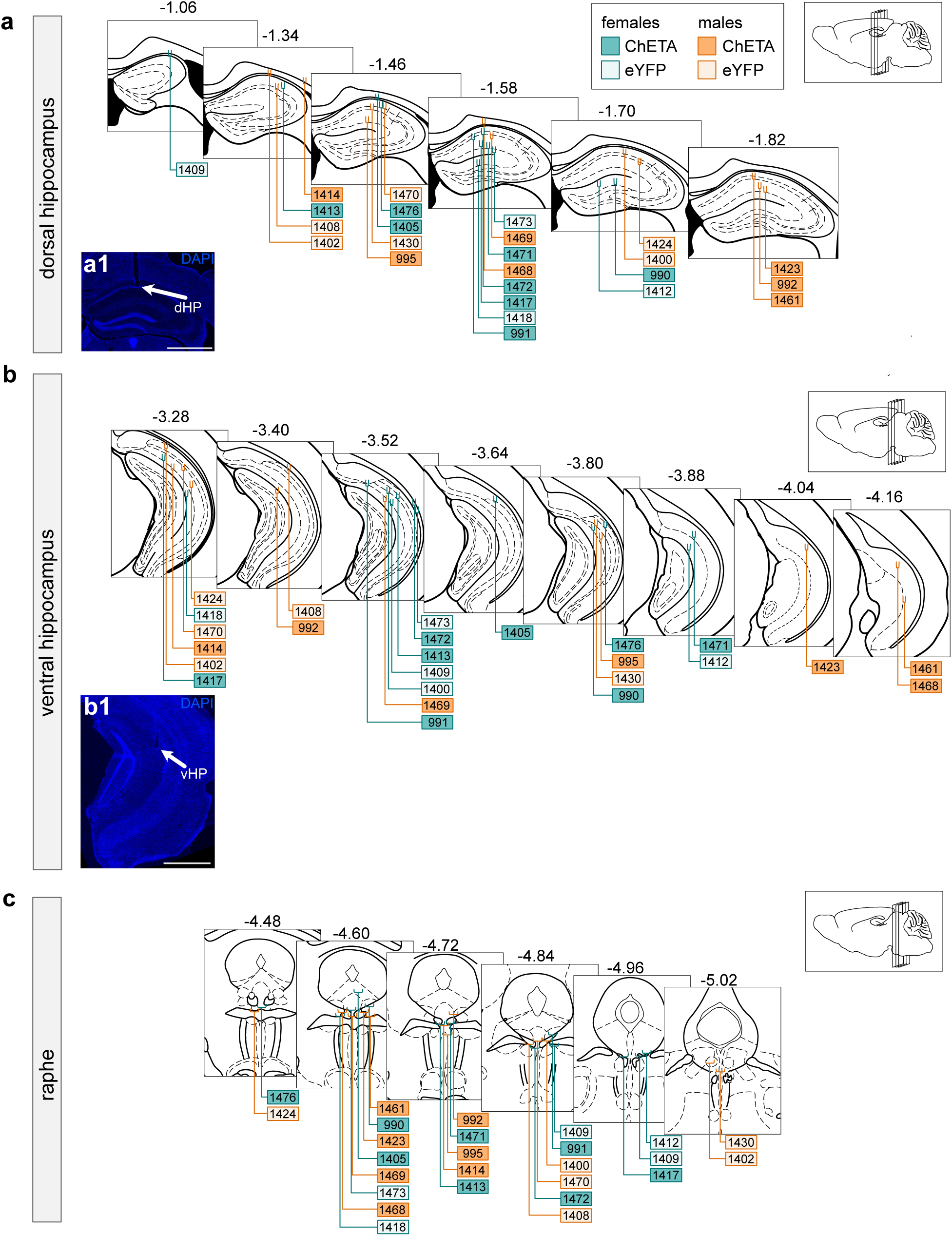
Electrode and optic fiber placements for LFP recordings and optogenetic stimulation. **a,** Electrode placement in the dHP, with representative electrode tract (**a1**, DAPI in blue; as indicated by white arrow). Scale bar = 1000 μm. Numbers in boxes indicate ID of the animals. Numbers above diagrams indicate distance from bregma in mm. **b,** Same as a, for the vHP (**b1**: representative tract as indicated by white arrow). Scale bar = 1000 μm. **c,** Fiber optic placement in the raphe.

**Extended Data Fig. 8:**
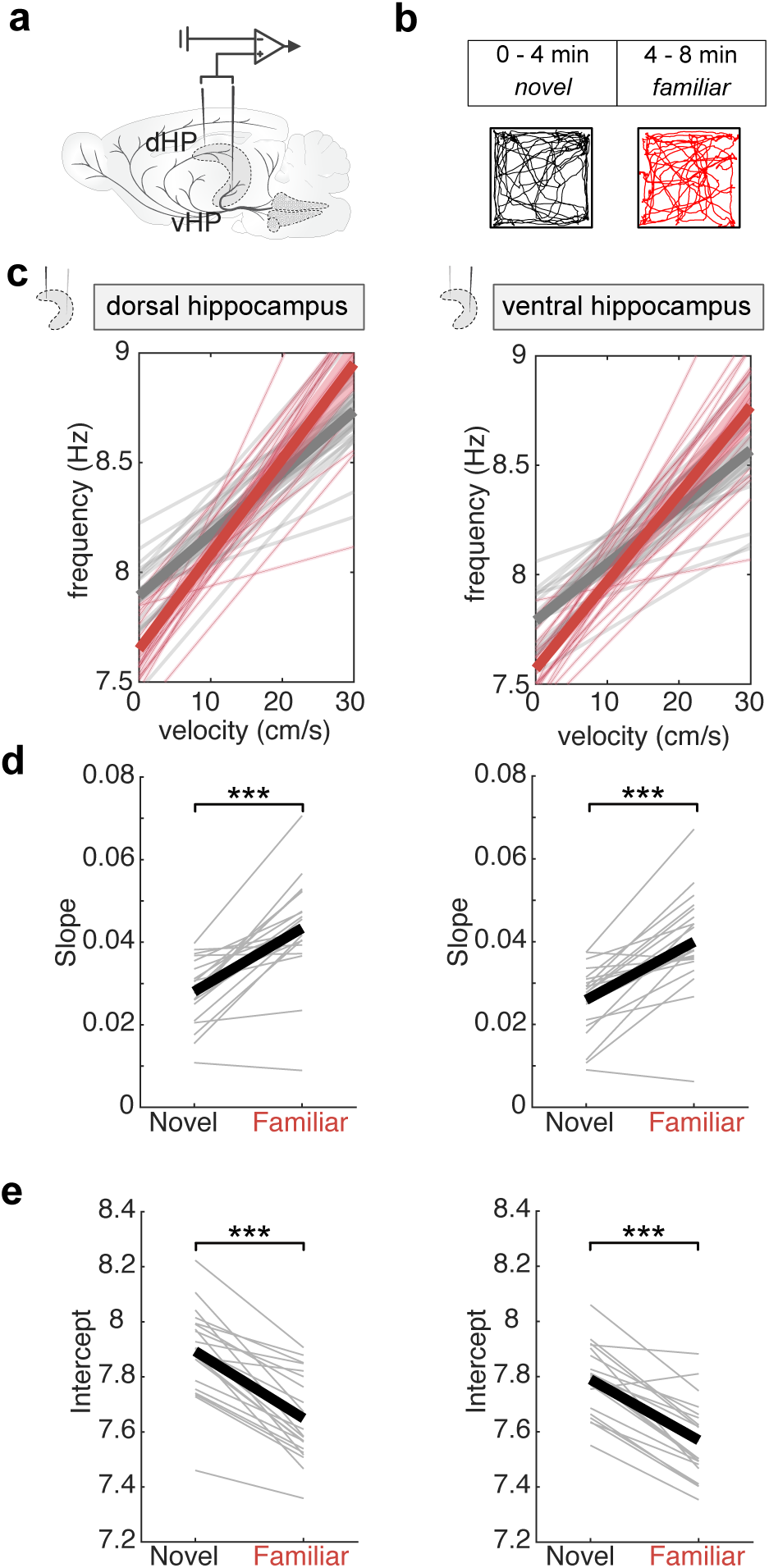
Theta-velocity relationship over familiarization to a novel environment. **a,** Schematic of simultaneous multi-site local field potential recording. **b,** Experimental paradigm and example trajectories for one animal for novel (0-4 min, grey) and familiar (4-8 min, red) periods. **c,** Instantaneous theta frequency as a function of running speed for the dHP (left) and vHP (right) for novel (grey) and familiar (red) period, n = 21 mice. Thin lines: individual animals; thick lines: group mean. Shaded areas represent s.e.m. **d,** Slope for novel and familiar epochs. For dHP (left): Paired sample t-test, *P* < 0.0001; for vHP (right): Paired sample t-test, *P* < 0.0001. **e**, Intercept for novel and familiar epochs. For dHP (left): Paired sample t-test: P < 0.0001; for vHP (right): Paired sample t-test: *P* < 0.0001. All data are shown as mean ± s.e.m; ***: *P* < 0.0001; dHP: dorsal hippocampus; vHP: ventral hippocampus. For detailed statistical information, see Extended Data Table 1.

**Extended Data Fig. 9:**
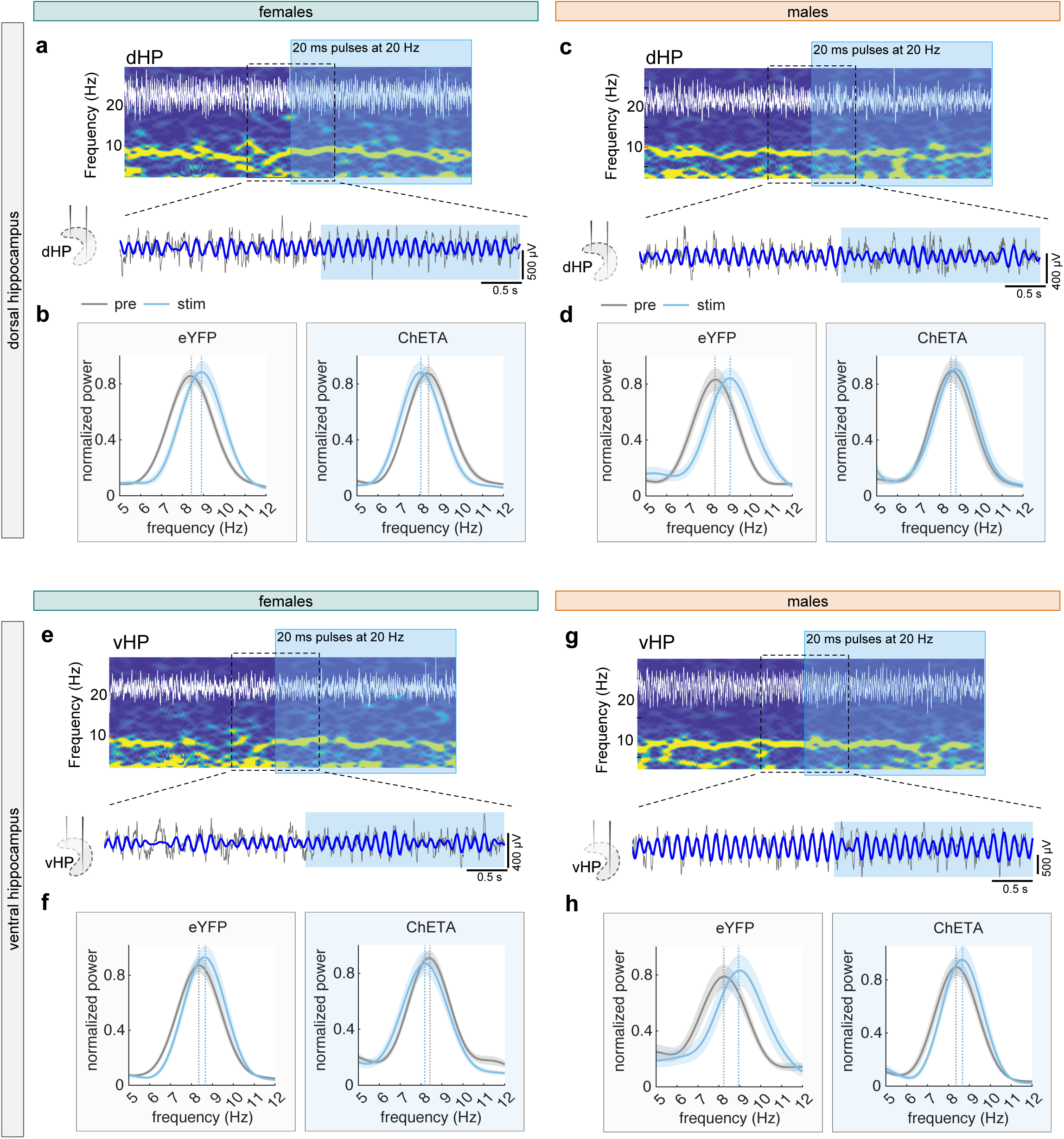
Effects of 5-HT^vHP^ neuron activation on hippocampal LFP activity. **a,** Representative spectrogram and corresponding raw trace (white) showing hippocampal LFP activity during the 10 s preceding and 10 s following the onset of 20 Hz photostimulation (blue box) in a ChETA female dHP. Bottom traces show the corresponding raw (grey) and theta filtered (blue) LFP signals during the 2.5 s before and 2.5 s after stimulation onset, as indicated by the blue box. **b**, Same as b, for vHP. **c**, Normalized power spectra for dHP during active locomotion (>10 cm/s) for pre (grey) and stim (blue) epochs for eYFP-control female (left) and ChETA female (right) **a**. Shaded areas represent s.e.m. Vertical lines indicate theta peak frequency. **d**, Same as c, for vHP. **e**-**f**, Same as a-b, for ChETA male. **g**-**h**, Same as c-d, for ChETA male.

**Extended Data Fig. 10:**
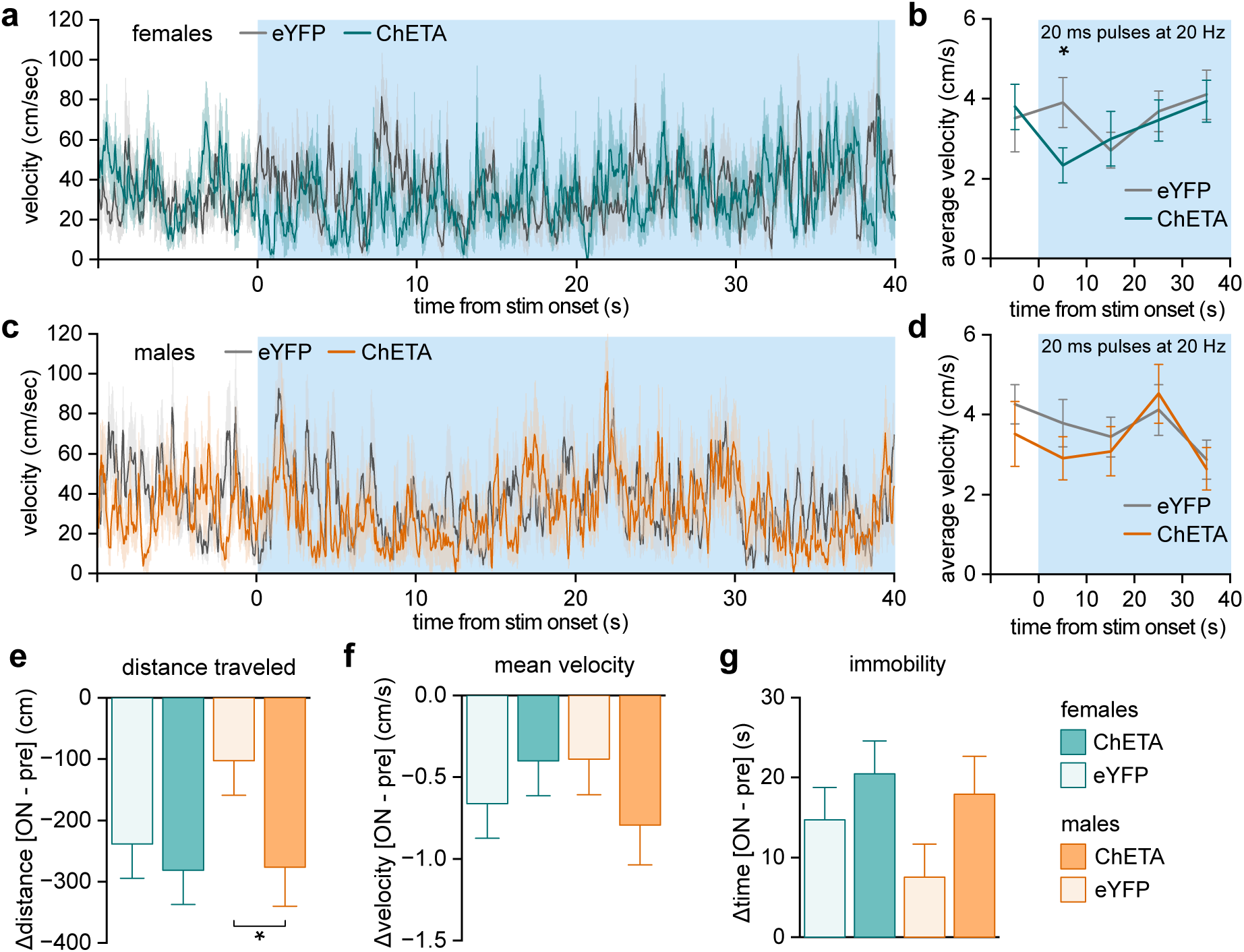
Optogenetic activation of 5-HT^vHP^ neurons does not induce major changes in locomotion. **a,** Instantaneous velocity over time, around light stimulation onset (time = 0) for female eYFP (gray) and ChETA (teal) mice in the open field. **b,** Velocity averaged by 10 s bouts, around light stimulation onset (time = 0) for female eYFP (gray) and ChETA (teal) mice in the open field. Time-binned analyses of average velocity confirmed the absence of prominent stimulation effects, aside from a transient (10 s) decrease in average velocity. Two-sample t-test, eYFP vs control, 0 - 10 s, *P* = 0.0461. Blue box indicates optogenetic stimulation onset. **c,** Same as **a**, but for male eYFP (gray) and ChETA (orange) mice. **d,** Same as **b**, but for male eYFP (gray) and ChETA (orange) mice. No change in velocity was observed in males. **e,** Difference between the distance travelled during the 4 min of optogenetic stimulation (ON) and the first 4 min of baseline (pre) periods in the open field for each group. No difference was observed for females while ChETA-expressing males showed a modest reduction in distance traveled. Three-way ANOVA with Holms multiple comparison testing: males (ChETA vs control), *P* = 0.0452). **f,** Same as **e**, but for mean velocity, no difference was observed for either sex. **g,** Same as **e**, but for time spent immobile. Here again, no difference was observed for either sex. For **e-g** data are from two independent experiments and were analyzed by three-way ANOVA. For detailed statistical information and for the numbers of female and male mice used, see Extended Data Table 1. For an overview of all behavioral variables analyzed, see Extended Data Table 2.

**Extended Data Fig. 11:**
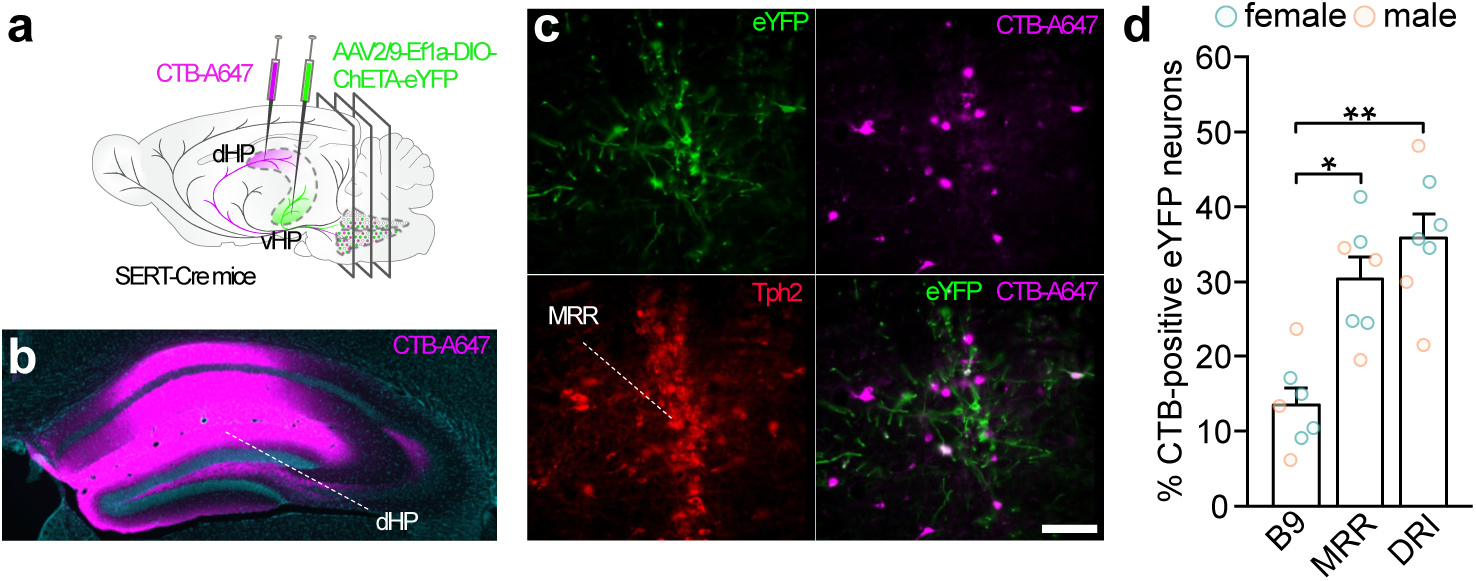
A subpopulation of 5-HT^vHP^ neurons collateralizes to dHP. **a,** Experimental strategy for dual retrograde labeling of 5-HT^vHP^ neurons projecting to both the dHP and the vHP. **b,** Representative image of CTB-A647 (magenta) in the dHP. **c,** Representative image of triple-labeled 5-HT neurons (Tph2+, red), expressing eYFP (eYFP+, green) and CTB-A647 (CTB+, magenta) in the MRR. Scale bar = 100 μm. **d,** Proportion of double-labeled 5-HT^vHP^ neurons across raphe subregions for females (teal, n = 4) and males (orange, n = 3): 13.39% in B9, 32.83% in the MRR and ≥ 35.71% in the DR. Kruskall-Wallis ANOVA, with Dunn’s post-hoc testing. B9 vs MRR: *P* = 0.0258, B9 vs DRI: *P* = 0.0017. Data are shown as mean ± s.e.m. * : *P* < 0.05, ** : *P* < 0.01. CTB: Cholera Toxin Subunit B; dHP: dorsal hippocampus; DRI: dorsal raphe, interfascicular subregion; eYFP: enhanced yellow fluorescent protein; MRR: median raphe region; For detailed statistical information, see Extended Data Table 1.

**Extended Data Fig. 12:**
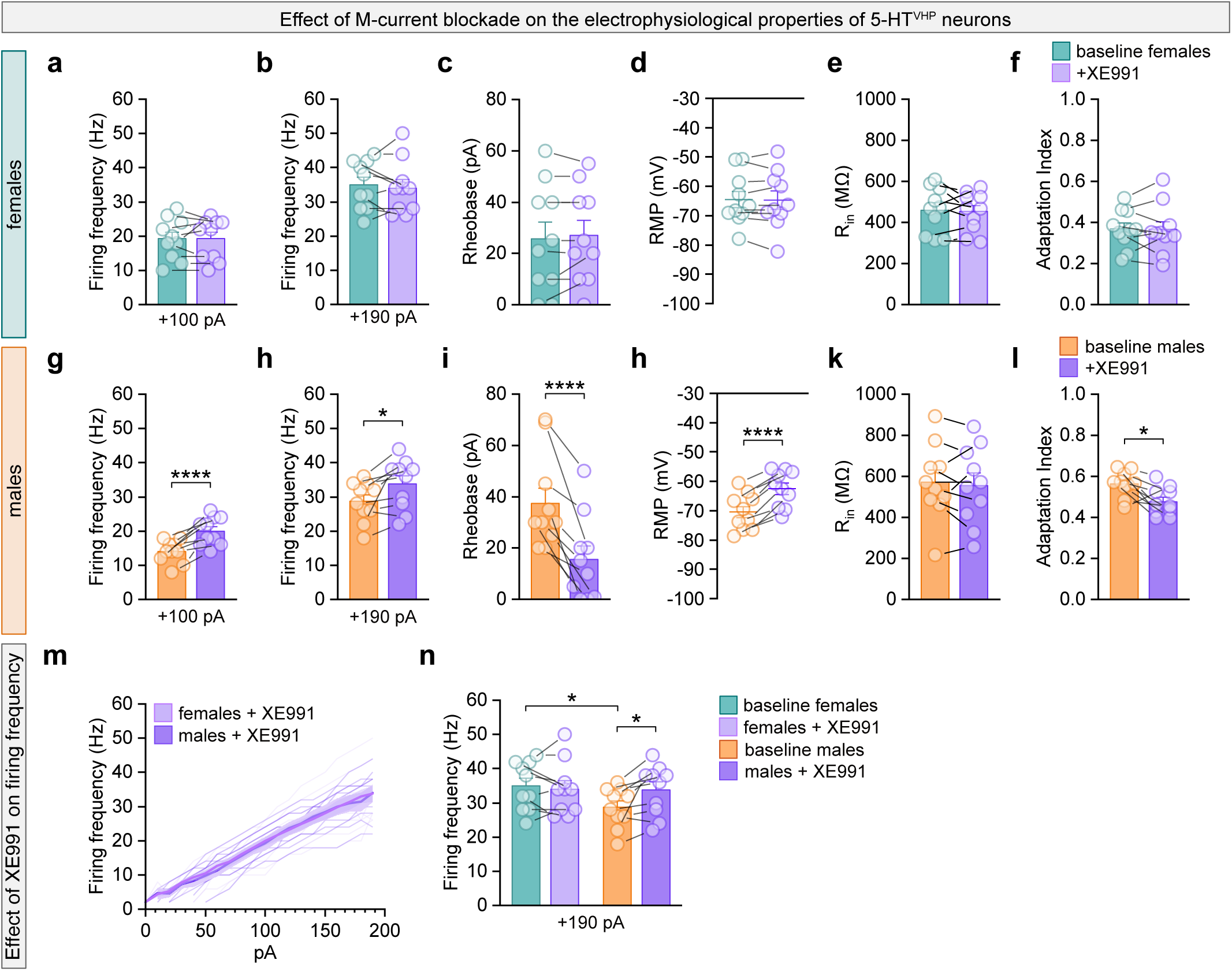
Effect of M-current blockade on the electrophysiological properties of females and males MRR 5-HT^vHP^ neurons. **a - k,** For female neurons, electrophysiological properties in response to application of the M-current blocker XE991. Data is the same as in Figure 7f-k but with baseline (teal) and M-current blocker XE911 (light purple) shown separately. **a**, Firing frequency at +100 pA. **b**, Firing frequency at +190 pA. **c,** Rheobase. **d**, Resting membrane potential. **e,** Input resistance. **f**, Adaptation index . **g- l,** Same as a-k for male neurons. **g,** Firing frequency at +100 pA, paired sample t-test *P* < 0.0001. **h,** Firing frequency at +190 pA, paired sample t-test *P* = 0.0106. **i,** Rheobase, paired sample t-test *P* = 3.6663e-4. **j,** Resting membrane potential, paired sample t-test *P* < 0.0001. **l,** Input resistance, paired sample t-test *P* = 0.0139. **m,** Firing frequency for male and female neurons as a function of depolarizing current stimulation during bath application of XE991; data displayed is the same as in Figure 7 d and e but highlights the overlap between male and female responses. **n,** Firing frequency at + 190 pA for female and male neurons at baseline (females = teal, males = orange) and with XE991 (females = light purple, males = dark purple); data displayed is the same as in Figure 7g. Baseline females vs. baseline males two-sample t-test *P* = 0.0463. Females: n = 10 cells from 7 mice, males: n = 10 cells from 9 mice. For detailed statistical information, see Extended Data Table 1.

**Extended Data Table 1:**
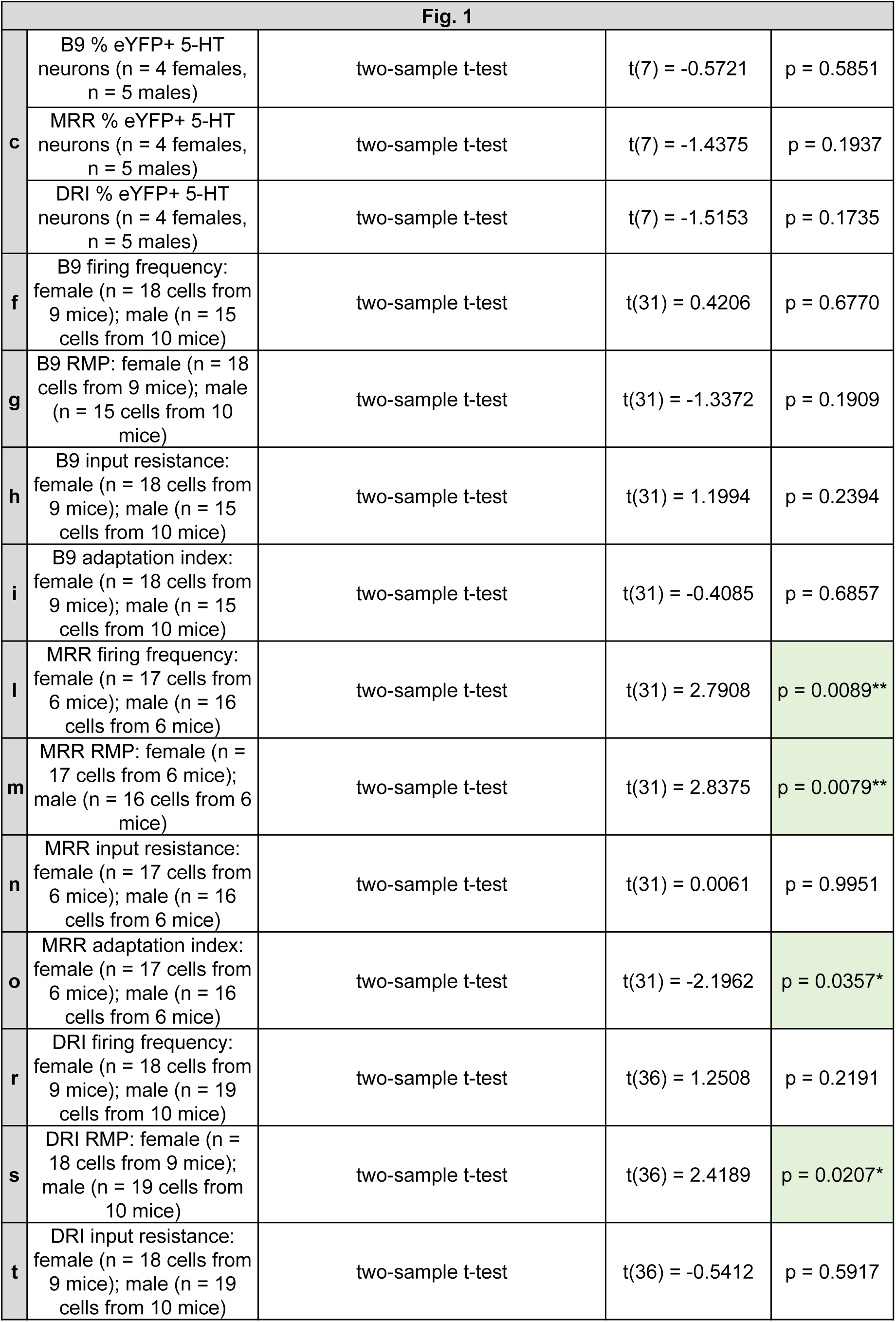
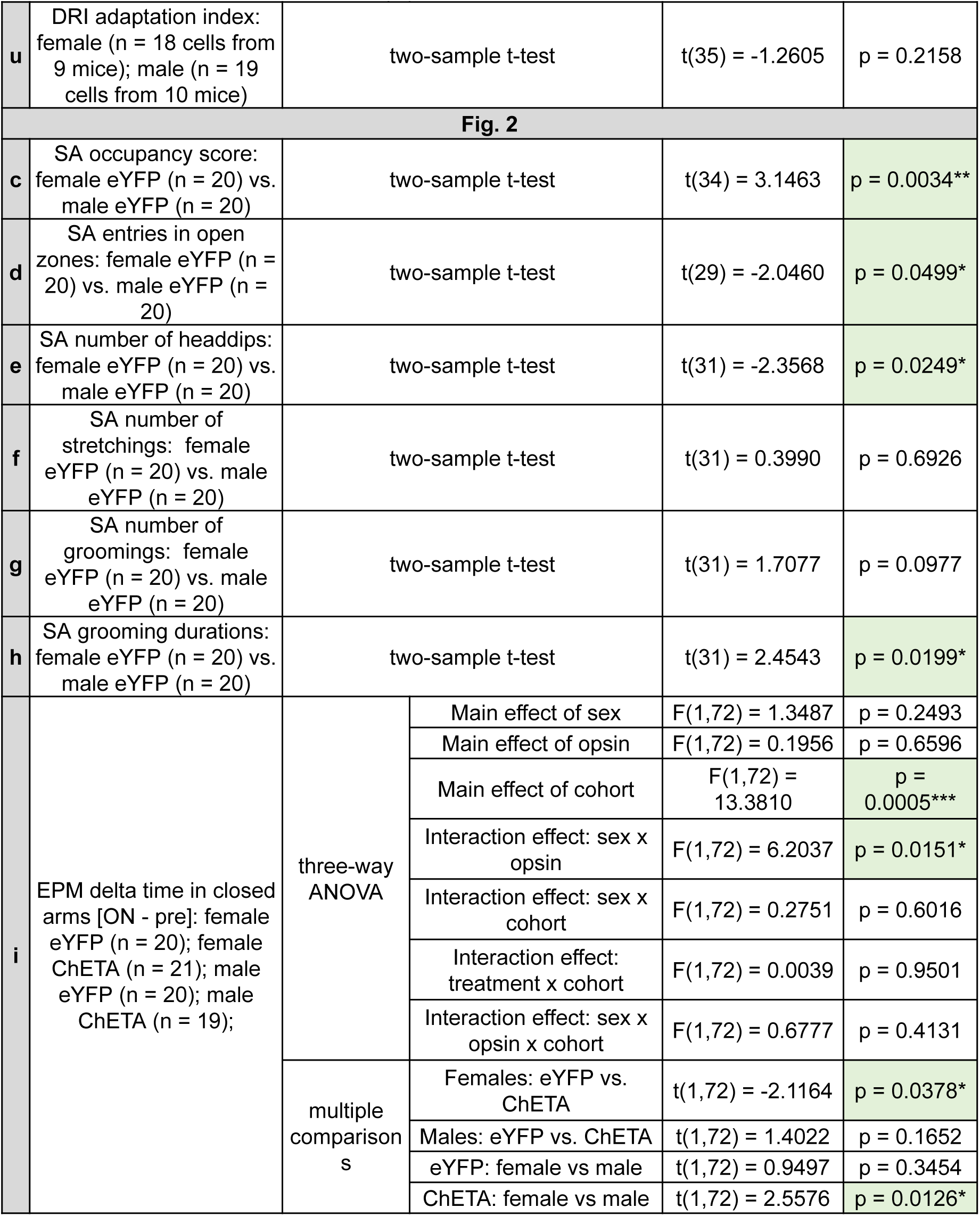
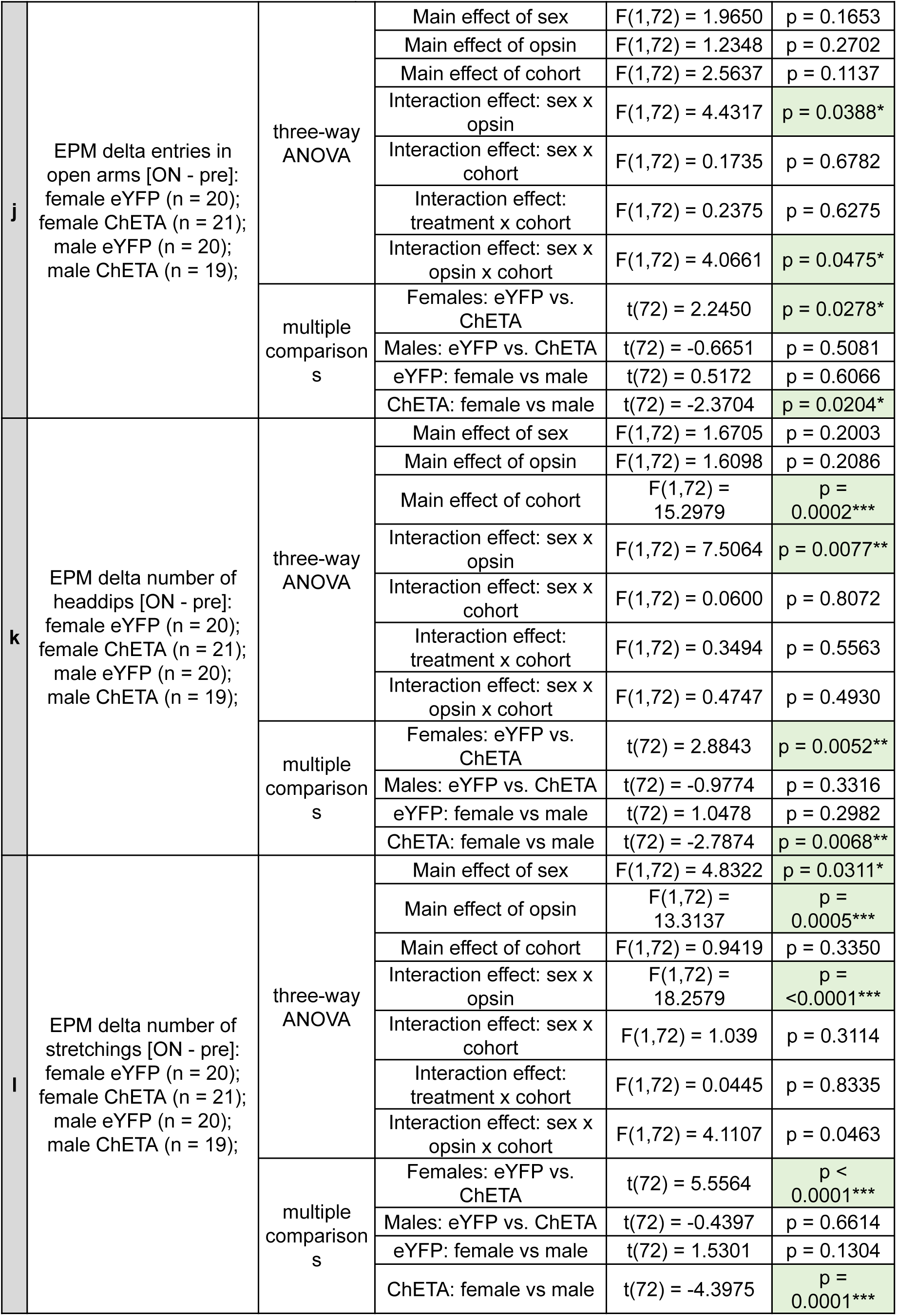
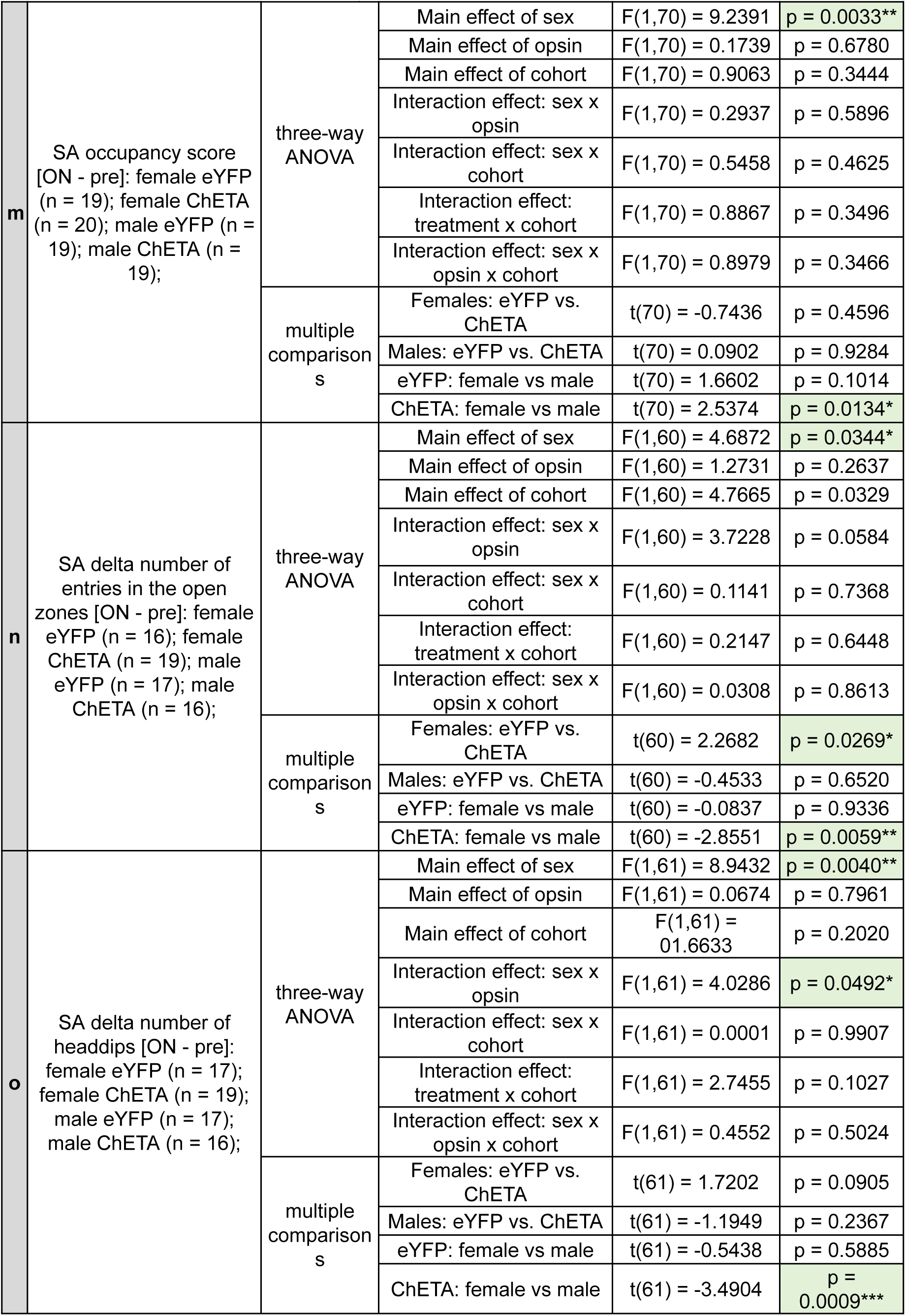
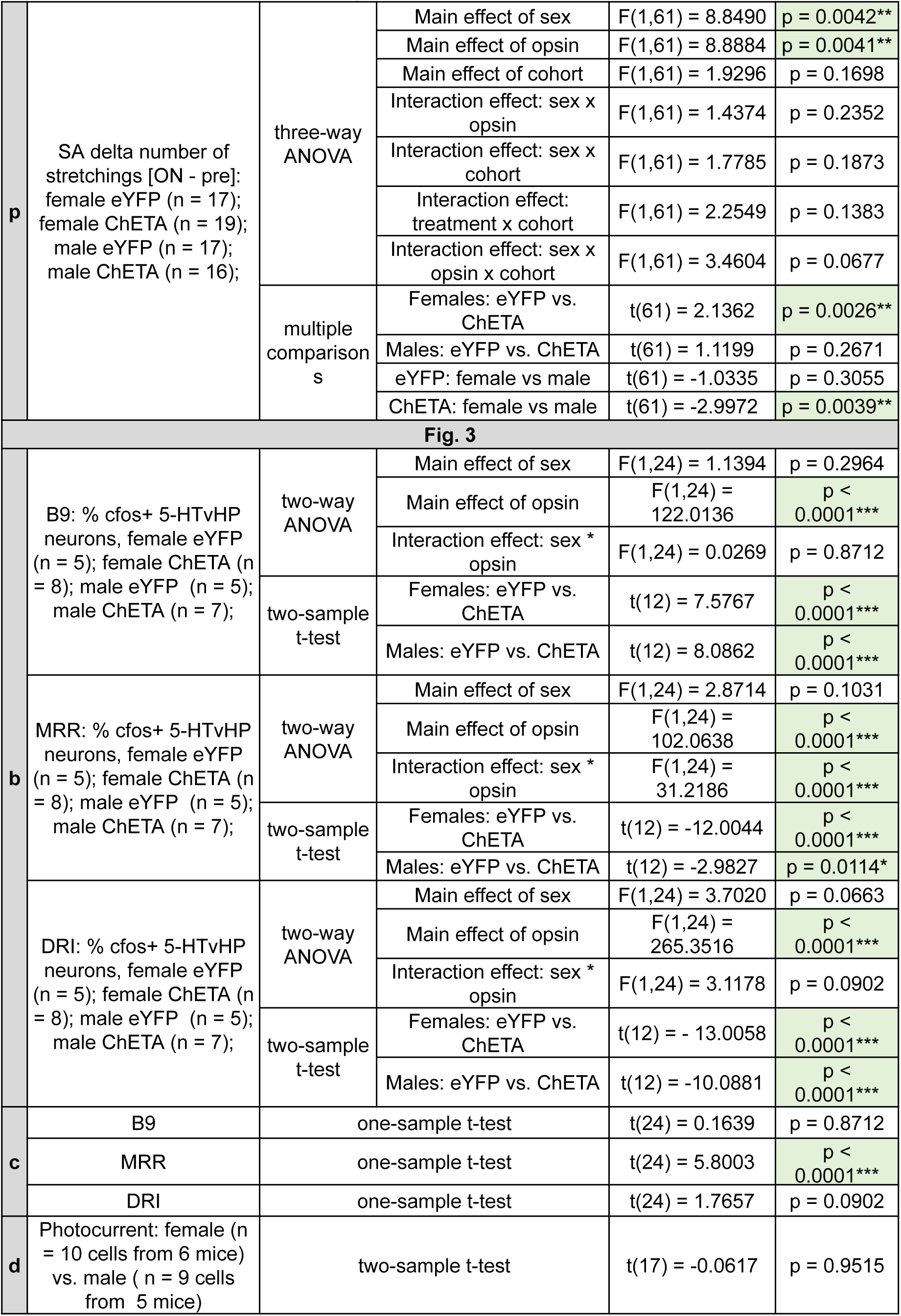
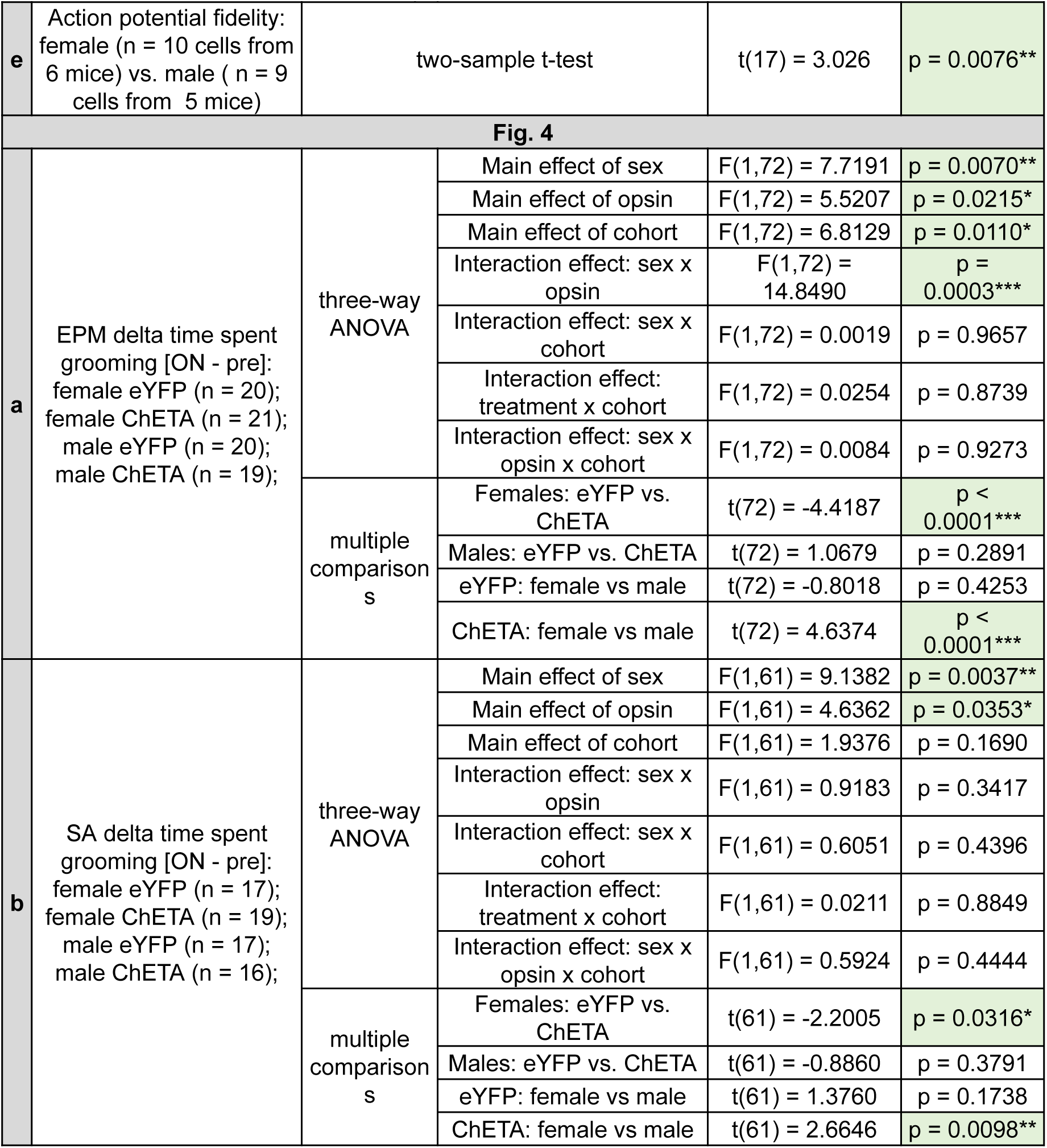
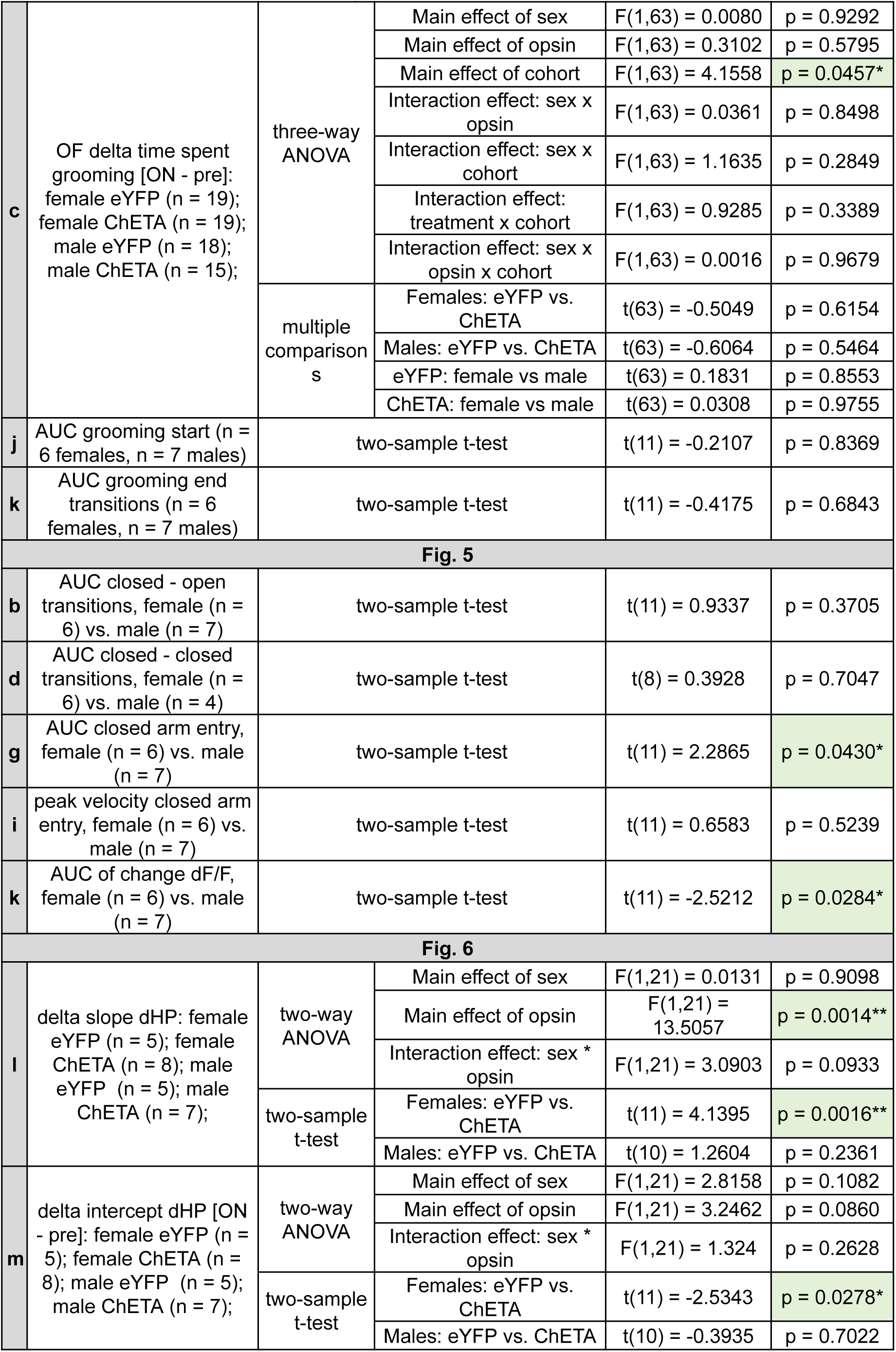
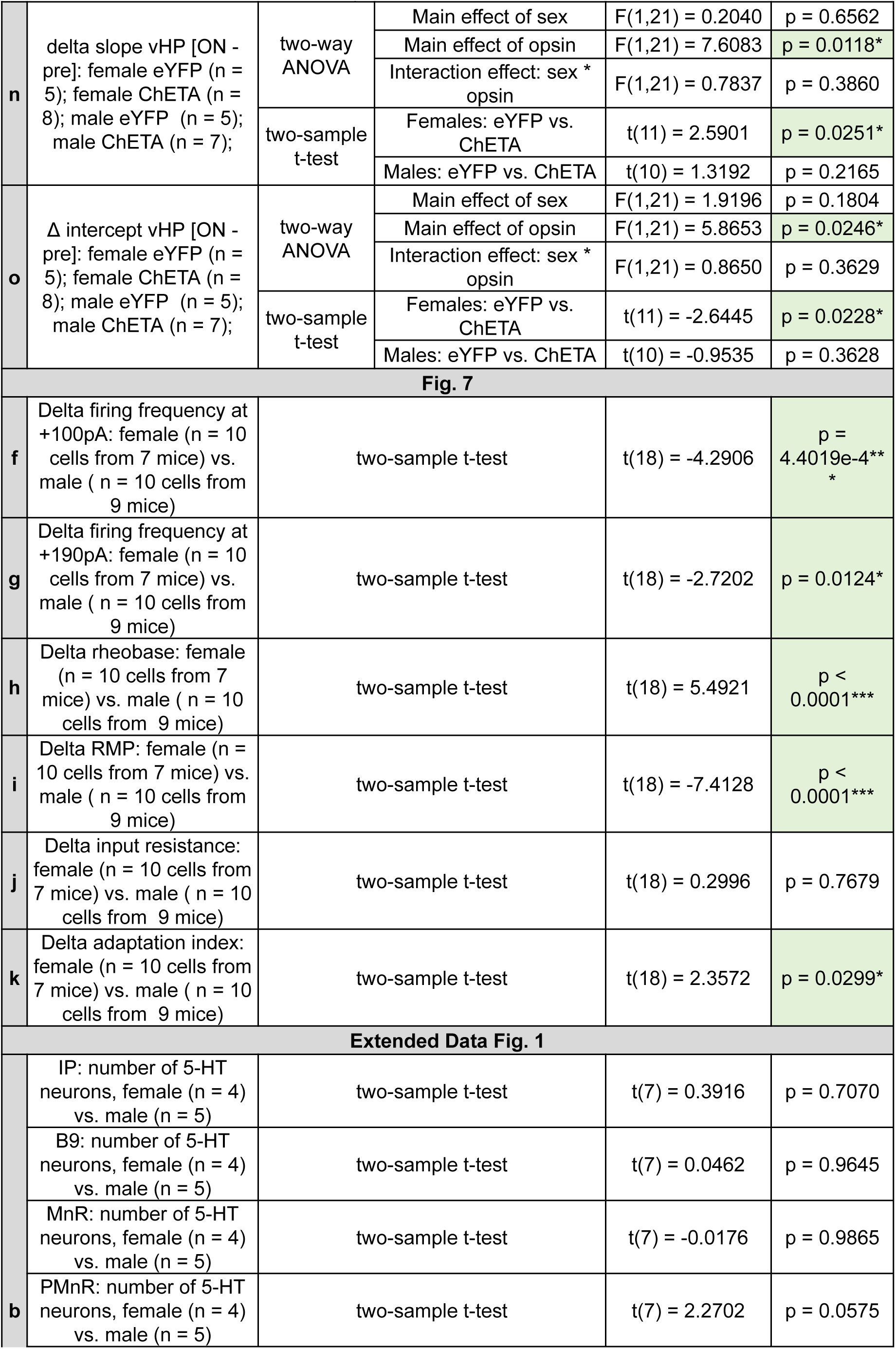
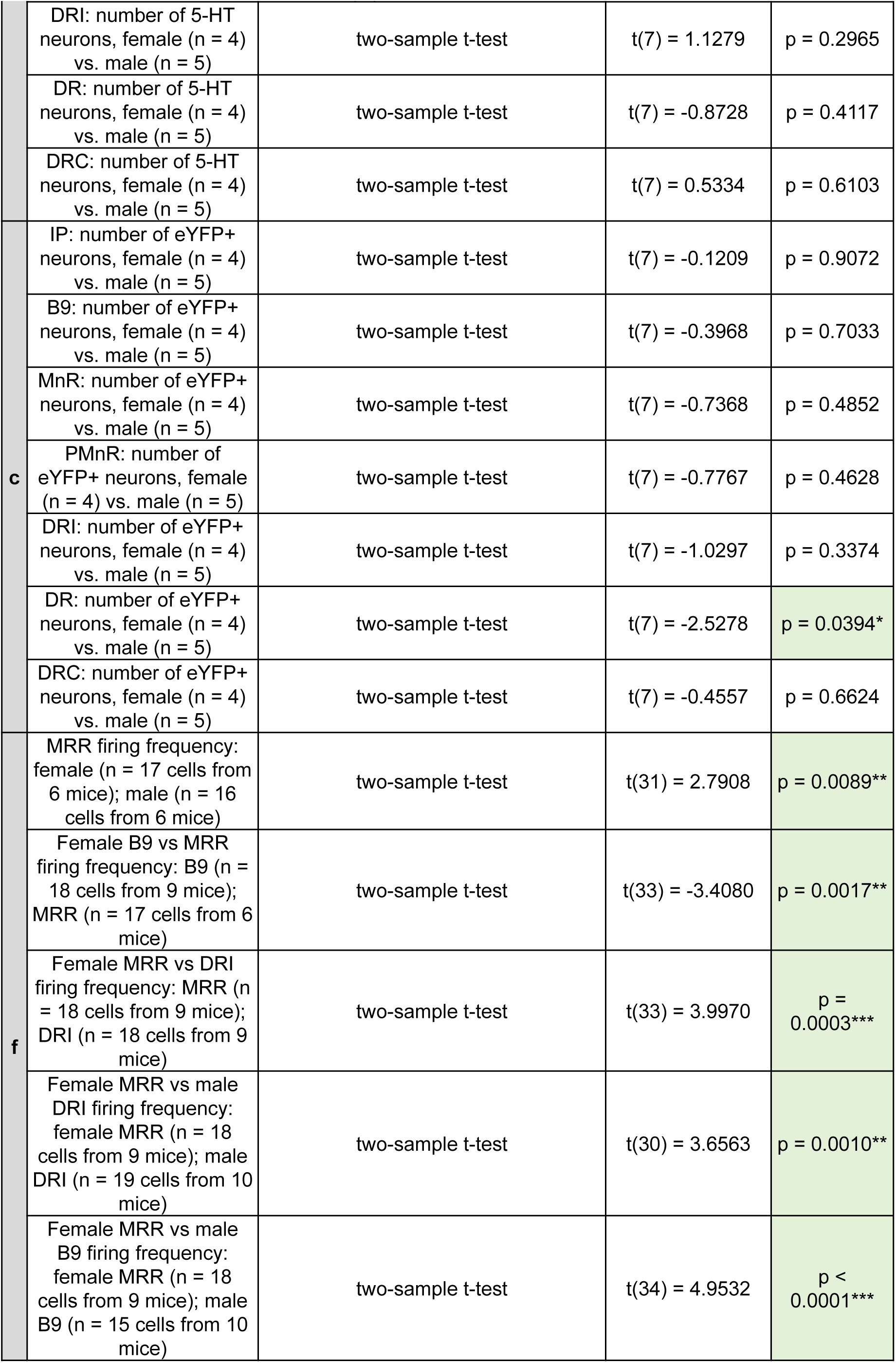
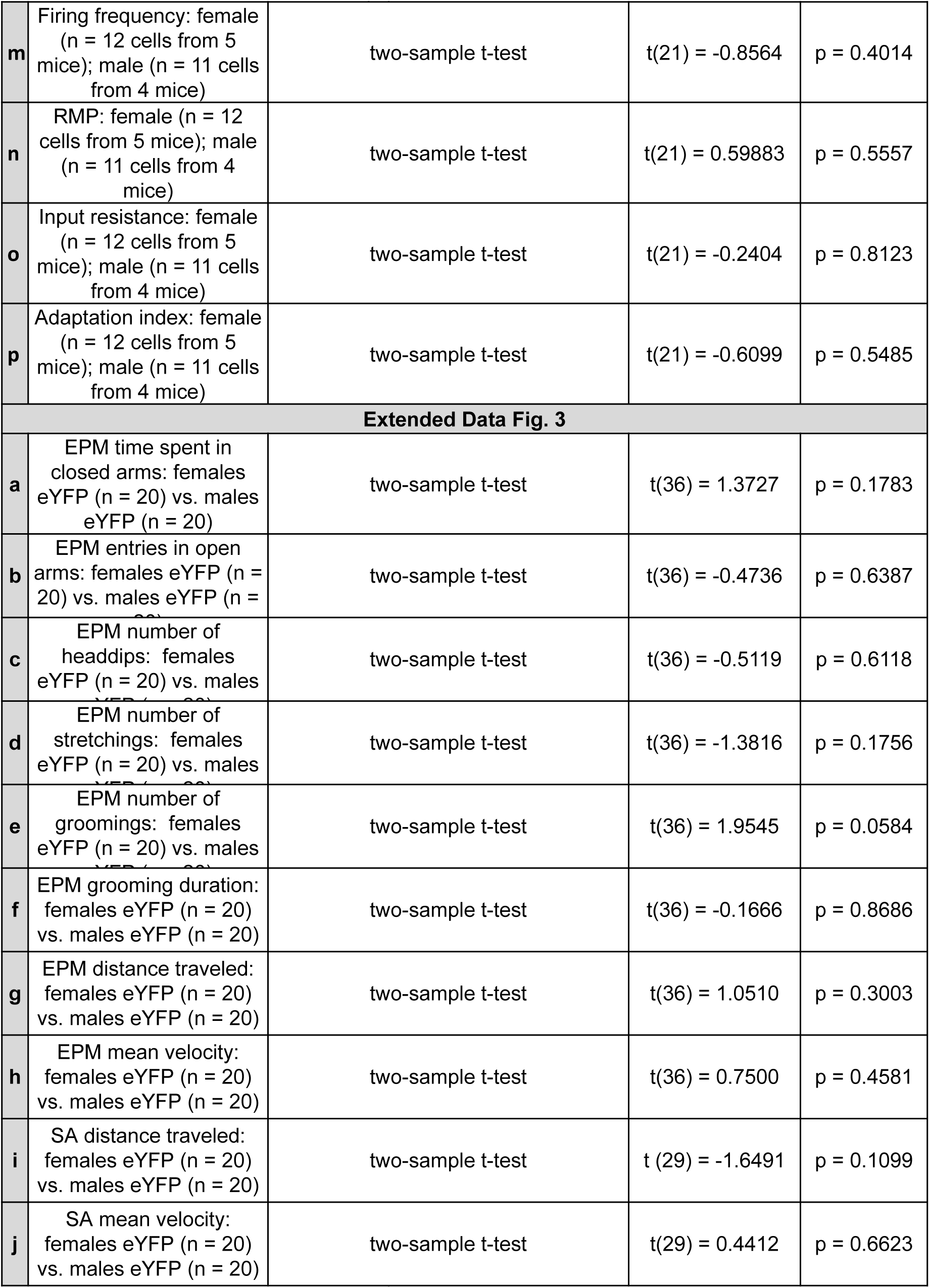
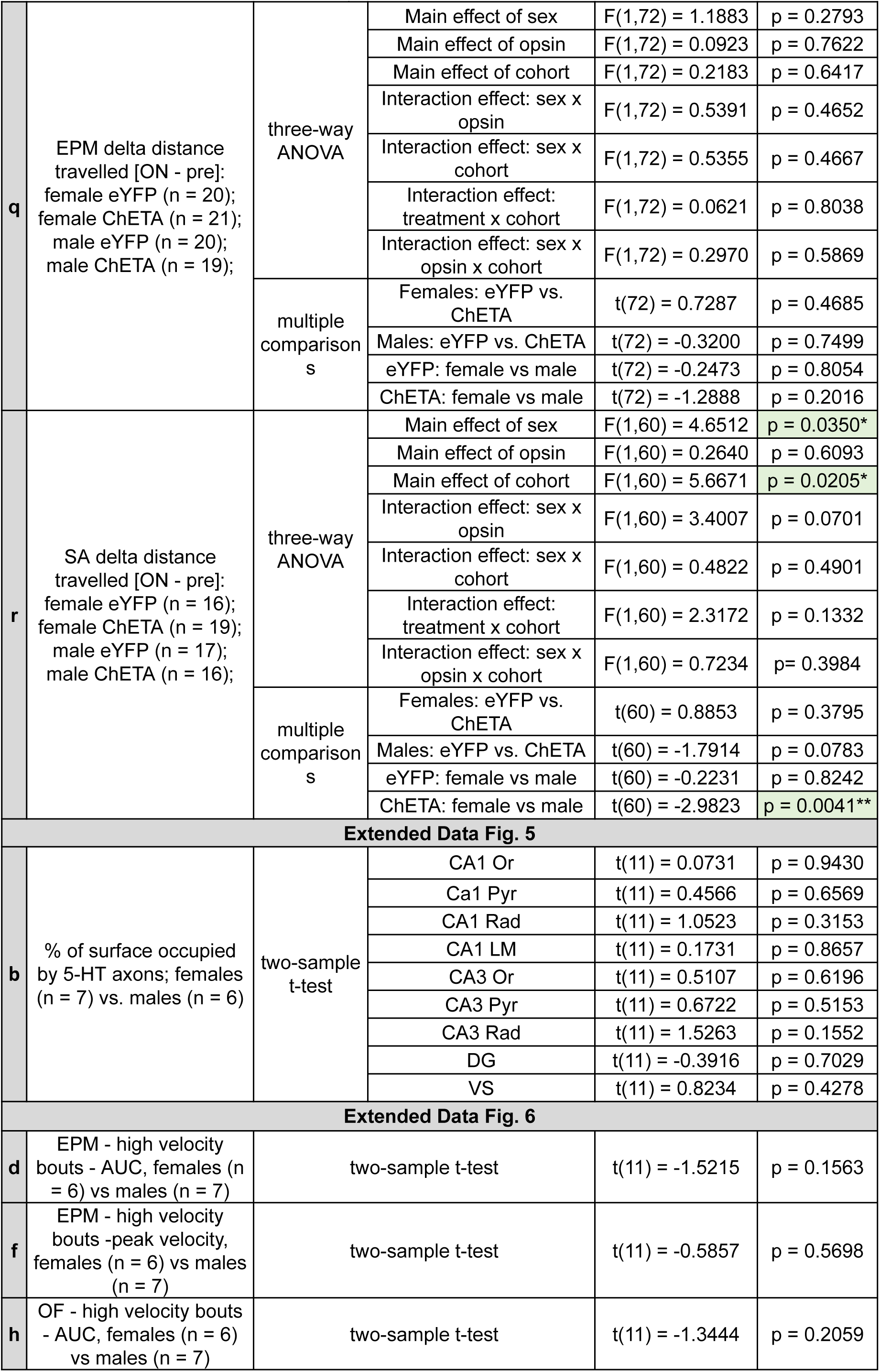
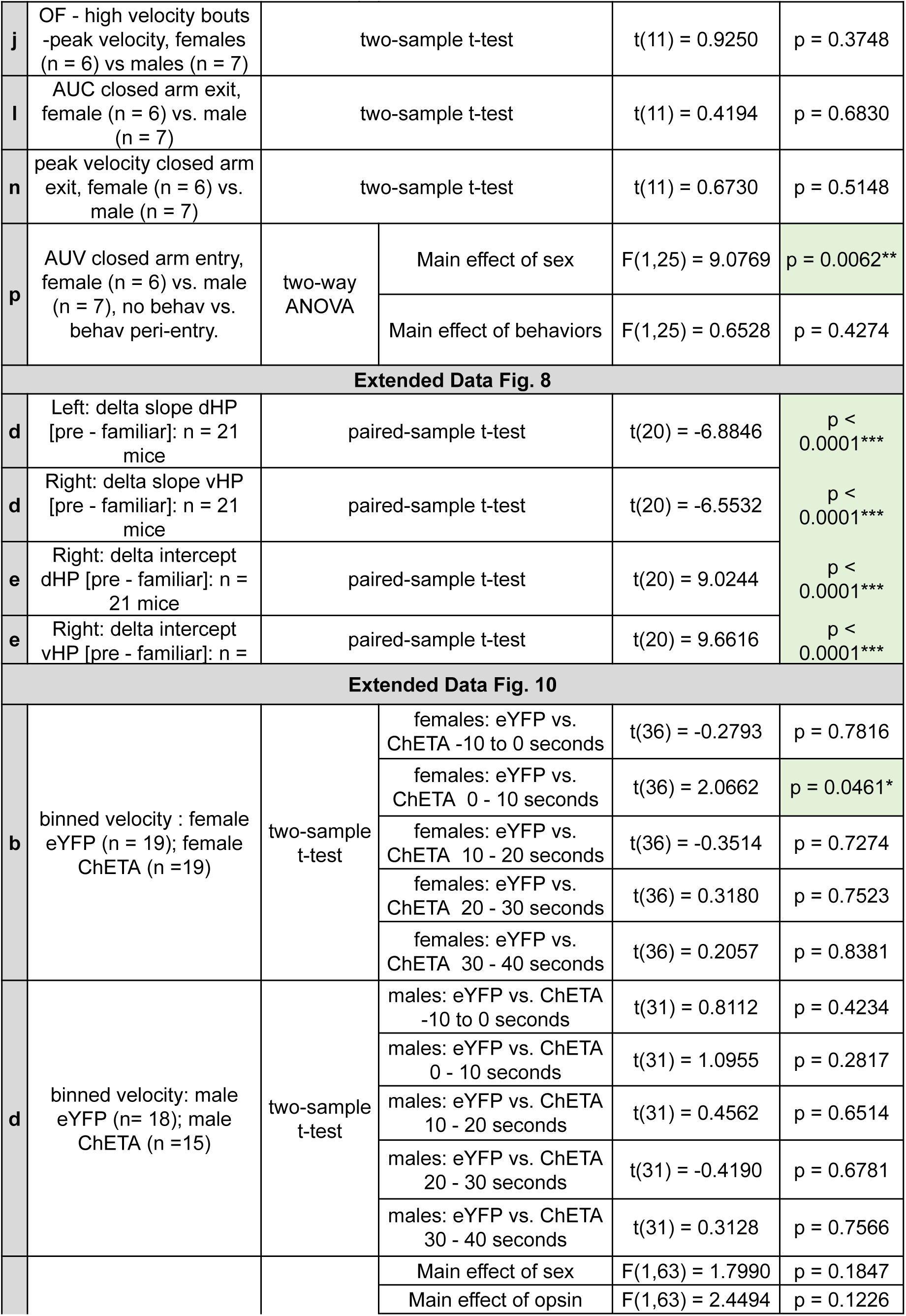
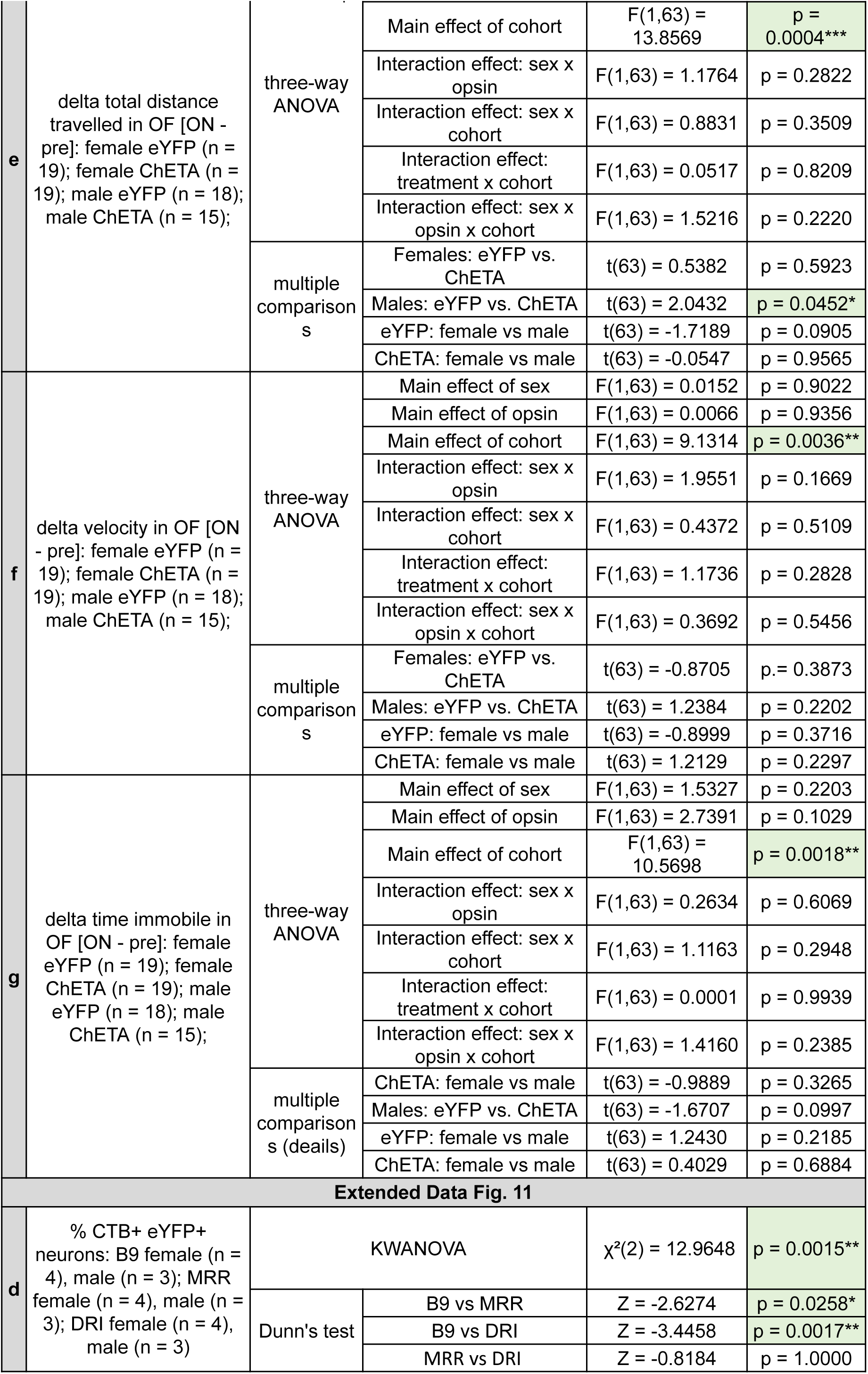
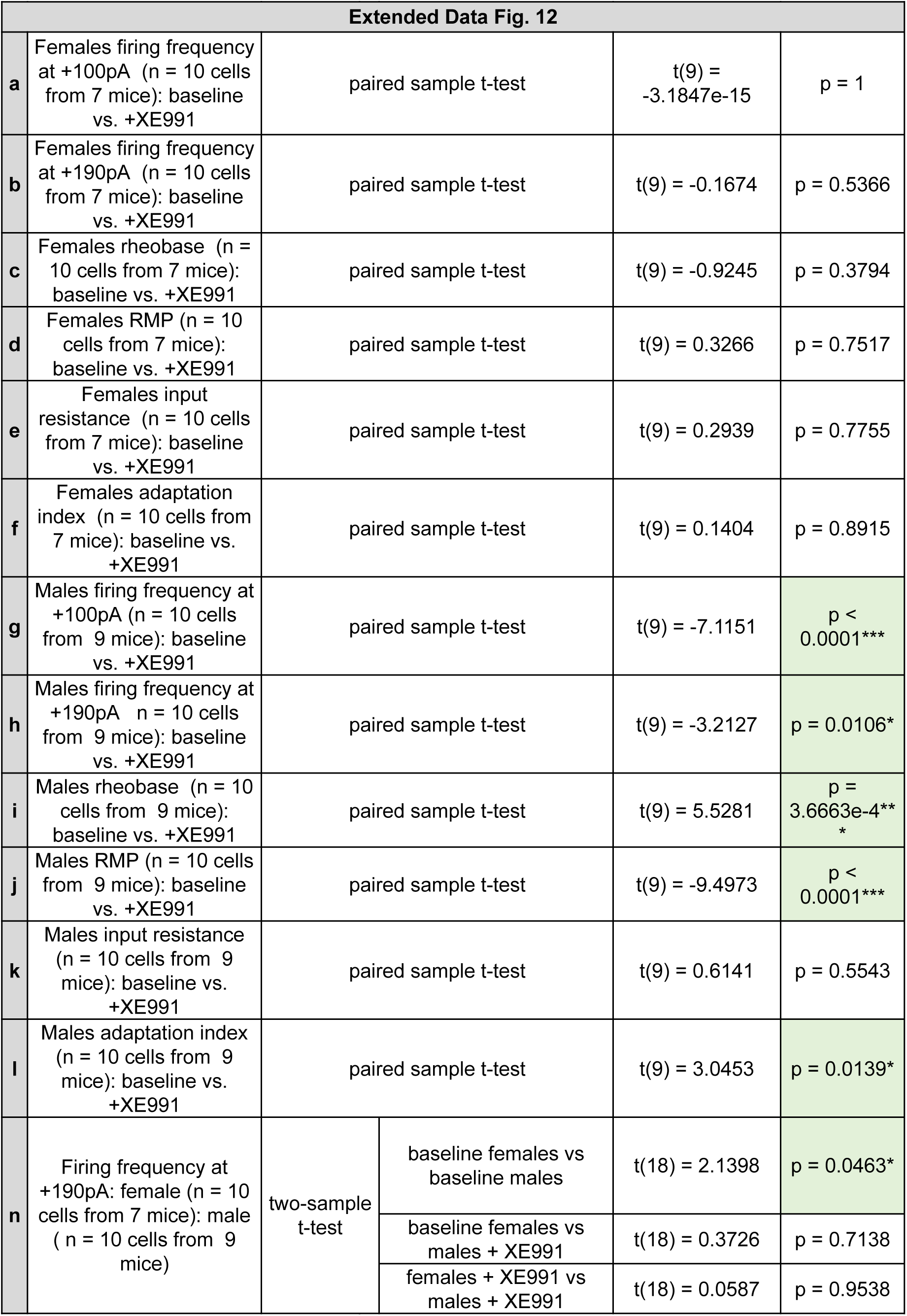
Summary of statistical analyses.

**Extended Data Table 2.**
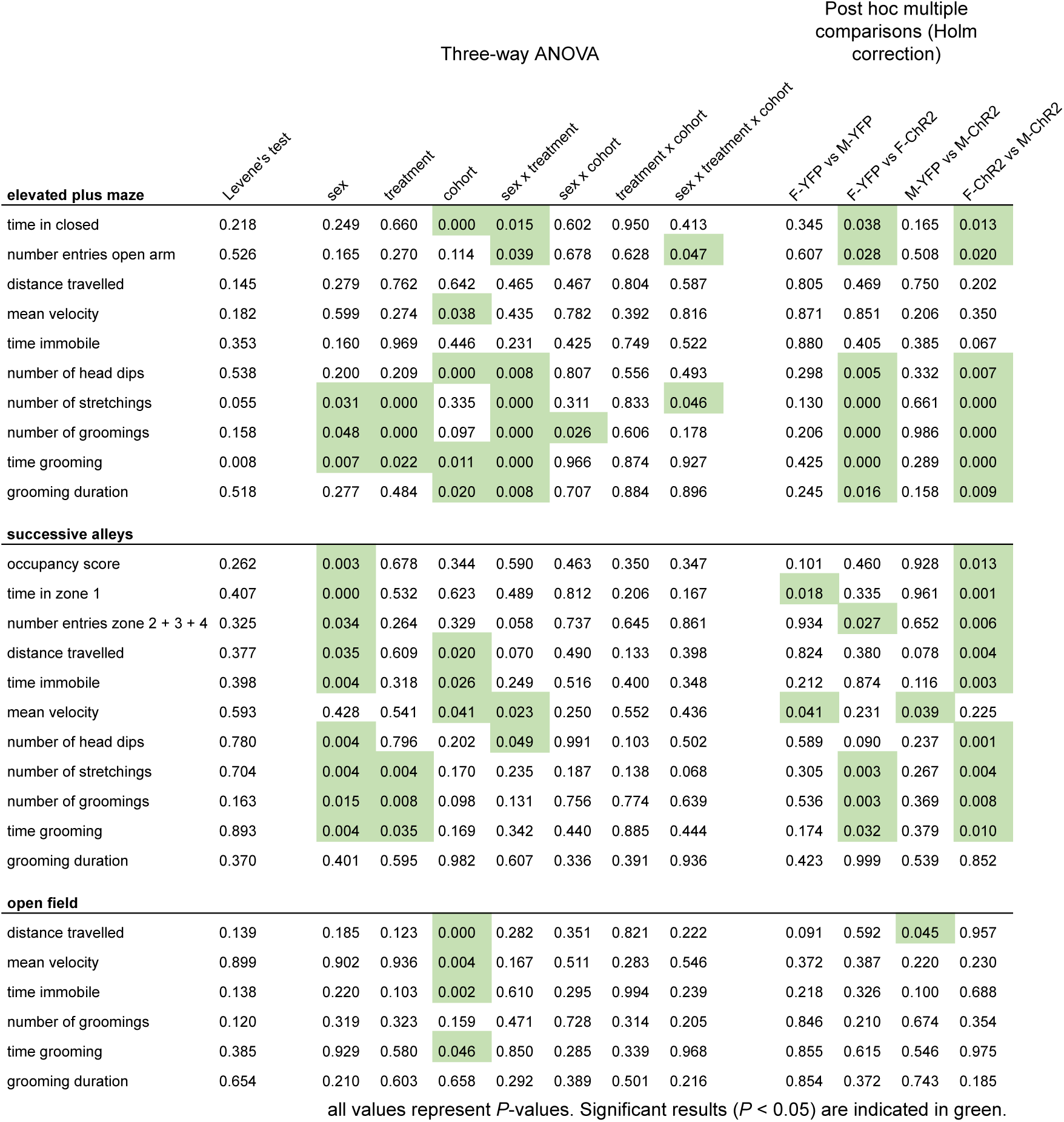
Comprehensive summary of all statistical testing for behavioral experiments detailed in Fig. 2, Fig. 3, Extended Data Fig. 2, Extended Data Fig. 7.

## References

1. Bangasser, D. A. & Cuarenta, A. Sex differences in anxiety and depression: circuits and mechanisms. Nat Rev Neurosci 22, 674–684 (2021).

2. McLean, C. P., Asnaani, A., Litz, B. T. & Hofmann, S. G. Gender differences in anxiety disorders: Prevalence, course of illness, comorbidity and burden of illness. Journal of Psychiatric Research 45, 1027–1035 (2011).

3. Graeff, F. G. & Zangrossi, H. The dual role of serotonin in defense and the mode of action of antidepressants on generalized anxiety and panic disorders. Cent Nerv Syst Agents Med Chem 10, 207–217 (2010).

4. Ren, J. et al. Anatomically defined and functionally distinct dorsal raphe serotonin sub-systems. Cell 175, 472–487.e20 (2018).

5. Marcinkiewcz, C. A. et al. Serotonin engages an anxiety and fear-promoting circuit in the extended amygdala. Nature 537, 97–101 (2016).

6. Urban, D. J. et al. Elucidation of the behavioral program and neuronal network encoded by dorsal raphe serotonergic neurons. Neuropsychopharmacology 41, 1404–1415 (2016).

7. Teissier, A. et al. Activity of raphé serotonergic neurons controls emotional behaviors. Cell Reports 13, 1965–1976 (2015).

8. Ohmura, Y. et al. Different roles of distinct serotonergic pathways in anxiety-like behavior, antidepressant-like, and anti-impulsive effects. Neuropharmacology 167, 107703 (2020).

9. Okaty, B. W., Commons, K. G. & Dymecki, S. M. Embracing diversity in the 5-HT neuronal system. Nat Rev Neurosci 20, 397–424 (2019).

10. Gaspar, P. & Lillesaar, C. Probing the diversity of serotonin neurons. Philosophical Transactions of the Royal Society B: Biological Sciences 367, 2382–2394 (2012).

11. Hale, M. W. & Lowry, C. A. Functional topography of midbrain and pontine serotonergic systems: implications for synaptic regulation of serotonergic circuits. Psychopharmacology 213, 243–264 (2011).

12. Awasthi, J. R., Tamada, K., Overton, E. T. N. & Takumi, T. Comprehensive topographical map of the serotonergic fibers in the male mouse brain. Journal of Comparative Neurology 529, 1391–1429 (2021).

13. Muzerelle, A., Scotto-Lomassese, S., Bernard, J. F., Soiza-Reilly, M. & Gaspar, P. Conditional anterograde tracing reveals distinct targeting of individual serotonin cell groups (B5–B9) to the forebrain and brainstem. Brain Struct Funct 221, 535–561 (2016).

14. Gagnon, D. & Parent, M. Distribution of VGLUT3 in highly collateralized axons from the rat dorsal raphe nucleus as revealed by single-neuron reconstructions. PLoS One 9, e87709 (2014).

15. Ren, J. et al. Single-cell transcriptomes and whole-brain projections of serotonin neurons in the mouse dorsal and median raphe nuclei. eLife 8, e49424 (2019).

16. Sharp, T. & Barnes, N. M. Central 5-HT receptors and their function; present and future. Neuropharmacology 177, 108155 (2020).

17. 17. Vilaró, M. T., Cortés, R., Mengod, G. & Hoyer, D. Distribution of 5-HT receptors in the central nervous system: an update. in Handbook of the Behavioral Neurobiology of Serotonin (eds Müller, C. P. & Cunnigham, K. A.) vol. 31 (Academic Press, 2020).

18. Yu, X.-D. et al. Distinct serotonergic pathways to the amygdala underlie separate behavioral features of anxiety. Nat Neurosci 25, 1651–1663 (2022).

19. Béïque, J.-C. et al. Serotonergic regulation of membrane potential in developing rat prefrontal cortex: coordinated expression of 5-hydroxytryptamine (5-HT)1A, 5-HT2A, and 5-HT7 receptors. J Neurosci **24**, 4807–4817 (2004).

20. Teixeira, C. M. et al. Hippocampal 5-HT input regulates memory formation and schaffer collateral excitation. Neuron 98, 992–1004.e4 (2018).

21. Kjelstrup, K. G. et al. Reduced fear expression after lesions of the ventral hippocampus. Proc. Natl. Acad. Sci. U.S.A. 99, 10825–10830 (2002).

22. Strange, B. A., Witter, M. P., Lein, E. S. & Moser, E. I. Functional organization of the hippocampal longitudinal axis. Nat Rev Neurosci 15, 655–669 (2014).

23. Bannerman, D. M. et al. Regional dissociations within the hippocampus--memory and anxiety. Neurosci Biobehav Rev 28, 273–283 (2004).

24. McNaughton, N. & Gray, J. A. Anxiolytic action on the behavioural inhibition system implies multiple types of arousal contribute to anxiety. J Affect Disord 61, 161–176 (2000).

25. Jimenez, J. C. et al. Anxiety cells in a hippocampal-hypothalamic circuit. Neuron 97, 670–683.e6 (2018).

26. Parfitt, G. M. et al. Bidirectional control of anxiety-related behaviors in mice: role of inputs arising from the ventral hippocampus to the lateral septum and medial prefrontal cortex. Neuropsychopharmacology 42, 1715–1728 (2017).

27. Ciocchi, S., Passecker, J., Malagon-Vina, H., Mikus, N. & Klausberger, T. Selective information routing by ventral hippocampal CA1 projection neurons. Science 348, 560–563 (2015).

28. Adhikari, A., Topiwala, M. A. & Gordon, J. A. Synchronized activity between the ventral hippocampus and the medial prefrontal cortex during anxiety. Neuron 65, 257–269 (2010).

29. Padilla-Coreano, N. et al. Direct ventral hippocampal-prefrontal input is required for anxiety-related neural activity and behavior. Neuron 89, 857–866 (2016).

30. Royer, S., Sirota, A., Patel, J. & Buzsáki, G. Distinct representations and theta dynamics in dorsal and ventral hippocampus. J Neurosci 30, 1777–1787 (2010).

31. Li, K., Koukoutselos, K., Sakaguchi, M. & Ciocchi, S. Distinct ventral hippocampal inhibitory microcircuits regulating anxiety and fear behaviors. Nat Commun 15, 8228 (2024).

32. Cerquetella, C., Gontier, C., Forro, T., Pfister, J.-P. & Ciocchi, S. Scaling of ventral hippocampal activity during anxiety. J Neurosci 45, e1128242025 (2025).

33. Sánchez-Bellot, C., AlSubaie, R., Mishchanchuk, K., Wee, R. W. S. & MacAskill, A. F. Two opposing hippocampus to prefrontal cortex pathways for the control of approach and avoidance behaviour. Nat Commun 13, 339 (2022).

34. Yeates, D. C. M. et al. Parallel ventral hippocampus-lateral septum pathways differentially regulate approach-avoidance conflict. Nat Commun 13, 3349 (2022).

35. Badarnee, M., Wen, Z., Nassar, N. & Milad, M. R. Gray matter associations with extinction-induced neural activation in patients with anxiety disorders. Journal of Psychiatric Research 162, 180–186 (2023).

36. Bach, D. R. et al. Human hippocampus arbitrates approach-avoidance conflict. Curr Biol 24, 541–547 (2014).

37. Ito, R. & Lee, A. C. H. The role of the hippocampus in approach-avoidance conflict decision-making: Evidence from rodent and human studies. Behavioural Brain Research 313, 345–357 (2016).

38. Cheng, H. et al. Projection-defined median raphe Pet+ subpopulations are diversely implicated in seizure. Neurobiology of Disease 189, 106358 (2023).

39. Fernandez, S. P. et al. Multiscale single-cell analysis reveals unique phenotypes of raphe 5-HT neurons projecting to the forebrain. Brain Struct Funct 221, 4007–4025 (2016).

40. Paquelet, G. E. et al. Single-cell activity and network properties of dorsal raphe nucleus serotonin neurons during emotionally salient behaviors. Neuron 110, 2664–2679.e8 (2022).

41. Sengupta, A. & Holmes, A. A discrete dorsal raphe to basal amygdala 5-HT circuit calibrates aversive memory. Neuron 103, 489–505.e7 (2019).

42. Harkin, E. F., Grossman, C. D., Cohen, J. Y., Béïque, J.-C. & Naud, R. A prospective code for value in the serotonin system. Nature 641, 952–959 (2025).

43. Dayan, P. & Huys, Q. J. M. Serotonin in affective control. Annual Review of Neuroscience 32, 95–126 (2009).

44. Andrade, T. G. & Graeff, F. G. Effect of electrolytic and neurotoxic lesions of the median raphe nucleus on anxiety and stress. Pharmacol Biochem Behav 70, 1–14 (2001).

45. De Almeida, R. M., Giovenardi, M., Charchat, H. & Lucion, A. B. 8-OH-DPAT in the median raphe nucleus decreases while in the medial septal area it may increase anxiety in female rats. Neurosci Biobehav Rev 23, 259–264 (1998).

46. Ahmadlou, M. et al. A subcortical switchboard for perseverative, exploratory and disengaged states. Nature 641, 151–161 (2025).

47. Kawai, H. et al. Median raphe serotonergic neurons projecting to the interpeduncular nucleus control preference and aversion. Nat Commun 13, 7708 (2022).

48. Maddaloni, G., Chang, Y. J., Senft, R. A. & Dymecki, S. M. Adaptation to photoperiod via dynamic neurotransmitter segregation. Nature 632, 147–156 (2024).

49. Huang, X. et al. A circuit from lateral hypothalamic to dorsal hippocampal dentate gyrus modulates behavioral despair in mice. Cerebral Cortex 34, bhae399 (2024).

50. Yoshida, K., Drew, M. R., Mimura, M. & Tanaka, K. F. Serotonin-mediated inhibition of ventral hippocampus is required for sustained goal-directed behavior. Nat Neurosci 22, 770–777 (2019).

51. Abela, A. R. et al. Median raphe serotonin neurons promote anxiety-like behavior via inputs to the dorsal hippocampus. Neuropharmacology 168, 107985 (2020).

52. Freund, T. F., Gulyás, A. I., Acsády, L., Görcs, T. & Tóth, K. Serotonergic control of the hippocampus via local inhibitory interneurons. Proceedings of the National Academy of Sciences 87, 8501–8505 (1990).

53. Varga, V. et al. Fast synaptic subcortical control of hippocampal circuits. Science 326, 449–453 (2009).

54. Turi, G. F. et al. Serotonin modulates infraslow oscillation in the dentate gyrus during Non-REM sleep. eLife 13, (2025).

55. Maru, E., Takahashi, L. K. & Iwahara, S. Effects of median raphe nucleus lesions on hippocampal EEG in the freely moving rat. Brain Research 163, 223–234 (1979).

56. Yamamoto, T., Watanabe, S., Oishi, R. & Ueki, S. Effects of midbrain raphe stimulation and lesion on EEG activity in rats. Brain Research Bulletin 4, 491–495 (1979).

57. Fernandez, S. P. et al. Constitutive and Acquired Serotonin Deficiency Alters Memory and Hippocampal Synaptic Plasticity. Neuropsychopharmacol 42, 512–523 (2017).

58. Ohmura, Y. et al. Serotonin 5-ht 7 receptor in the ventral hippocampus modulates the retrieval of fear memory and stress-induced defecation. Int J Neuropsychopharmacol 10.1093/ijnp/pyv131 (2015) doi:10.1093/ijnp/pyv131.

59. Karayol, R. et al. Serotonin receptor 4 in the hippocampus modulates mood and anxiety. Mol Psychiatry 26, 2334–2349 (2021).

60. Alves, S. H., Pinheiro, G., Motta, V., Landeira-Fernandez, J. & Cruz, A. P. M. Anxiogenic effects in the rat elevated plus-maze of 5-HT(2C) agonists into ventral but not dorsal hippocampus. Behav Pharmacol 15, 37–43 (2004).

61. Howerton, A. R. et al. Sex differences in corticotropin-releasing factor receptor-1 action within the dorsal raphe nucleus in stress responsivity. Biol Psychiatry 75, 873–883 (2014).

62. Torres Irizarry, V. C., et al. Estrogen signaling in the dorsal raphe regulates binge-like drinking in mice. Transl Psychiatry 14, 122 (2024).

63. Beck, S. G., Pan, Y.-Z., Akanwa, A. C. & Kirby, L. G. Median and dorsal raphe neurons are not electrophysiologically identical. J Neurophysiol 91, 994–1005 (2004).

64. Dominguez, R., Cruz-Morales, S. E., Carvalho, M. C., Xavier, M. & Brandao, M. L. Sex differences in serotonergic activity in dorsal and median raphe nucleus. Physiology & Behavior 80, 203–210 (2003).

65. Ohmura, Y., Tanaka, K. F., Tsunematsu, T., Yamanaka, A. & Yoshioka, M. Optogenetic activation of serotonergic neurons enhances anxiety-like behaviour in mice. International Journal of Neuropsychopharmacology 17, 1777–1783 (2014).

66. Konno, K. et al. Early postnatal stress affects the serotonergic function in the median raphe nuclei of adult rats. Brain Res 1172, 60–66 (2007).

67. Palanza, P. Animal models of anxiety and depression: how are females different? Neurosci Biobehav Rev 25, 219–233 (2001).

68. Tsao, C.-H., Wu, K.-Y., Su, N. C., Edwards, A. & Huang, G.-J. The influence of sex difference on behavior and adult hippocampal neurogenesis in C57BL/6 mice. Sci Rep 13, 17297 (2023).

69. Deacon, R. M. J. The successive alleys test of anxiety in mice and rats. J Vis Exp 2705 (2013) doi:10.3791/2705.

70. Kalueff, A. V. & Tuohimaa, P. The grooming analysis algorithm discriminates between different levels of anxiety in rats: potential utility for neurobehavioural stress research. Journal of Neuroscience Methods 143, 169–177 (2005).

71. Spruijt, B. M., van Hooff, J. A. & Gispen, W. H. Ethology and neurobiology of grooming behavior. Physiological Reviews 72, 825–852 (1992).

72. Kalueff, A. V. et al. Neurobiology of rodent self-grooming and its value for translational neuroscience. Nat Rev Neurosci 17, 45–59 (2016).

73. Berridge, K. C. Comparative fine structure of action: Rules of form and sequence in the grooming patterns of six rodent species. Behaviour 113, 21–56 (1990).

74. Morishita, H. et al. Delayed response of the median raphe serotonin neurons projecting to the ventral hippocampus to aversive stimuli. J Pharmacol Sci 160, 91–96 (2026).

75. Metzger, M., Bueno, D. & Lima, L. B. The lateral habenula and the serotonergic system. Pharmacology Biochemistry and Behavior 162, 22–28 (2017).

76. Vertes, R. P., Fortin, W. J. & Crane, A. M. Projections of the median raphe nucleus in the rat. J Comp Neurol 407, 555–582 (1999).

77. Amo, R. et al. The habenulo-raphe serotonergic circuit encodes an aversive expectation value essential for adaptive active avoidance of danger. Neuron 84, 1034–1048 (2014).

78. Rex, A., Voigt, J. P. & Fink, H. Anxiety but not arousal increases 5-hydroxytryptamine release in the rat ventral hippocampus in vivo. European Journal of Neuroscience 22, 1185–1189 (2005).

79. Kusljic, S., Copolov, D. L. & van den Buuse, M. Differential role of serotonergic projections arising from the dorsal and median raphe nuclei in locomotor hyperactivity and prepulse inhibition. Neuropsychopharmacol 28, 2138–2147 (2003).

80. Wirtshafter, D. & Asin, K. E. Evidence that electrolytic median raphe lesions increase locomotion but not exploration. Physiology & Behavior 28, 749–754 (1982).

81. Casarrubea, M. et al. Temporal structure of the rat’s behavior in elevated plus maze test. Behavioural Brain Research 237, 290–299 (2013).

82. Arabo, A., Potier, C., Ollivier, G., Lorivel, T. & Roy, V. Temporal analysis of free exploration of an elevated plus-maze in mice. Journal of Experimental Psychology: Animal Learning and Cognition 40, 457–466 (2014).

83. Akiti, K. et al. Striatal dopamine explains novelty-induced behavioral dynamics and individual variability in threat prediction. Neuron 110, 3789–3804.e9 (2022).

84. Hasselmo, M. E. & Stern, C. E. Theta rhythm and the encoding and retrieval of space and time. NeuroImage 85, 656–666 (2014).

85. Colgin, L. L. Rhythms of the hippocampal network. Nat Rev Neurosci 17, 239–249 (2016).

86. Buzsáki, G. & Moser, E. I. Memory, navigation and theta rhythm in the hippocampal-entorhinal system. Nat Neurosci 16, 130–138 (2013).

87. Whishaw, I. Q. & Vanderwolf, C. H. Hippocampal EEG and behavior: changes in amplitude and frequency of RSA (theta rhythm) associated with spontaneous and learned movement patterns in rats and cats. Behav Biol 8, 461–484 (1973).

88. Hinman, J. R., Penley, S. C., Long, L. L., Escabí, M. A. & Chrobak, J. J. Septotemporal variation in dynamics of theta: speed and habituation. J Neurophysiol 105, 2675–2686 (2011).

89. Jeewajee, A., Lever, C., Burton, S., O’Keefe, J. & Burgess, N. Environmental novelty is signaled by reduction of the hippocampal theta frequency. Hippocampus 18, 340–348 (2008).

90. Kennedy, J. P. et al. A direct Comparison of theta Power and frequency to speed and acceleration. J Neurosci 42, 4326–4341 (2022).

91. Wells, C. E. et al. Novelty and anxiolytic drugs dissociate two components of hippocampal theta in behaving rats. J. Neurosci. 33, 8650–8667 (2013).

92. Monaghan, C. K., Chapman, G. W. & Hasselmo, M. E. Systemic administration of two different anxiolytic drugs decreases local field potential theta frequency in the medial entorhinal cortex without affecting grid cell firing fields. Neuroscience 364, 60–70 (2017).

93. Hines, M. et al. Frequency matters: how changes in hippocampal theta frequency can influence temporal coding, anxiety-reduction, and memory. Front. Syst. Neurosci. 16, 998116 (2023).

94. Correia, P. A. et al. Transient inhibition and long-term facilitation of locomotion by phasic optogenetic activation of serotonin neurons. Elife 6, e20975 (2017).

95. Seo, C. et al. Intense threat switches dorsal raphe serotonin neurons to a paradoxical operational mode. Science 363, 538–542 (2019).

96. Brown, D. A. & Passmore, G. M. Neural KCNQ (Kv7) channels. Br J Pharmacol 156, 1185–1195 (2009).

97. Marrion, N. V. Control of M-current. Annual Review of Physiology 59, 483–504 (1997).

98. Brown, D. A. & Adams, P. R. Muscarinic suppression of a novel voltage-sensitive K+ current in a vertebrate neurone. Nature 283, 673–676 (1980).

99. Huang, H. & Trussell, L. O. KCNQ5 channels control resting properties and release probability of a synapse. Nat Neurosci 14, 840–847 (2011).

100. Hansen, H. H. et al. Kv7 channels: interaction with dopaminergic and serotonergic neurotransmission in the CNS. J Physiol 586, 1823–1832 (2008).

101. Zhao, C. et al. Selective modulation of K+ channel Kv7.4 significantly affects the excitability of DRN 5-HT neurons. Front. Cell. Neurosci. 11, (2017).

102. Bayasgalan, T., Csemer, A., Kovacs, A., Pocsai, K. & Pal, B. Topographical organization of M-current on dorsal and median raphe serotonergic neurons. Front. Cell. Neurosci. 15, (2021).

103. Wang, J., Wang, Y., Du, X. & Zhang, H. Potassium channel conductance is involved in phenylephrine-induced spontaneous firing of serotonergic neurons in the dorsal raphe nucleus. Front. Cell. Neurosci. 16, (2022).

104. Köhler, C. & Steinbusch, H. Identification of serotonin and non-serotonin-containing neurons of the mid-brain raphe projecting to the entorhinal area and the hippocampal formation. A combined immunohistochemical and fluorescent retrograde tracing study in the rat brain. Neuroscience 7, 951–975 (1982).

105. McKenna, J. T. & Vertes, R. P. Collateral projections from the median raphe nucleus to the medial septum and hippocampus. Brain Res Bull 54, 619–630 (2001).

106. Jacobs, B. L. & Azmitia, E. C. Structure and function of the brain serotonin system. Physiol Rev 72, 165–229 (1992).

107. Guajardo, H. M., Hatini, P. G. & Commons, K. G. The mouse dorsal raphe nucleus as understood by temporal Fgf8 lineage analysis. J Comp Neurol 529, 2042–2054 (2021).

108. Jensen, P. et al. Redefining the serotonergic system by genetic lineage. Nat Neurosci 11, 417–419 (2008).

109. Senft, R. A., Freret, M. E., Sturrock, N. & Dymecki, S. M. Neurochemically and hodologically distinct ascending VGLUT3 versus serotonin subsystems comprise the r2-Pet1 median raphe. Journal of Neuroscience 41, 2581–2600 (2021).

110. Amilhon, B. et al. VGLUT3 (Vesicular Glutamate Transporter Type 3) Contribution to the regulation of serotonergic transmission and anxiety. Journal of Neuroscience 30, 2198–2210 (2010).

111. Fortin-Houde, J., Henderson, F., Dumas, S., Ducharme, G. & Amilhon, B. Parallel streams of raphe VGLUT3-positive inputs target the dorsal and ventral hippocampus in each hemisphere. J Comp Neurol 531, 702–719 (2023).

112. Henderson, F. et al. Regulation of stress-induced sleep perturbations by dorsal raphe VGLUT3 neurons in male mice. Cell Rep 43, 114411 (2024).

113. Sengupta, A., Bocchio, M., Bannerman, D. M., Sharp, T. & Capogna, M. Control of amygdala circuits by 5-HT neurons via 5-HT and glutamate cotransmission. J Neurosci 37, 1785–1796 (2017).

114. Wang, H.-L. et al. Dorsal raphe dual serotonin-glutamate neurons drive reward by establishing excitatory synapses on VTA mesoaccumbens dopamine neurons. Cell Rep 26, 1128–1142.e7 (2019).

115. Cohen, J. Y., Amoroso, M. W. & Uchida, N. Serotonergic neurons signal reward and punishment on multiple timescales. Elife 4, e06346 (2015).

116. Matias, S., Lottem, E., Dugué, G. P. & Mainen, Z. F. Activity patterns of serotonin neurons underlying cognitive flexibility. Elife 6, e20552 (2017).

117. Zhong, W., Li, Y., Feng, Q. & Luo, M. Learning and stress shape the reward response patterns of serotonin neurons. J Neurosci 37, 8863–8875 (2017).

118. Sörman, E., Wang, D., Hajos, M. & Kocsis, B. Control of hippocampal theta rhythm by serotonin: Role of 5-HT2c receptors. Neuropharmacology 61, 489–494 (2011).

119. Olvera-Cortés, M. E., Gutiérrez-Guzmán, B. E., López-Loeza, E., Hernández-Pérez, J. J. & López-Vázquez, M. Á. Serotonergic modulation of hippocampal theta activity in relation to hippocampal information processing. Exp Brain Res 230, 407–426 (2013).

120. Vertes, R. P. & Linley, S. B. Chapter 19 - Serotonergic regulation of hippocampal rhythmical activity. in Handbook of Behavioral Neuroscience (eds Müller, C. P. & Cunningham, K. A.) vol. 31 337–360 (Elsevier, 2020).

121. Jackson, J., Dickson, C. T. & Bland, B. H. Median raphe stimulation disrupts hippocampal theta via rapid inhibition and state-dependent phase reset of theta-related neural circuitry. J Neurophysiol 99, 3009–3026 (2008).

122. Lee, A. T. et al. VIP interneurons contribute to avoidance behavior by regulating information flow across hippocampal-prefrontal networks. Neuron 102, 1223–1234.e4 (2019).

123. Song, C., Berridge, K. C. & Kalueff, A. V. ‘Stressing’ rodent self-grooming for neuroscience research. Nat Rev Neurosci 17, 591–591 (2016).

124. Fernández-Teruel, A. & Estanislau, C. Meanings of self-grooming depend on an inverted U-shaped function with aversiveness. Nat Rev Neurosci 17, 591–591 (2016).

125. Tanaka, K. F., Samuels, B. A. & Hen, R. Serotonin receptor expression along the dorsal–ventral axis of mouse hippocampus. Philos Trans R Soc Lond B Biol Sci 367, 2395–2401 (2012).

126. Jin, J. & Maren, S. Fear renewal preferentially activates ventral hippocampal neurons projecting to both amygdala and prefrontal cortex in rats. Sci Rep 5, 8388 (2015).

127. Jimenez, J. C. et al. Contextual fear memory retrieval by correlated ensembles of ventral CA1 neurons. Nat Commun 11, 3492 (2020).

128. Gergues, M. M. et al. Circuit and molecular architecture of a ventral hippocampal network. Nat Neurosci 23, 1444–1452 (2020).

129. Wee, R. W. S. & MacAskill, A. F. Biased connectivity of brain-wide inputs to ventral subiculum output neurons. Cell Rep 30, 3644–3654.e6 (2020).

130. Paul, E. D. & Lowry, C. A. Functional topography of serotonergic systems supports the Deakin/Graeff hypothesis of anxiety and affective disorders. J Psychopharmacol 27, 1090–1106 (2013).

131. Ravenelle, R. et al. Serotonergic modulation of the BNST-CeA pathway reveals sex differences in fear learning. Nat Neurosci 10.1038/s41593-025-02025-x (2025) doi:10.1038/s41593-025-02025-x.

132. Craske, M. G. & Stein, M. B. Anxiety. Lancet 388, 3048–3059 (2016).

133. Kessler, R. C. et al. The global burden of mental disorders: An update from the WHO World Mental Health (WMH) Surveys. Epidemiol Psichiatr Soc 18, 23–33 (2009).

134. Kessler, R. C. et al. Lifetime prevalence and age-of-onset distributions of DSM-IV disorders in the National Comorbidity Survey Replication. Archives of General Psychiatry 62, 593–602 (2005).

135. Keevers, L. J. & Jean-Richard-dit-Bressel, P. Obtaining artifact-corrected signals in fiber photometry via isosbestic signals, robust regression, and dF/F calculations. Neurophotonics 12, 025003 (2025).

136. Pereira, T. D. et al. SLEAP: A deep learning system for multi-animal pose tracking. Nat Methods 19, 486–495 (2022).

137. Siegle, J. H. et al. Open Ephys: an open-source, plugin-based platform for multichannel electrophysiology. J Neural Eng 14, 045003 (2017).

138. Mathis, A. et al. DeepLabCut: markerless pose estimation of user-defined body parts with deep learning. Nature Neuroscience 21, 1281–1289 (2018).

139. Nath, T. et al. Using DeepLabCut for 3D markerless pose estimation across species and behaviors. Nat Protoc 14, 2152–2176 (2019).

140. Paxinos, G. & Franklin, K. B. J. Paxinos and Franklin’s the Mouse Brain in Stereotaxic Coordinates. (Academic Press, 2019).

141. Ting, J. T., Daigle, T. L., Chen, Q. & Feng, G. Acute brain slice methods for adult and aging animals: application of targeted patch clamp analysis and optogenetics. Methods Mol Biol 1183, 221–242 (2014).

142. Nigro, M. J., Mateos-Aparicio, P. & Storm, J. F. Expression and functional roles of Kv7/KCNQ/M-channels in rat medial entorhinal cortex layer II stellate cells. J Neurosci 34, 6807–6812 (2014).

143. Bordas, C., Kovacs, A. & Pal, B. The M-current contributes to high threshold membrane potential oscillations in a cell type-specific way in the pedunculopontine nucleus of mice. Front. Cell. Neurosci. 9, (2015).

144. Wang, H. S., Brown, B. S., McKinnon, D. & Cohen, I. S. Molecular basis for differential sensitivity of KCNQ and I(Ks) channels to the cognitive enhancer XE991. Mol Pharmacol 57, 1218–1223 (2000).

145. Zhang, X. et al. Selective activation of vascular Kv7.4/Kv7.5 K+ channels by fasudil contributes to its vasorelaxant effect. Br J Pharmacol 173, 3480–3491 (2016).

146. Brown, M. B. & Forsythe, A. B. Robust tests for the equality of variances. Journal of the American Statistical Association 69, 364–367 (1974).

